# Ripple-locked coactivity of stimulus-specific neurons supports human associative memory

**DOI:** 10.1101/2022.10.17.512635

**Authors:** Lukas Kunz, Bernhard P. Staresina, Peter C. Reinacher, Armin Brandt, Tim A. Guth, Andreas Schulze-Bonhage, Joshua Jacobs

**Affiliations:** Department of Biomedical Engineering, Columbia University, New York, NY, USA; Department of Experimental Psychology, University of Oxford, Oxford, UK; Oxford Centre for Human Brain Activity, Wellcome Centre for Integrative Neuroimaging, Department of Psychiatry, University of Oxford, Oxford, UK; Department of Stereotactic and Functional Neurosurgery, Medical Center – University of Freiburg, Faculty of Medicine, University of Freiburg, Freiburg, Germany; Fraunhofer Institute for Laser Technology, Aachen, Germany; Epilepsy Center, Medical Center – University of Freiburg, Faculty of Medicine, University of Freiburg, Freiburg, Germany

## Abstract

Associative memory is the ability to encode and retrieve relations between different stimuli. To better understand its neural basis, we investigated whether associative memory involves precisely timed spiking of neurons in the medial temporal lobes that exhibit stimulus-specific tuning. Using single-neuron recordings from epilepsy patients performing an associative object–location memory task, we identified the object- and place-specific neurons that encoded the separate elements of each memory. When patients encoded and retrieved particular memories, the relevant object- and place-specific neurons activated synchronously during hippocampal ripples. This ripple-locked coactivity of stimulus-specific neurons emerged over time as the patients’ associative learning progressed. Our results suggest a cellular account of associative memory, in which hippocampal ripples coordinate the activity of specialized cellular populations to facilitate links between stimuli.

## Introduction

Associative memory is an essential cognitive function for everyday life that allows us to learn and remember relations between different stimuli (*1*). Impairments in associative memory caused by aging and memory disorders (*2, 3*) are thus a growing problem for society, which makes it important to better understand its neural basis. A large body of research has implicated the hippocampus and neighboring medial temporal lobe (MTL) regions in the encoding and retrieval of associative memories (*4, 5*). Here, we sought to further elucidate the mechanisms underlying associative memory in the human MTL at the single-cell level. Given prior theories on temporally precise neural binding in perception and memory (*6–8*), we considered that the individual stimuli contributing to particular associative memories are encoded by separate sets of stimulus-specific neurons and that these neurons interact transiently when subjects encode and retrieve the memories (Fig. 1A).

**Fig. 1.**
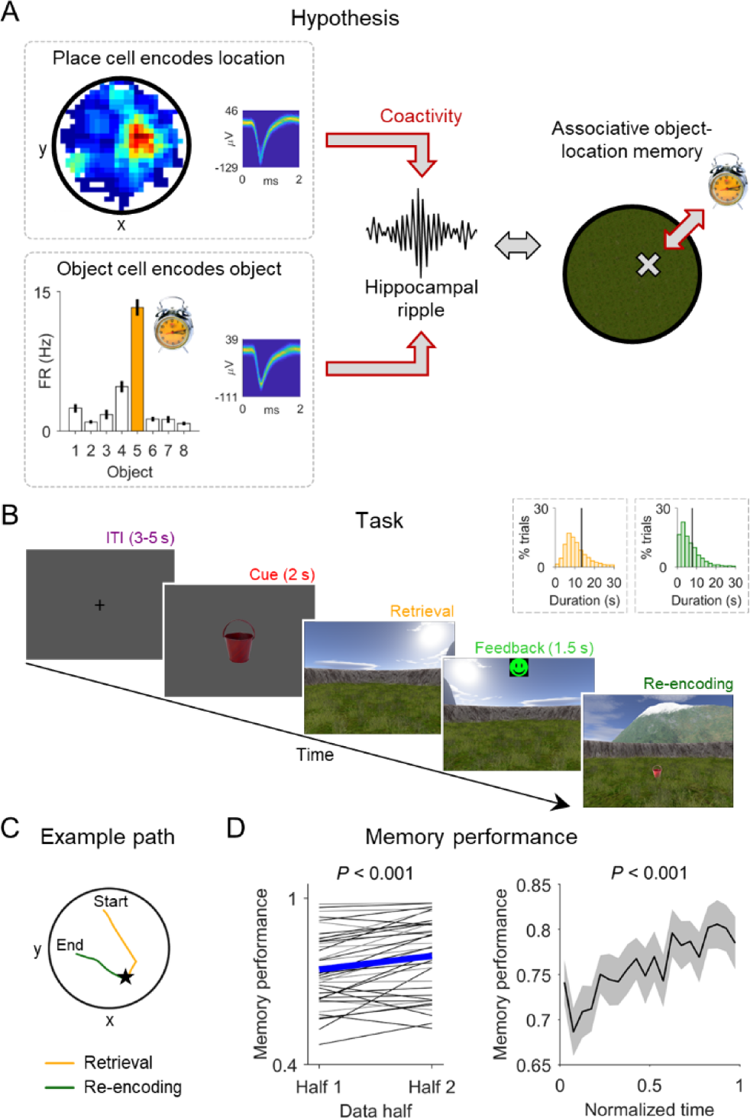
Hypothesis and associative object–location memory task. (**A**) Illustration of the hypothesis that human associative object–location memory is linked to the coactivity of object cells and place cells during hippocampal ripples. We propose that the coactivity is specific to pairs of object and place cells that encode “associative information,” which are those cell pairs where the location of the preferred object of the object cell is inside the place field of the place cell. (**B**) Subjects performed an associative object–location memory task while navigating freely in a virtual environment. After collecting eight different objects from their associated locations during an initial encoding period (not shown), subjects performed a series of test trials. At the beginning of each test trial, after an inter-trial interval (“ITI”), one of the eight objects was presented (“Cue”), which the subject placed as accurately as possible at its associated location during retrieval (“Retrieval”). Subjects received feedback depending on the accuracy of their response (“Feedback”) and collected the then visible object from its correct location (“Re-encoding”). Insets show histograms of the self-paced durations of retrieval (yellow) and re-encoding (green) periods. (**C**) Example paths during retrieval (yellow) and re-encoding (green) in one trial. The subject’s response location is indicated by a star. (**D**) Left: memory performance during the first versus the second data half showing that subjects improved their memories over time. Blue thick line, mean across sessions; thin lines, session-wise data (black, sessions with single-neuron recordings). Right: memory performance as a function of normalized time. Black, mean across sessions; gray, SEM across sessions. FR, firing rate.

We examined the neural basis of associative memory in the setting of object–location associations. We hypothesized that the encoding and retrieval of such object–location memories is supported by the simultaneous activation of object cells, which represent specific objects (*9, 10*), and place cells, which code for particular spatial locations (*11, 12*). We specifically hypothesized that these coactivations would occur in a temporally precise manner during hippocampal ripples, which are bouts of high-frequency oscillations (*13–15*). Such ripple-locked coactivity of object and place cells could underlie the encoding and retrieval of associative object–location memories by inducing and activating synaptic connections between the object and place cells that represent the different memory elements (*16*). In addition, ripple-locked coactivity of stimulus-specific neurons could elicit conjunctive memory representations in downstream neurons that only respond to the unique combination of all memory elements (*17–20*).

We reasoned that hippocampal ripples might coordinate the coactivity of object and place cells because they synchronize neural activity across close and distant brain regions (*13, 14, 21–24*). Hippocampal ripples may thus help induce and activate large-scale cellular networks (*16*), which we considered instrumental in linking and coordinating otherwise unconnected neurons for the encoding and retrieval of associative memories. Prior studies in rodents showed that ripples are linked to precisely organized multicellular activity (*25–30*) in the service of learning, memory, and planning (*13, 14, 31, 32*). Similarly, neural recordings in epilepsy patients indicated that ripples correlate with various cognitive functions in humans, including memory encoding, retrieval, and consolidation (*33*– *42*). How human ripples coordinate the activity of stimulus-specific single neurons to support associative memory has remained elusive, however.

We thus examined the hypothesis that human hippocampal ripples support the formation and retrieval of associative memories by defining time windows for the coactivity of stimulus-specific neurons. To test this idea, we conducted single-neuron and intracranial electroencephalographic (EEG) recordings from the MTL of human epilepsy patients who performed an associative object–location memory task in a virtual environment (*9*). In line with our hypothesis, we show that object- and place-specific neurons that represent the separate elements of particular object–location associations activate together at the moments of hippocampal ripples. Our work therefore suggests that ripple-locked coactivity of stimulus-specific neurons constitutes a neural mechanism for the formation and retrieval of associative memories in the human brain.

## Results

### Ripples in the human hippocampus during an associative object–location memory task

To study the neural mechanisms underlying human associative memory, we recorded single-neuron activity and intracranial EEG from the MTL of epilepsy patients undergoing presurgical monitoring (Materials and Methods; Table S1) (*43, 44*). During the recordings, subjects performed an associative object–location memory task in a virtual environment (Fig. 1, B–C). In this task (*9*), subjects encoded the locations of eight different objects once during an initial encoding period and then performed a series of test trials that included periods for retrieving and re-encoding the object–location associations. Briefly, each test trial started with an inter-trial interval (ITI), followed by a cue period where the subject viewed one of the eight objects that they had encountered during the initial encoding period. Then, in the retrieval phase, subjects navigated to the remembered location of this object and received feedback depending on their response accuracy. After feedback, in the re-encoding phase, the object appeared in its correct location and subjects traveled to this location, allowing them to refine their associative memory for this object–location pair. Thirty subjects contributed a total of 41 sessions and performed 103 trials per session on average (for detailed information on all statistics in the main text, see Table S2). They successfully formed associative memories between the eight objects and their corresponding locations, as their memory performance increased over the course of the task [paired *t*-test: *t*(40) = −4.788, *P* < 0.001; Fig. 1D].

We identified human hippocampal ripples during the task by examining local field potentials (LFPs) from bipolar macroelectrode channels, which were mostly located in the CA1 region of the anterior hippocampus (Fig. 2, A–D; Fig. S1). Following previous ripple-detection algorithms (*34, 39, 45*), we recorded a total of 35,948 ripples across all sessions (Fig. 2, E–G). Before ripple detection, we conservatively excluded interictal epileptic discharges (IEDs; Text S1; Fig. S2) to help interpret our ripples and ripple-related findings as physiological phenomena (*15*). We characterized the identified ripples with regard to various properties and confirmed that they reflected time periods with strongly elevated power at about 90 Hz (Fig. 2H; Fig. S3), consistent with previous human studies (*34, 39, 45*). These periodic high-frequency events are presumably the human homolog of rodent sharp-wave ripples, although their overlap with human high-gamma or epsilon oscillations (*46, 47*) as well as their relation to ripples in animal models is not yet fully clear (*15*). When we examined the relation between ripples and low-frequency (delta, 0.5–2 Hz) activity of the LFPs, we found that the ripples were preferentially locked to a mean delta phase of 34° (Rayleigh test: *z* = 5.614, *P* = 0.003; Fig. 2I). Thus, consistent with previous results (*21, 33, 48*), ripples in this dataset generally appeared at the descending phase of hippocampal slow oscillations (Fig. 2H), which may have a role in triggering the ripples (*49*).

**Fig. 2.**
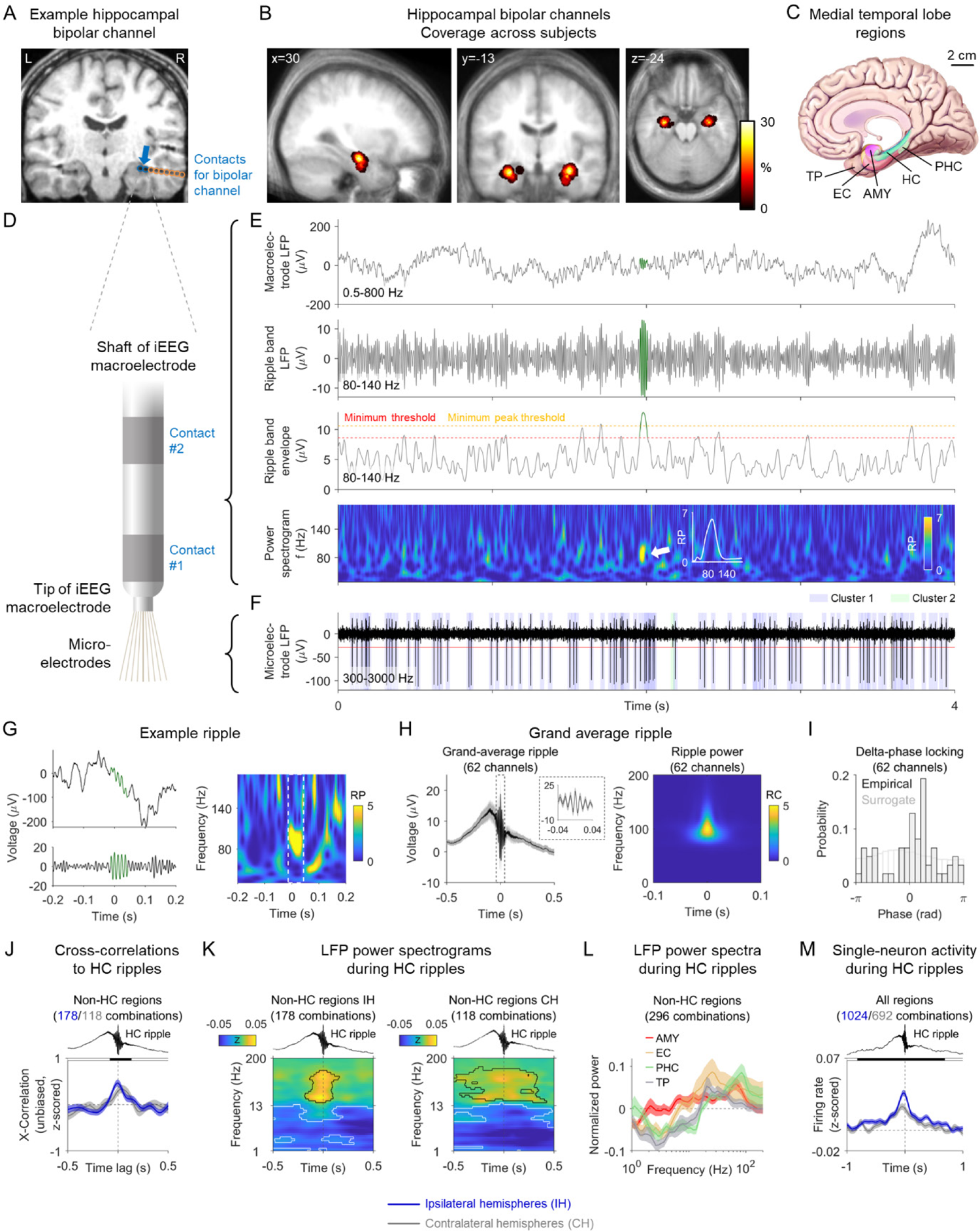
Hippocampal ripples are associated with changes in LFP power and firing rates across the human MTL. (**A**) Location of an example bipolar hippocampal channel (blue arrow). Blue circles, electrode contacts contributing to the bipolar channel; orange circles, other contacts. (**B**) Probability distribution of all bipolar hippocampal channels, overlaid on the subjects’ average MRI scan. (**C**) MTL regions used for the recordings of LFPs and single-neuron activity. (**D**) Illustration of the two innermost electrode contacts of an intracranial EEG macroelectrode with microelectrodes protruding from its tip. (**E**) Analysis procedure for identifying ripples. Top to bottom: raw macroelectrode LFP; macroelectrode LFP filtered in the 80–140 Hz ripple band; smoothed envelope of the ripple-band macroelectrode LFP; spectrogram of the macroelectrode LFP. (**F**) Action potentials of two clusters from a microelectrode simultaneously recorded with the macroelectrode data. (**G**) Raw voltage trace of an example hippocampal ripple (green) in the time domain (left) and its relative power spectrogram in the time-frequency domain (right). Time 0, ripple peak. (**H**) Grand-average voltage trace of hippocampal ripples across all channels in the time domain (left) and their power spectrogram in the time-frequency domain (right). Voltage traces are baseline-corrected with respect to ±3 s around the ripple peak. Ripple power is shown as the relative change with respect to the average power within ±3 s around the ripple peak. Time 0, ripple peak. (**I**) Delta-phase locking of hippocampal ripples. Black histogram, ripple-associated delta phase for each channel. Gray histogram, delta phases of surrogate ripples. (**J**) Cross-correlations between hippocampal ripples and ripples in extrahippocampal MTL regions (temporal pole, amygdala, entorhinal cortex, parahippocampal cortex). Blue and gray numbers indicate the number of ipsilateral and contralateral channel pairs, respectively. Time 0, peak of hippocampal ripples. Cross-correlations are smoothed with a Gaussian kernel of 0.2-s duration and normalized by *z*-scoring cross-correlation values over time lags of ±0.5 s. Shaded region, SEM across channels pairs. Black shading at top indicates cross-correlations from both ipsilateral and contralateral channel pairs significantly above 0 (cluster-based permutation test: *P* < 0.05). (**K**) Time–frequency resolved LFP power (*z*-scored relative to the entire experiment) in extrahippocampal MTL regions during hippocampal ripples. Power values are smoothed over time with a Gaussian kernel of 0.2-s duration. Time 0, ripple peak. Black contours, significantly increased power; white contours, significantly decreased power (two-sided cluster-based permutation tests: *P* < 0.025). (**L**) Normalized LFP power extracted from the time periods of hippocampal ripples and averaged over time. (**M**) Single-neuron firing rates (*z*-scored relative to the entire experiment) in hippocampal and extrahippocampal regions during hippocampal ripples. Firing rates are smoothed over time with a Gaussian kernel of 0.2-s duration. Blue and gray numbers indicate the number of ipsilateral and contralateral neuron–ripple channel pairs, respectively. Black shading at top indicates firing rates of ipsilateral and contralateral pairs significantly above 0 (cluster-based permutation test: *P* < 0.05). AMY, amygdala; EC, entorhinal cortex; HC, hippocampus; PHC, parahippocampal cortex; TP, temporal pole. CH, contralateral hemispheres; IH, ipsilateral hemispheres. LFP, local field potential; RC, relative change; RP, relative power; X-Correlation, cross-correlation.

We furthermore asked how hippocampal ripples related to the subjects’ behavioral state and memory performance in our associative memory task (Text S2). We observed that ripple rates varied as a function of trial period; that increased ripple rates during cue periods were associated with better performance in the subsequent retrieval periods; and that retrieval periods with poorer performance were followed by increased ripple rates during re-encoding (Fig. S4). These direct associations between hippocampal ripples and behavior in our object–location memory task replicate and extend the previously established links between hippocampal ripples and various memory processes in humans, which together suggest that hippocampal ripples are functionally important for encoding and retrieving memories (*33, 36–41, 50*).

### Hippocampal ripples are associated with changes in LFP power and firing rates across the human MTL

Hippocampal ripples are neural events with brain-wide effects that are considered optimal for inducing synaptic plasticity (*13, 21, 51*). We thus reasoned that hippocampal ripples could support associative memory by triggering brain states in which otherwise separate neural representations become linked. To assess the potential for awake hippocampal ripples to generally modulate neural activity across the human MTL, we performed an array of analyses to quantify the effects of ripples on single-neuron spiking and LFP changes in various regions of the human MTL (temporal pole, entorhinal cortex, amygdala, hippocampus, and parahippocampal cortex; Figs. S5 and S6).

We first tested whether ripple events that appeared in extrahippocampal MTL regions were coupled to hippocampal ripples. We identified ripples in extrahippocampal MTL regions using the same procedure as for hippocampal ripples and found that extrahippocampal MTL ripples occurred in temporal proximity with hippocampal ripples using cross-correlation analyses (cluster-based permutation test: *P* < 0.001; Fig. 2J; Fig. S5B). High-frequency oscillatory events therefore seem to be synchronized across the human MTL, in line with previous studies in rodents and humans showing that hippocampal ripples are temporally coupled to ripples in various other brain areas (*23, 33, 37, 52*). This ripple coupling may help to bind separate groups of neurons from different brain regions together.

We also examined how hippocampal ripples related to brain-state changes as reflected in the power of LFP oscillations. Across all macroelectrode channels in both ipsilateral and contralateral extrahippocampal MTL regions, we found that the normalized LFP power at higher frequencies (>20 Hz) increased during hippocampal ripples (cluster-based permutation test across all extrahippocampal MTL channels ipsilateral to hippocampal ripple channels: *P* = 0.018; contralateral: *P* < 0.001). Inversely, normalized power at lower frequencies decreased during hippocampal ripples (ipsilateral: *P* < 0.001; contralateral: *P* < 0.001; Fig. 2, K–L; Fig. S5C). These MTL-wide power changes were strongest at the exact moments when hippocampal ripples occurred but started before and ended after them [see also (*53*)]. Given that increased high-frequency power and decreased low-frequency power are indicators of elevated neuronal excitation (*54*), these results suggest that hippocampal ripples are associated with an overall excitatory state of the human MTL. This increased excitation further indicates that hippocampal ripples support brain states that are suitable for inducing and activating synaptic connections between otherwise segregated neurons.

In a third analysis of the large-scale effects of hippocampal ripples, we examined coincident single-neuron spiking across the MTL at the moments of hippocampal ripples. Across all 27 sessions with single-neuron recordings, we recorded a total of 1063 neurons across multiple regions (Fig. 2, C and F; Fig. S6) including temporal pole, entorhinal cortex, amygdala, hippocampus, and parahippocampal cortex. Overall, neuronal firing rates increased when hippocampal ripples occurred (cluster-based permutation test: *P* < 0.001), where neuronal firing rates started to rise at about 0.25 s before the ripples peaked (Fig. 2M). This increased spiking during hippocampal ripples was strongest for neurons in the hippocampus itself and the amygdala but was also present across cells in the other MTL regions (Fig. S5D). Behavior-related analyses showed that ripple-locked firing-rate increases were largely similar across the different trial phases and for varying levels of memory performance (Text S3; Fig. S4E).

Overall, these results show that hippocampal ripples are associated with broad changes in neural activity across the human MTL. When hippocampal ripples occurred, there was an increased probability of ripples in extrahippocampal MTL regions; a shift in LFP power from lower to higher frequencies in these regions; and an increase in MTL-wide neuronal spiking activity. Related long-range effects of hippocampal ripples have been described in both animals and humans (*21, 22, 24, 37, 53, 55*). Through these excitatory effects, hippocampal ripples may play a key role in the formation and retrieval of associative memories by coordinating and combining diverse patterns of neural activity from multiple regions.

### Neurons in the human MTL encode objects and spatial locations

To examine whether ripples in the human hippocampus are linked to synchronized activity of object- and place-specific cells during associative memory processes, we next tested for these cell types in our data. We identified neurons as object cells if they increased their firing rates in response to a particular object (*9, 10*), and we considered neurons as place cells if they activated when the subject was at a particular spatial location in the virtual environment (*11, 12*).

Each object cell had a particular “preferred object” in response to which they activated most strongly during the cue period. For example, the first object cell shown in Fig. 3A exhibited its highest firing when the subject viewed object 7. We observed 120 object cells (11% of all neurons; binomial test, *P* < 0.001; Table S1), which were most prevalent in the entorhinal cortex, parahippocampal cortex and temporal pole (Fig. 3B). Object cells activated most strongly in response to their preferred object during the first second after cue onset (Fig. 3C); their firing rates returned to baseline shortly after the object disappeared from screen (Fig. 3D); and their tuning curves were overall stable over time [one-sample *t*-test: *t*(118) = 10.387, *P* < 0.001; Fig. 3E]. Most object cells were pure object cells in the sense that they did not also meet the criteria for being place cells (81% of all object cells; Fig. 3F).

**Fig. 3.**
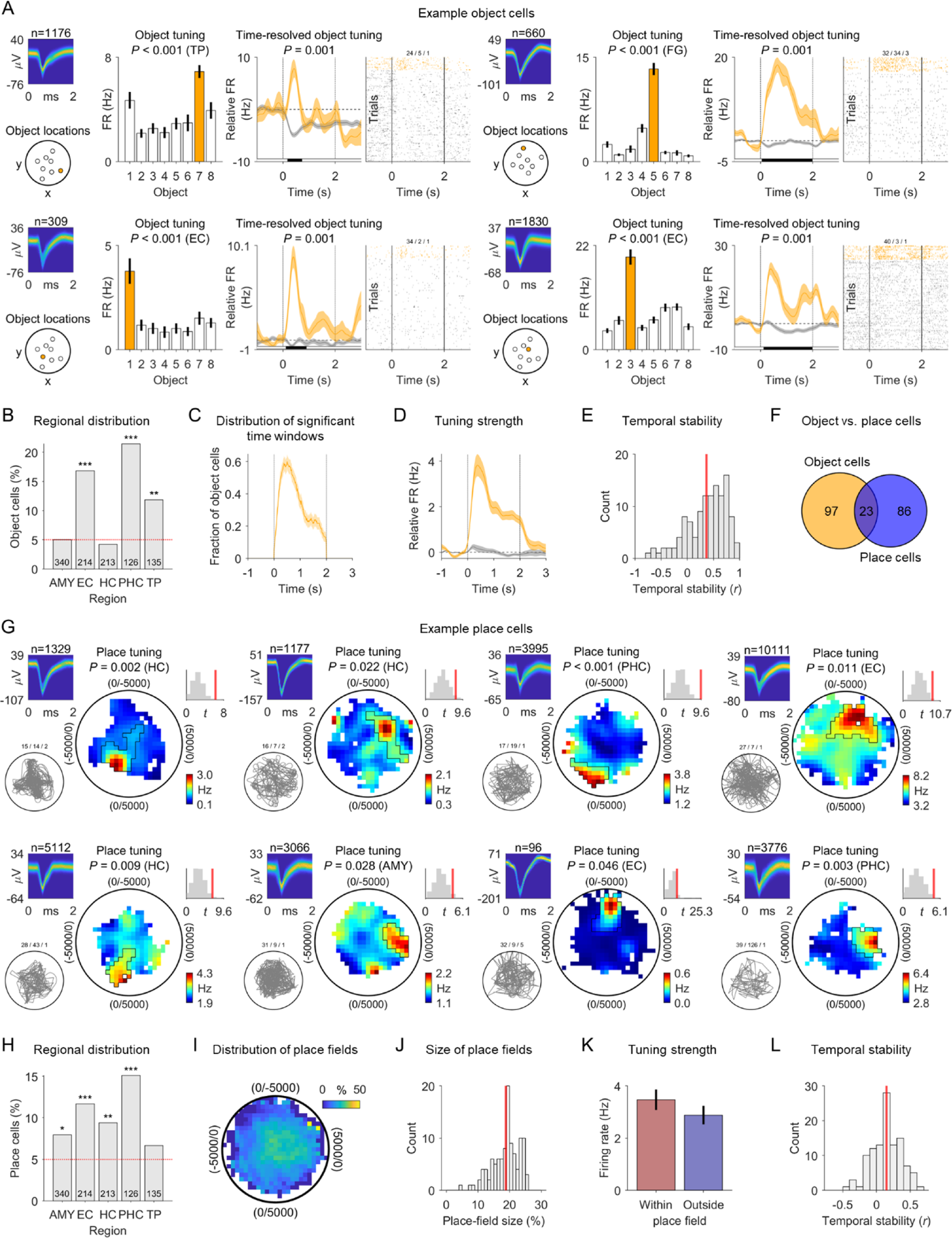
Neurons in the human MTL encode objects and spatial locations. (**A**) Example object cells. For each cell, from left to right: action potentials as density plot; locations of the objects in the environment; absolute firing rates in response to the different objects during the cue period; time-resolved firing rates (baseline-corrected relative to −1 to 0 s before cue onset) for the preferred and the non-preferred objects; spike raster for all trials. Time 0, cue onset. Orange, data for the preferred object; gray, data for unpreferred objects. Error bars, SEM across trials. Black shading below the time-resolved firing rates, significant difference between firing rates (cluster-based permutation test: *P* < 0.05). (**B**) Distribution of object cells across brain regions; red line, 5% chance level. (**C**) Distribution of significant time windows across object cells (mean ± SEM). (**D**) Tuning strength across object cells (mean ± SEM). Orange, data for the preferred object; gray, data for unpreferred objects. (**E**) Temporal stability of object-cell tuning. Red line, mean. (**F**) Overlap between object and place cells. (**G**) Example place cells. For each cell, from left to right: action potentials as density plot; navigation path of the subject through the environment (gray line); smoothed firing-rate map (unvisited areas are shown in white); empirical *t*-statistic (red line) and surrogate *t*-statistics (gray histogram); color bar, firing rate. (**H**) Distribution of place cells across brain regions; red line, 5% chance level. (**I**) Spatial distribution of the place fields of all place cells (in percent relative to the spatial distribution of the firing-rate maps). (**J**) Histogram of place-field sizes, in percent relative to the sizes of the firing-rate maps; red line, mean. (**K**) Firing rate of place cells inside versus outside the place fields; error bars, SEM. (**L**) Temporal stability of the firing-rate maps of place cells; red line, mean. AMY, amygdala; EC, entorhinal cortex; FG, fusiform gyrus; HC, hippocampus; PHC, parahippocampal cortex; TP, temporal pole. \**P* < 0.05; \*\**P* < 0.01; \*\*\**P* < 0.001.

Each place cell activated preferentially when the subject was at a particular location in the virtual environment (Fig. 3G). Over the past decades, studies in rodents have provided ample evidence for such place tuning in neurons of the hippocampus and surrounding brain areas (*11, 56*). Across all neurons, we identified 109 place cells (10%; binomial test, *P* < 0.001) and found them at significant levels in multiple regions, including the entorhinal cortex, hippocampus, and parahippocampal cortex (Fig. 3H), consistent with earlier work (*57*). The firing fields of place cells were broadly distributed across the virtual environment (Fig. 3, I–J), showing that all parts of the environment were neurally represented. The cells’ firing rates were about 20% higher inside versus outside the place fields (Fig. 3K) and their firing patterns were stable over time [one-sample *t*-test: *t*(108) = 7.080, *P* < 0.001; Fig. 3L], indicating that spatial information was robustly encoded by the place cells of our dataset. Place and object tuning was largely independent of each other because place and object cells only marginally overlapped with conjunctive cells, which exhibited significant place tuning solely during trials with one particular object (Fig. S7).

Together, these results show that cells in the human MTL encoded the separate elements of to-be-established and to-be-remembered object–location memories in our task: object cells encoded individual objects and place cells encoded particular spatial locations. This allowed us to next probe the coactivity of object and place cells during hippocampal ripples as a potential neural correlate of associative memory.

### Ripple-locked coactivity of object and place cells during the retrieval and formation of associative memories

Having examined hippocampal ripples and stimulus-specific (object and place) neurons separately, we next tested our principal hypothesis that object and place cells would activate together during the same hippocampal ripples when subjects formed and retrieved associative memories that linked the separate elements these neurons encoded (Fig. 1A). We reasoned that coactivity during memory formation would facilitate synaptic connections between the stimulus-specific neurons (*58*) and that coactivity during memory retrieval would reflect their reciprocal activation through their previously established synaptic connections.

To investigate our hypothesis, we analyzed whether simultaneously recorded object and place cells were active during the same ripples and performed this analysis separately for ripples during retrieval and during re-encoding. Briefly, for each combination of an object cell, a place cell, and a ripple channel, we computed coactivity scores between the two cells across ripples that indicated how often both cells were (in)active during the same ripples (controlling for the overall activity level of both cells). We systematically and independently varied the ripple-locked time point for determining the underlying activity of both cells and thus obtained a two-dimensional time-by-time coactivity map for each cell pair. These coactivity maps showed the coactivity scores at various time points between −0.25 s and +0.25 s around the ripple peaks, where high coactivity scores indicated a consistent across-ripple coactivation of both cells at specific time points relative to the ripple peaks (Fig. 4A; Fig. S8).

**Fig. 4.**
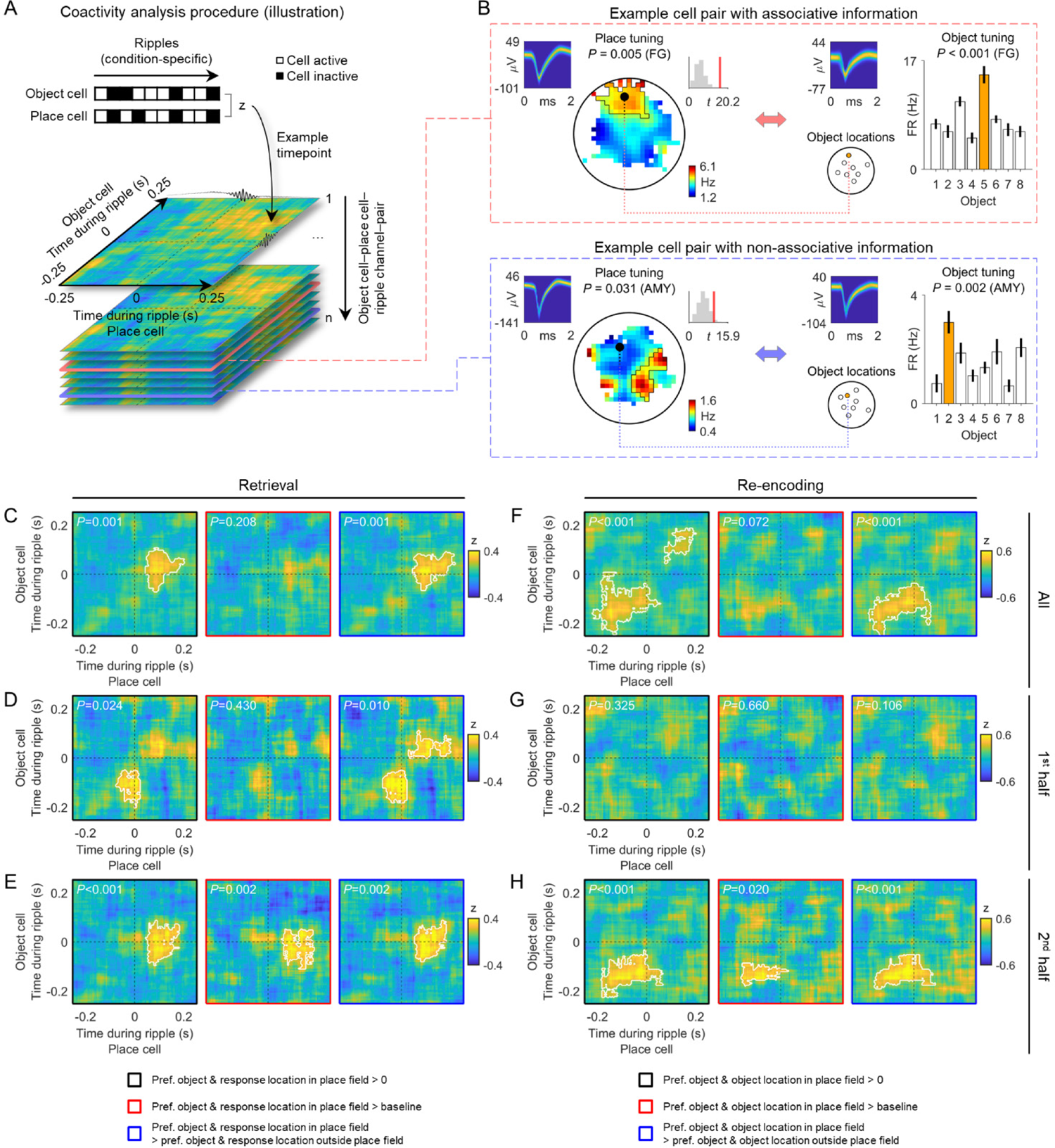
Ripple-locked coactivity of object and place cells during the retrieval and formation of associative memories. (**A**) Analysis of ripple-locked coactivity of object and place cells (illustration). High coactivity *z*-values occur when the object cell and the place cell are (in)active during the same ripples (top). Two-dimensional coactivity *z*-score maps are calculated across ripples for pairs of object and place cells, separately for various time points relative to the ripple peak (bottom). (**B**) Example pairs of object and place cells with associative (top) and non-associative information (bottom). In cell pairs with associative information, the place field of the place cell contains the remembered location (retrieval) or the correct location (re-encoding) of the preferred object of the object cell. (**C**) Coactivity maps estimated using all ripples during retrieval periods, considering only trials in which the subject is asked to remember the location of the preferred object of the object cell and in which the subject’s response location is inside the place field of the place cell. Left, comparison of the coactivity maps against chance (i.e., 0). Middle, comparison against baseline coactivity maps. Right, comparison against coactivity maps estimated using ripples from trials in which the subject is asked to remember the location of the preferred object of the object cell and in which the subject’s response location is outside the place field of the place cell. (**D**) Same as in C for the first half of all retrieval-related hippocampal ripples. (**E**) Same as in C for the second half of all retrieval-related hippocampal ripples. (**F**) Coactivity maps estimated using all ripples from the re-encoding periods, considering only trials in which the subject is asked to re-encode the correct location of the preferred object of the object cell and in which the object’s correct location is inside the place field of the place cell. Left, comparison of the coactivity maps against chance (i.e., 0). Middle, comparison against baseline coactivity maps. Right, comparison against coactivity maps estimated using ripples from trials in which the subject is asked to re-encode the location of the preferred object of the object cell and in which the object’s correct location is outside the place field of the place cell. (**G**) Same as in F for the first half of all re-encoding-related hippocampal ripples. (**H**) Same as in F for the second half of all re-encoding-related hippocampal ripples. White lines delineate significant clusters based on cluster-based permutation tests, which control for multiple comparisons and whose *P*-values are stated in the upper left corners of the coactivity maps (see the main text for Bonferroni corrected *P*-values). AMY, amygdala; FG, fusiform gyrus; pref., preferred.

Based on our principal hypothesis (Fig. 1A), our analysis focused on the ripple-locked coactivity of pairs of object and place cells where the preferred object of the object cell was located inside the place field of the place cell. Cells in these “associative cell pairs” represented the different pieces of information that the subjects had to associate with each other (Fig. 4B). For retrieval periods, the location of the preferred object of the object cell was defined by the subject’s response locations, whereas during re-encoding periods it was defined by the object’s true location. We hypothesized that coactivations of associative object and place cells during retrieval would indicate that the subject remembered a particular location given a specific object, whereas their coactivations during re-encoding would indicate that the subject aimed at (re)learning the correct location of a given object. Overall, we thus reasoned that the coactivity of associative cell pairs could underlie the encoding and retrieval of particular associative memories.

To evaluate the statistical significance of the coactivity for pairs of object and place cells encoding associative information on each trial, we designed a series of three complementary statistical tests (Fig. S9). These tests contrasted the coactivity maps (i) against chance to see whether the cell pairs showed a positive increase in coactivity; (ii) against coactivity maps from a baseline period to show that the coactivity increases were specific to the timing of ripples; and (iii) against the coactivity maps of non-associative object and place cell pairs (where the preferred object of the object cell was located outside the place cell’s place field) to demonstrate that the coactivity increases were unique to cell pairs representing the exact components of the associative memories. Together, these three tests provided robust information about whether associative pairs of object and place cells exhibited coactivity during hippocampal ripples.

We first examined ripple-locked coactivity of associative object cell–place cell pairs during retrieval (Fig. 4, C–E; Fig. S10, A–C; Fig. S11A). Across all retrieval periods, we found indications of ripple-locked coactivity in associative cell pairs as their coactivity scores were significantly positive and significantly greater than in control cell pairs that did not encode associative information (cluster-based permutation test vs. chance: *P* = 0.001; vs. non-associative cell pairs: *P* = 0.001; Fig. 4C). The comparison against the coactivity maps from the baseline period was not significant (*P* = 0.208), however, which indicates that ripple-locked coactivity of object and place cells during retrieval was not fully developed when estimated across the entire task. We therefore performed this analysis separately for “early” hippocampal ripples (first half of all ripples per session) and “late” ripples (second half of all ripples per session) because we hypothesized that retrieval-related coactivity would be expressed more strongly during late ripples, after the subjects had already formed associations between the objects and their corresponding locations (Fig. 1D). Indeed, when only considering late ripples, we found clear ripple-locked coactivity of object cell–place cell pairs representing associative information as the coactivity maps of these cell pairs were significant for all three statistical tests (vs. chance: *P* = 0.001; vs. baseline: *P* = 0.005; vs. non-associative cell pairs: *P* = 0.004; *P*-values are Bonferroni corrected for performing this analysis on both early and late ripples; Fig. 4E). Similarly strong effects were not present when considering early ripples (vs. chance: *P* = 0.048; vs. baseline: *P* = 0.860; vs. non-associative cell pairs: *P* = 0.020; Bonferroni corrected; Fig. 4D), which explains why ripple-locked coactivity during retrieval was not fully developed when considering all ripples.

These findings demonstrate that ripples from later retrieval periods were associated with coactivations in pairs of object and place cells that represented associative information of to-be-retrieved associative memories. We assume that the synaptic connections between object and place cells gradually developed over time, which is why the ripple-locked coactivity of associative object and place cells was not robustly visible during early retrieval periods. Follow-up analyses confirmed these results by showing that ripple-locked coactivity was present at the level of individual cell pairs (Fig. S12) and that the same effects appeared with another method for estimating coactivity maps based on Pearson correlations (Fig. S13). The coactivity maps showed that the increased coactivations occurred at the moment of the ripple peaks for object cells and slightly after the ripple peaks for place cells (Fig. 4E; Fig. S14A), which may be related to the propagation of information from the hippocampus to extrahippocampal regions during retrieval.

To next understand the contribution of ripple-coordinated single-neuron activity to the formation of associative memories, we examined ripple-locked coactivity of the object cell–place cell pairs that represented associative information during the re-encoding periods of our task (Fig. 4, F–H; Fig. S10, D–F; Fig. S11B). Similar to our retrieval-related results, we found indications of increased coactivity in associative object cell–place cell pairs when considering ripples from all re-encoding periods, whereby the comparisons against chance and non-associative cell pairs were significant, but the comparison against the baseline period was not significant (vs. chance: *P* < 0.001; vs. baseline: *P* = 0.072; vs. non-associative cell pairs: *P* < 0.001; Fig. 4F). Parallel to our coactivity analyses during retrieval, we therefore examined re-encoding-related coactivations separately for early and late ripples. Indeed, we found again that when only late ripples were considered associative pairs of object and place cells showed robust increases in coactivity that were significant for all three statistical tests (vs. chance: *P* = 0.001; vs. baseline: *P* = 0.040; vs. non-associative cell pairs: *P* < 0.001; *P*-values are Bonferroni corrected for performing this analysis on both early and late ripples; Fig. 4H; Figs. S12 and S13). In contrast, such coactivations were not present for early re-encoding-related ripples (vs. chance: *P* = 0.650; vs. baseline: *P* = 1; vs. non-associative cell pairs: *P* = 0.212; Bonferroni corrected; Fig. 4G), which again explains why ripple-locked coactivity during re-encoding was not fully developed when measured across the entire session.

This set of results demonstrates that ripple-locked coactivity between associative object and place cells occurred when subjects re-encoded object–location associations during later periods of the task. During these later task periods, the subjects were already familiar with the associations. We therefore speculate that ripple-locked coactivity during re-encoding is more closely related to the stabilization, updating, or early consolidation of associative object–location memories rather than to their initial formation (*6, 14, 31*). In contrast to the timing of ripple-locked coactivity during retrieval, the increased coactivations during late re-encoding-related ripples shifted to slightly precede the hippocampal ripples (Fig. 4H; Fig. S14B), which may reflect the propagation of information from extrahippocampal regions to the hippocampus during memory encoding.

To further investigate whether the distinction between early and late ripples paralleled a distinction between ripples occurring before versus after the initial formation of the object–location memories, we estimated the time of strongest improvement in memory performance for each object. We reasoned that this time reflected the moment when the object–location associations emerged initially. We then grouped the ripples according to whether they occurred before or after this time of strongest memory improvement. Ripple-locked coactivations of object and place cells before and after initial memory formation appeared very similar to their coactivity patterns during early and late ripples, respectively (Fig. S15). This suggests that the coactivations during late ripples from both retrieval and re-encoding periods were at least partly dependent upon an initial formation of the object–location associations.

Studies in rodents highlighted the functional relevance of ripples during immobility (*31, 32*). We thus differentiated between ripples from movement or non-movement periods and examined whether the coactivity effects appeared preferentially during non-movement-related ripples. As hypothesized, the increased ripple-locked coactivity between associative object and place cells were driven by ripples occurring during non-movement periods (Fig. S16).

Overall, these results provide empirical support for our principal hypothesis that human hippocampal ripples are associated with coactivations of stimulus-specific neurons (which represent particular objects and locations in this study) during the formation and retrieval of associative memories.

## Discussion

Associative memory allows us to learn and later retrieve new links and relations between previously unrelated stimuli and is thus an essential cognitive function for everyday life (*1*). Here, we conducted neural recordings in epilepsy patients performing an associative object–location memory task in a virtual environment (Fig. 1) to investigate the role of stimulus-specific single neurons and hippocampal ripples for human associative memory. We found that object- and place-specific neurons (Fig. 3) are simultaneously active during hippocampal ripples (Fig. 2) when subjects form and retrieve particular associative object–location memories (Fig. 4). This mechanism of ripple-locked coactivity of stimulus-specific neurons suggests a cellular account of how humans interconnect previously unrelated stimuli and how they recall a stimulus from memory after being cued with another stimulus. Hippocampal ripples may support these functions by binding distinct cellular populations together.

### Neural mechanisms of associative memory

Different theories suggest explanations of how neural circuits in the brain encode associative memories (*1*). Following the “conjunctive hypothesis,” the relevant neural circuits may contain conjunctive representations as a neural substrate for associative memories, in which neurons encode the unique combination of two or more stimuli (*17*). Such conjunctive neurons do not respond to the different stimuli in isolation but only when the stimuli are encoded or retrieved together (*18–20, 59*). In contrast, following the “coactivity hypothesis,” associative memories may be enabled by stimulus-specific neurons that encode the individual elements of associative memories and that coactivate temporarily when subjects encode or recall these associative memories (*6–8*). Coactivity during memory encoding would establish synaptic connections between the stimulus-specific neurons (*51, 58*), and coactivity during retrieval would reflect the reciprocal activation of stimulus-specific neurons based on their previously established synaptic connections.

In line with the “conjunctive hypothesis,” several studies described the existence of conjunctive cells in MTL regions. For example, in rodents, the spiking of hippocampal neurons encodes particular object–location associations and this conjunctive code strengthens with learning (*19*). Similarly, single neurons in the monkey hippocampus change their firing rates when monkeys learn particular scene–eye movement associations (*18*), which indicates that these neurons represent a particular eye movement in a given spatial context after associative learning has occurred [see also (*60*)]. In humans, MTL neurons rapidly adapt their selectivity to represent person–scene conjunctions (*20*) and hippocampal neurons respond to unique combinations of more than two stimuli (*59*). These studies support the view that associative memory is linked to conjunctive neural coding.

Our study provides empirical support for the “coactivity hypothesis” of associative memory by identifying single-neuron representations of the separate memory elements and by demonstrating how these separate neural representations are simultaneously active at particular time points. Specifically, our results show that stimulus-specific object and place cells coactivate during hippocampal ripples when humans encode or retrieve object–location associations. We found that this ripple-locked coactivity was specific to object and place cells jointly representing associative information (Fig. 4), that it was significantly expressed only during the second half of the task (Fig. 4), and that it occurred mostly after initial memory formation had taken place (Fig. S15). These results support the “coactivity hypothesis” by indicating that precisely timed cellular coactivations play a role in human associative memory. Given that not only associative memories require an interplay between different mental contents, we propose that these findings are relevant to various cognitive functions that involve transient interactions between otherwise independent neural representations. For example, episodic memories comprise event, time, and place information and may thus rely on transient interactions between concept cells containing semantic information (*10*), time cells coding temporal information (*61*), and spatially modulated cells representing locations and directions (*9, 11, 12*).

The “conjunctive hypothesis” and the “coactivity hypothesis” are not mutually exclusive. One possible scenario is that neural activity in line with the “coactivity hypothesis” leads to the emergence of neural activity proposed by the “conjunctive hypothesis.” For instance, simultaneous coactivity in object and place cells may induce conjunctive object–place coding in downstream neurons.

### Hippocampal ripples and cognition

A large body of previous work, primarily performed in rodents, suggests that hippocampal ripples serve multiple cognitive functions including memory encoding, consolidation, and retrieval (*13, 14, 31, 32*). For example, classical findings in rodents demonstrated that the suppression of sharp-wave ripples during post-training sleep impairs spatial memory (*62*), providing evidence for a role of ripples in memory consolidation. Studies in awake humans showed that ripple rates increase when subjects encode new memories (*36, 41*) and when they freely recall memories (*36, 41, 50*), implicating ripples in both memory encoding and retrieval. Moreover, ripple-associated place-cell sequences depict the animal’s future paths through an environment, which indicates that hippocampal ripples help plan future behavior (*13, 27, 29*).

In this study, we extended the evidence for important roles of hippocampal ripples in cognition by detailing their involvement in human associative memory. Specifically, hippocampal ripples may define time windows for coactive spiking by stimulus-specific neurons that represent different types of information. During the retrieval of object–location memories, we observed this ripple-locked coactivity on trials when the subject was asked to recall the location of the preferred object of the object cell and in which the subject’s response location was inside the place field of the place cell. This result implicates human ripples in the retrieval of associative memories and also supports their implication in planning future behavior, as the coactivity occurred on trials in which the subject’s response location—which is the location the subject was heading to—was located inside the place field of the place cell. As we observed significant coactivations only during the second half of each session, it suggests that subjects first had to establish some intuition about the locations of the different objects before ripple-locked coactivity during retrieval could emerge.

During re-encoding periods of our object–location memory task, we observed coactivity of object and place cells on trials when the subject was asked to re-encode the correct location of the preferred object of the object cell and when the correct location of the object was inside the place field of the place cell. This coactivity of object and place cells during re-encoding may thus have helped the subjects to build or stabilize accurate associations between the different objects and their spatial locations, implicating ripples in memory formation. We again observed significant ripple-locked coactivity of object and place cells only during the second half of the recording sessions. We therefore propose that ripple-locked coactivity during re-encoding relates more closely to the stabilization, updating, or early consolidation of associative memories rather than to their initial formation (*6, 14, 31, 63, 64*). The differences between memory initialization, stabilization, updating, and early consolidation are not exactly defined, though, and hippocampal ripples may be involved in all of these cognitive operations (*32*).

Our results suggest that the ripple-locked timing of neuronal coactivity shifts between different behavioral states. During retrieval, the strongest object cell–place cell coactivity occurred shortly after the peaks of hippocampal ripples, whereas during re-encoding the coactivations mainly preceded hippocampal ripple peaks (Fig. 4; Fig. S14). This timing shift may reflect task-related changes in the direction of information flow during hippocampal ripples. Specifically, hippocampal ripples during retrieval may induce the coactivity of stimulus-specific neurons and may support the propagation of information from the hippocampus to extrahippocampal regions. Conversely, during re-encoding, hippocampal ripples may be triggered by stimulus-specific neuronal activity and may be involved in information transfer from extrahippocampal regions to the hippocampus, which could also generate conjunctive hippocampal representations. Our finding of object cell–place cell coactivity starting prior to hippocampal ripples is in line with prior results in rodents showing that place-cell sequences can start about 100 ms earlier than the hippocampal ripple (*26, 27*) and that the reactivation of brain-wide neural representations of word pairs, faces, and scenes emerges already about 100 ms before hippocampal ripples in humans (*36, 37*). This temporal delay raises the question whether hippocampal ripples actually trigger the (re)activation of stimulus-specific neural representations or whether another mechanism—for example, the excitatory sharp wave (*13, 32*)—is in fact the trigger event for both the (re)activation of stimulus-specific neural representations and the hippocampal ripples.

A further way in which our study implicates hippocampal ripples in human associative memory is by showing that ripple rates correlated with the subjects’ behavioral state and memory performance in our task (Fig. S4). Ripple rates increased during memory cues prior to good memory retrieval and during re-encoding periods after poor memory retrieval. These observations match previous reports that hippocampal ripples are important for the accurate encoding and retrieval of human memories (*33, 35–42, 50*) and help extend the functional role of hippocampal ripples from the rodent to the human brain (*15*). Moving forward, these insights may have translational relevance, as augmenting ripples (*30*) might be a useful target for improving cognitive performance in patients with memory disorders (for further discussion points, see Text S4).

## Acknowledgements

We are very grateful to all patients who participated in this study. We thank the clinical team of the Freiburg Epilepsy Center, Freiburg, Germany, for their continuous support. We thank Daniel Lachner-Piza for help with data collection and Kamran Diba, John J. Sakon, Serra E. Favila, and Tamara Gedankien for comments on the manuscript.

## Author contributions

L.K. developed hypotheses and designed the study; L.K., J.J., and A.S.-B. acquired funding; L.K., P.C.R., A.B., and A.S.-B. recruited study participants; P.C.R. implanted electrodes; L.K., A.B., and T.A.G. collected the data; L.K. analyzed the data; L.K., J.J., and B.P.S. discussed the results; L.K. and J.J. wrote the paper; all authors reviewed and revised the final manuscript.

## Data and materials availability

Further information and requests for resources and reagents should be directed to and will be fulfilled by the lead contact, Lukas Kunz. All custom MATLAB code generated during this study for data analysis and the data to recreate the figures will be made publicly available upon publication. Raw data are not publicly available because they could compromise research participant privacy, but are available upon request from the lead contact, Lukas Kunz. Any additional information required to reanalyze the data reported in this paper is available from the lead contact upon request.

## Supplementary Materials

### Materials and Methods

#### Human subjects

We tested *N* = 35 human subjects, who were epilepsy patients undergoing treatment for pharmacologically intractable epilepsy at the Freiburg Epilepsy Center, Freiburg, Germany. Of those, 5 patients had to be excluded because of technical issues (*n* = 1); no hippocampal electrode contacts (*n* = 2); hippocampal channels that were close to the resection border of a previous surgery (*n* = 1); and a very low number of ripples (*n* = 1). This resulted in a final sample of *n* = 30 subjects (16 female; age range, 19–61 years; mean age ± SEM, 36 ± 2 years), contributing a total of *n* = 41 experimental sessions with intracranial EEG recordings including the left and/or right hippocampus (*n* = 62 hippocampal bipolar channels). For 20 of these 30 subjects, additional single-neuron recordings from various MTL regions were available (*n* = 27 sessions; *n* = 43 hippocampal bipolar channels). Further subject information is presented in Table S1. The study conformed to the guidelines of the ethics committee of the University Hospital Freiburg, Freiburg, Germany, and all patients provided informed written consent.

#### Neurophysiological recordings

Patients were surgically implanted with intracranial depth electrodes in the MTL for diagnostic purposes in order to isolate their epileptic seizure focus for potential subsequent surgical resection. The exact electrode numbers and locations varied across subjects and were determined solely by clinical needs. Electrodes were provided by Ad-Tech (Ad-Tech, Racine, WI, USA). Macroelectrode recordings were performed at a sampling rate of 2 kHz using a Compumedics system (Compumedics, Abbotsford, Victoria, Australia). Microelectrode recordings were performed using Behnke-Fried electrodes (Ad-Tech, Racine, WI, USA) at a sampling rate of 30 kHz using a NeuroPort system (Blackrock Microsystems, Salt Lake City, UT, USA). Each Behnke-Fried electrode contained a bundle of 9 platinum–iridium microelectrodes with a diameter of 40 µm that protruded from the tip of the macroelectrode (*43*). The first eight microelectrodes were used to record action potentials and local field potentials. The ninth microelectrode served as reference. Microelectrode coverage included amygdala (AMY), entorhinal cortex (EC), fusiform gyrus (FG), hippocampus (HC), insula, parahippocampal cortex (PHC), temporal pole (TP), and visual cortex (VC). Temporal pole refers to a broader area in the anterior temporal lobe, situated ventral and anterior to the amygdala and entorhinal cortex.

#### Associative object-location memory task

During the invasive neural recordings, patients sat in their hospital bed and performed an associative object–location memory task in a virtual environment on a laptop computer (Fig. 1), which was adapted from previous studies (*9, 65, 66*). The task was developed using Unreal Engine 2 (Epic Games, Cary, NC, USA).

During the task, subjects first learned the locations of eight everyday objects by collecting each object from its location once (this initial encoding period was excluded from all analyses because it only lasted a few minutes). Afterwards, subjects completed variable numbers of test trials depending on compliance. Each test trial started with an inter-trial-interval (“ITI”) of 3–5-s duration (uniformly distributed). Subjects were then shown one of the eight objects (“cue”; duration of 2 s). During the subsequent retrieval period (“retrieval”; self-paced), subjects navigated to the remembered object location and indicated their arrival with a button press. Next, subjects received feedback on the accuracy of their response using one of five different emoticons (“feedback”; duration of 1.5 s). The retrieved object then appeared in its correct location and the subjects collected it from there to further improve their associative object–location memories (“re-encoding”; self-paced). Across all trials, the average duration of the retrieval periods was 13.415 ± 0.218 s (mean ± SEM) and the average duration of the re-encoding periods was 7.685 ± 0.158 s (mean ± SEM). The subjects could use several different strategies to retrieve the locations of the objects, including allocentric, egocentric, and landmark-based strategies (*9*).

The virtual environment comprised a grassy plain with a diameter of ∼10000 virtual units (vu), surrounded by a cylindrical cliff. There were no landmarks within the environment. The background scenery comprised a large and a small mountain, clouds, and the sun. All distal landmarks were rendered at infinity and remained stationary throughout the task. Subjects were asked to complete up to 160 trials but were instructed to pause or quit the task whenever they wanted. Subjects navigated the virtual environment using the arrow keys of the laptop computer (forward; turn left; turn right). Instantaneous virtual locations and heading directions (which are identical with viewing directions in our task) were sampled at 50 Hz. We aligned the behavioral data with the macroelectrode and microelectrode data using visual triggers, which were detected by a phototransistor attached to the screen of the laptop computer. The phototransistor signal was recorded together with the macroelectrode and microelectrode data at temporal resolutions of 2 kHz and 30 kHz, respectively.

#### General information on statistics

All analyses were carried out in MATLAB 2020b (The MathWorks, Inc., Natick, MA, USA) using MATLAB toolboxes, the CircStat toolbox (*67*), FieldTrip (version 20210614; http://fieldtriptoolbox.org) (*68*), and custom MATLAB scripts. All custom MATLAB scripts will be publicly available on GitHub upon publication.

Unless otherwise indicated, we considered results statistically significant when the corresponding *P*-value fell below an alpha level of α = 0.05. Analyses were two-sided, unless otherwise specified. Binomial tests evaluated the significance of proportions of neurons relative to a chance level of 5%, unless otherwise specified. Surrogate statistics were one-sided to assess whether an empirical test statistic significantly exceeded (or fell below) a distribution of surrogate statistics, unless otherwise specified. Correction for multiple comparisons was applied when necessary. All cluster-based permutation tests controlled for multiple comparisons across all relevant data dimensions.

#### Behavioral analysis

For each trial, we quantified the subject’s associative memory performance by calculating the Euclidean distance between the subject’s response location and the object’s correct location (“drop error”). Drop errors were transformed into memory performance values by ranking each drop error within a distribution of surrogate drop errors (*n* = 10^7^). Surrogate drop errors were generated synthetically as the distances between the trial-specific correct object location and random locations within the virtual environment. The transformation into memory performance values accounted for the fact that the possible range of drop errors is smaller for object locations in the center of the virtual environment as compared to object locations in the periphery of the virtual environment (*9, 69*): For objects in the environment center, the potential drop errors are in the range between [0, *r*], whereas they are in the range between [0, 2\**r*] for objects in the periphery of the arena, where *r* is the arena radius. Using the transformation procedure, performance values are mapped onto a range between [0, 1], irrespective of whether the associated objects are located in the center or the periphery of the environment. A memory performance value of 1 represents the smallest possible drop error, whereas a memory performance value of 0 represents the largest possible drop error.

To quantify performance increases within sessions, we computed the change in memory performance between the first and the second half of all trials (averaged across trials). We observed a significant increase in memory performance across all subjects and when only considering subjects with microelectrodes (Table S2). We also estimated memory-performance values per normalized time bin (20 bins; averaged across trials falling into a given normalized time bin) and calculated a Pearson correlation between bin index and average memory performance afterwards (averaged across sessions).

#### Intracranial EEG: preprocessing

Intracranial macroelectrode recordings were performed to identify ripples on hippocampal channels and to examine the MTL-wide effects of hippocampal ripples on local field potentials. Signals were sampled at 2 kHz and initial recordings were referenced to a common surface EEG contact (Cz). We visually inspected the data from each channel and removed channels without reasonable signals (for example, because they were located outside the brain). This resulted in a total number of 2,897 intracranial EEG channels across all 41 sessions (519 out of 3,416 channels were removed because of artifactual data).

To eliminate potentially system-wide artifacts or noise and to better sense ripples locally, we then applied bipolar re-referencing between pairs of neighboring contacts (*37, 38, 50*). Following bipolar re-referencing, we used band-stop filters to perform line-noise removal at 50, 100, 150, and 200 Hz (±2 Hz; two-pass 4^th^ order Butterworth filter) in FieldTrip.

To remove time periods with ripple-like artifacts that were present on the majority of all intracranial EEG channels, we computed the grand-average signal across all intracranial EEG channels (*36, 40*). This grand-average signal was also band-stop filtered at 50, 100, 150, and 200 Hz to remove line noise.

#### Intracranial EEG: electrode locations

To identify which bipolar channels were located inside the hippocampus and could thus be used to detect hippocampal ripples, we visually inspected all hippocampal electrodes on the post-implantation MRI scans (Fig. S1). Following the hippocampal segmentation in (*36*), we assigned the hippocampal bipolar channels to particular hippocampal subregions, which showed that most bipolar channels were located in CA1. We similarly inspected all amygdala, entorhinal cortex, parahippocampal cortex, and temporal pole electrodes to identify which bipolar channels were located in these regions (to examine the MTL-wide effects of hippocampal ripples). The locations of all bipolar channels in the different regions are displayed in Fig. S5A.

To display a summary of all hippocampal bipolar channels, we determined the location of each macroelectrode channel in MNI space using PyLocator (http://pylocator.thorstenkranz.de/), following our previous procedure (*70*), and estimated the location of each bipolar channel as the mean of the MNI coordinates of the two contributing channels. We display the distribution of all hippocampal bipolar channels as a probability map on the group-average MRI scan (Fig. 2B).

#### Intracranial EEG: interictal epileptic discharges (IEDs)

To reduce the probability that the detected ripples were a result of IEDs (*38*), we identified IEDs before ripple detection using an automated procedure, which we double-checked with visual inspection. We automatically detected IEDs following previously established methods (*33, 34, 45*). The raw data was filtered using a high-pass filter of 0.5 Hz (two-pass 5^th^-order Butterworth filter) to remove slow-frequency drifts and a low-pass filter of 150 Hz (two-pass 6^th^-order Butterworth filter) in Fieldtrip. A time point was considered belonging to an IED, if (i) its amplitude exceeded four inter-quartile ranges above or below the median amplitude calculated across the entire recording; if (ii) the gradient to the next time point exceeded four interquartile ranges above or below the median gradient; or if (iii) the sum power across the frequencies 1–60 Hz (30 logarithmically spaced frequencies; time-frequency decomposition using Morlet wavelets with 7 cycles, followed by taking the natural logarithm and frequency-specific *z*-scoring across time) exceeded four interquartile ranges above the median sum power. The rationale behind these criteria was that IEDs exhibit high amplitudes, sharp amplitude changes, and power increases across a broad frequency range. We used interquartile ranges instead of standard deviations to reduce the influence of outliers. We inspected the output of our automated IED detection, which appeared suitable for detecting IEDs. Example IEDs are shown in Fig. S2A.

#### Intracranial EEG: relationship between IEDs and behavior

To test for systematic relationships between hippocampal IEDs and behavior, we estimated the prevalence of IEDs per trial phase. Using a repeated measures ANOVA (dependent variable: IED prevalence, averaged across trials; independent variable: trial phase; Tukey-Kramer correction for multiple comparisons), we tested whether the occurrence of IEDs varied as a function of trial phase.

To understand whether IEDs were associated with memory performance, we performed partial correlations between IED prevalence and memory-performance values across trials, separately for each trial phase (controlling for trial index and the interaction between memory performance and trial index). We then computed one-sample *t*-tests across channels to see whether the correlations were significantly above or below 0, including Bonferroni correction for the number of trial phases. Moreover, to test whether IEDs increased or decreased over the course of the task, we performed partial correlations between IED prevalence and trial indices across trials, separately for each trial phase (controlling for memory performance and the interaction between memory performance and trial index). To corroborate the results from the partial correlations, we also computed linear mixed models with IED prevalence as dependent variable and various independent variables (Table S3).

#### Intracranial EEG: ripples

We detected hippocampal ripples on bipolar channels of hippocampal macroelectrodes. If a subject was implanted bilaterally, hippocampal channels from both hemispheres were used for ripple detection. We first ensured reasonable signals on each hippocampal ripple channel by visually inspecting the raw intracranial EEG traces during preprocessing. If the signal of the most medial bipolar hippocampal channel did not have sufficient quality (which was the case in 5 of the 62 hippocampal channels), we used the second most medial bipolar hippocampal channel for ripple detection. In one patient, the hippocampal electrode was implanted from posterior to anterior along the longitudinal axis of the hippocampus—in this case, we selected the two most anterior hippocampal channels so that the resulting bipolar channel was located in a similar hippocampal subregion as the channels from all other patients (i.e., in the anterior hippocampus). For each bipolar hippocampal channel, we visually verified that it was located inside the hippocampus. Most hippocampal channels were located in the CA1 region (Fig. S1). In total, 62 hippocampal channels were included (33 from the right hemisphere). 43 of these channels were from subjects with microelectrode recordings.

Ripple detection was preceded by a detection of IEDs (see above) and ripple-like events in the grand-average signal to exclude those time periods from ripple detection. To reduce the probability that the detected ripples were a result of artifacts, we conservatively excluded ±1 s around each detected IED (*34*) and each time point that co-occurred with a ripple-like event in the grand-average signal.

To detect ripple candidates, we filtered the raw LFP between 80 and 140 Hz (two-pass 4^th^ order Butterworth filter) and computed the instantaneous analytic amplitude within that band using a Hilbert transform (*37, 38*). We then smoothed the amplitudes using a smoothing time window of 20 ms (*45*). Time points with artifacts were excluded (i.e., set to NaNs) in the smoothed amplitude time series. Next, we computed the mean and standard deviation of the smoothed amplitudes across the entire recording and detected candidate ripple events as time periods in which the signal exceeded 2 standard deviations above the mean (*37–39*). Each candidate event then had to fulfill additional criteria to qualify as a putatively physiological ripple: (i) the peak of the smoothed amplitude had to exceed 3 standard deviations above the mean (*37, 38*); (ii) the duration had to last longer than 20 ms (*36*) and be shorter than 500 ms (*39*); (iii) the bandpass signal needed to have at least 3 peaks and at least 3 troughs (*45*); and (iv) the power spectrum, computed for frequencies between 30 and 190 Hz in steps of 2 Hz (using Morlet wavelets with 7 cycles) and divided by the power spectrum estimated across the entire recording, had to exhibit a global peak between 80 and 140 Hz. Only candidate events that fulfilled all these criteria were considered as ripples.

Next, for each ripple, we extracted its peak time as the time point at which the band-pass signal was highest. Ripple duration was defined as the time difference between the start and end time of a given ripple. The frequency of a ripple was estimated based on the average temporal delay between the peaks and troughs in the band-pass signal. To show the time-domain signal, the frequency-domain power spectrum (2 to 200 Hz in steps of 4 Hz), and the time–frequency-domain power spectrogram of the ripples (2 to 200 Hz in steps of 4 Hz), we extracted the raw LFP within ±3 s around each ripple. Fig. S3 shows various ripple characteristics including ripple rates, durations, frequencies, power spectra, and inter-ripple intervals.

To compare the putatively physiological ripples against an identical number of “surrogate ripples” (Fig. 2I; Fig. S3C), we selected a random time point within ±60 s of each putatively physiological ripple (excluding time periods with artifacts), which we denoted as the peak time of the corresponding surrogate ripple (*45*).

#### Intracranial EEG: delta-phase locking of hippocampal ripples

To investigate whether ripples were locked to particular phases of low-frequency oscillations (*21, 33*), we filtered the hippocampal intracranial EEG with a two-pass finite impulse response (FIR) filter using MATLAB’s *designfilt* and *filtfilt* functions (filter order, 8000; lower cutoff frequency, 0.5 Hz; upper cutoff frequency, 2 Hz). We then estimated the phases of this delta-band filtered data using a Hilbert transform. For each ripple, we extracted its corresponding delta phase at the ripple peak time, and averaged the ripple-locked delta phases afterwards. Across channels, we tested whether the average delta phases were clustered using a Rayleigh test (*67*).

To assess statistical significance of ripple-phase coupling, we compared the empirical Rayleigh *z* value against 1001 surrogate Rayleigh *z* values, which we generated by performing the same steps as described above with the only difference that the inter-ripple intervals were randomly shuffled. We computed the *P*-value of the empirical Rayleigh *z* value in comparison to the surrogate Rayleigh *z* values as *P* = 1 − *rank*, where *rank* is the fraction of surrogate Rayleigh *z* values that were smaller than the empirical Rayleigh *z* value. This analysis showed that the delta-phase locking of empirical ripples was significantly stronger than in surrogate ripples with shuffled inter-ripple intervals (*P* = 0.017). To also test whether the empirical average delta phases (one per channel) were different from the surrogate average delta phases (one per channel in each of 1001 surrogate rounds), we performed a two-sample Kuiper’s test (*67*). This showed that the empirical preferred delta phases were significantly different from the preferred delta phases of surrogate ripples (Kuiper’s test: *k* = 1132988.000, *P* = 0.001).

#### Intracranial EEG: extrahippocampal ripple detection

To characterize the MTL-wide effects of hippocampal ripples, we detected ripples on bipolar channels from the amygdala, entorhinal cortex, parahippocampal cortex, and temporal pole using the same procedure as for hippocampal ripples (see above).

We then performed cross-correlations between the ripple time series of a given hippocampal channel and the ripple time series of another channel (where each ripple time series is a vector of zeros and ones, with values of 1 indicating ripple periods). We considered maximum time lags of ±5 s between both time series and used unbiased estimations of the cross correlations by means of MATLAB’s *xcorr* function. We smoothed the pairwise cross-correlations with a Gaussian filter (kernel length of 0.2 s) and *z*-scored the cross-correlations across time lags. To evaluate whether the *z*-scored cross-correlations were significantly positive, we then performed a cluster-based permutation test against 0 across channels (1001 surrogates).

In this cluster-based permutation test, we first applied a one-sample *t*-test to the empirical data, separately for each time lag, and identified contiguous clusters of time lags in which the uncorrected *P*-value of the *t*-test was significant (α = 0.05) and the *t*-value was positive. For each cluster, we computed an empirical cluster statistic by summing up all *t*-values that were part of that cluster (*t*_cluster-empirical_). We then compared the empirical cluster statistics against surrogate cluster statistics, which we obtained by inverting the sign of the cross-correlation values of a random subset of the cross-correlation series (*69*), performing exactly the same steps as described above for the empirical data, and keeping only the maximum cluster statistic (resulting in 1001 *t*_max-cluster-surrogate_ values). We considered an empirical cluster statistic (*t*_cluster-empirical_) significant if it exceeded the 95^th^ percentile of all surrogate maximum cluster statistics (*t*_max-cluster-surrogate_).

#### Intracranial EEG: LFP power during hippocampal ripples

To characterize the MTL-wide effects of hippocampal ripples, we computed ripple-aligned time– frequency-resolved power spectrograms in different MTL regions. Across the 41 sessions with macroelectrode recordings, 400 combinations of electrodes in the left/right hippocampus and electrodes in the left/right amygdala, left/right entorhinal cortex, left/right hippocampus, left/right parahippocampal cortex, or left/right temporal pole were available (240 in ipsilateral and 160 in contralateral hemispheres). 45 macroelectrode channels were located in the left amygdala (26 ipsilateral to their co-recorded hippocampal channel), 48 in the right amygdala (29 ipsilateral), 11 in the left entorhinal cortex (7 ipsilateral), 33 in the right entorhinal cortex (21 ipsilateral), 50 in the left hippocampus (29 ipsilateral), 54 in the right hippocampus (33 ipsilateral), 33 in the left parahippocampal cortex (18 ipsilateral), 31 in the right parahippocampal cortex (20 ipsilateral), 45 in the left temporal pole (26 ipsilateral), and 50 in the right temporal pole (31 ipsilateral).

For each macroelectrode channel, we computed the time–frequency spectrogram across the entire recording (using Morlet wavelets with 7 cycles at 50 logarithmically spaced frequencies between 1 and 200 Hz). Power values at time points with IEDs were excluded (i.e., set to NaN). Power values were then *z*-scored across time (using MATLAB’s *normalize* function), separately for each frequency. Around each hippocampal ripple (±3 s), we extracted the *z*-scored power values, averaged across ripples in each channel, and smoothed the average spectrograms with a Gaussian filter across time (kernel length, 0.2 s). Spectrograms were then truncated to ±0.5 s around the ripple peak time point. Next, we averaged the *z*-scored power spectrograms across channels for depiction and performed cluster-based permutation tests (1001 surrogates) across channels to statistically evaluate whether hippocampal ripples were associated with significant changes in LFP power in other MTL regions.

In these cluster-based permutation tests, we first applied a one-sample *t*-test to the empirical data, separately for each time–frequency bin, and identified contiguous clusters of time–frequency bins in which the uncorrected *P*-value of the *t*-test was significant (α = 0.025). For each cluster, we computed an empirical cluster statistic by summing up all *t*-values being part of that cluster (*t*_cluster-empirical_). We then compared the empirical cluster statistics against surrogate cluster statistics, which we obtained by flipping the sign of the power values of a random subset of the power spectrograms, performing exactly the same steps as described above for the empirical data, and keeping only the maximum cluster statistic (resulting in 1001 *t*_max-cluster-surrogate_ values). We considered an empirical cluster statistic (*t*_cluster-empirical_) significant if it exceeded the 97.5^th^ percentile or if it fell below the 2.5^th^ percentile of all surrogate maximum cluster statistics (*t*_max-cluster-surrogate_). We used a first-level alpha of α = 0.025 (for identifying significant time–frequency bins in the power spectrograms) and a second-level alpha of α = 0.025 (for identifying significant clusters of significant time–frequency bins), because the cluster-based permutation tests tested for both increases and decreases in power.

#### Intracranial EEG: relationship between hippocampal ripples and behavior

Given previous findings on associations between hippocampal ripples and human behavior (*36–41*), we asked whether hippocampal ripples were modulated by the subjects’ behavior in our associative object–location memory task. To test whether ripple characteristics varied as a function of trial period (i.e., ITI, cue, retrieval, feedback, and re-encoding), we estimated their rate, duration, and frequency in each trial phase. We then tested for significant associations between ripple characteristics and behavior using repeated measures ANOVAs (dependent variable: ripple characteristic, averaged across trials; independent variable: trial phase; Tukey-Kramer correction for multiple comparisons) and linear mixed models (Tables S4 and S5).

To test whether ripple rates were correlated with memory performance, we computed a partial correlation between trial-wise ripple rate and memory performance for each channel, separately for each trial phase (controlling for trial index and the interaction between memory performance and trial index). Across channels, we then tested whether the correlation values were significantly different from 0 using one-sample *t*-tests including Bonferroni correction for the number of trial phases. We also tested whether ripple rates were associated with trial index (i.e., time within the task) using partial correlations as described above for memory performance (controlling for memory performance and the interaction between memory performance and trial index). To corroborate the results from the partial correlations, we also performed linear mixed models (Tables S4 and S5).

To describe the time course of ripple rates during the trial phases, we computed peristimulus time histograms for the occurrence of ripples relative to the start and end time of the different trial phases (i.e., relative to the start for the cue and the feedback periods; relative to the end for the ITI, retrieval, and re-encoding periods). For each hippocampal ripple channel, we computed the time course of instantaneous ripple rates during trials with good memory performance and bad memory performance (based on a median split of each subject’s memory performance values) and used two-sided cluster-based permutation tests in Fieldtrip to evaluate whether time-resolved ripple rates changed during particular periods of the different trial phases between trials with good versus bad memory performance. Although the ITI, retrieval, and re-encoding periods had variable durations across trials, we restricted our analysis of their time-resolved ripple rates to −3 to +1 s relative to the end of these periods to examine ripple rates as a function of absolute time. We obtained very similar results when we examined ripple rates in normalized time, where we split each period into a fixed number of time bins (data not shown).

#### Single-neuron recordings: spike detection and sorting

Neuronal spikes were detected and sorted using Wave_Clus 3 (*71*). We used default settings with the following exceptions (*9*): “template_sdnum” was set to 1.5 to assign unsorted spikes to clusters in a more conservative manner; “min_clus” was set to 60 and “max_clus” was set to 10 in order to avoid over-clustering; and “mintemp” was set to 0.05 to avoid under-clustering. All clusters were visually inspected and judged based on their spike shape and its variance, inter-spike interval (ISI) distribution, and the presence of a plausible refractory period. If necessary, clusters were manually adjusted or excluded. Spike waveforms are shown as density plots in all figures. Spike times were aligned to the macroelectrode time axis using the trigger timestamps in order to investigate the relationship of single-neuron activity to events in the macroelectrode and behavioral data.

In total, we identified *N* = 1063 clusters (also referred to as “units,” “neurons,” or “cells” throughout the manuscript) across 27 experimental sessions from 20 patients that had microelectrodes implanted. We localized the tips of the depth electrodes to brain regions based on post-implantation MRI scans to assign neurons recorded from the corresponding microelectrodes to these regions (e.g., Fig. S6A). We recorded *n* = 340 neurons from the amygdala, *n* = 214 neurons from the entorhinal cortex, *n* = 24 neurons from the fusiform gyrus, *n* = 213 neurons from the hippocampus, *n* = 2 neurons from the insula, *n* = 126 neurons from the parahippocampal cortex, *n* = 135 neurons from the temporal pole, and *n* = 9 from the visual cortex. Due to low numbers of neurons in fusiform gyrus, insula, and visual cortex, we excluded these regions from region-specific analyses. Fourteen microelectrode patients of this study were also part of a previous study (*9*).

For recording quality assessment (Fig. S6), we calculated the number of units recorded on each microelectrode (for all microelectrodes with at least one unit); the ISI refractoriness of each unit; the mean firing rate of each unit; and the waveform peak signal-to-noise ratio (SNR) of each unit (*9*). The ISI refractoriness was assessed as the percentage of ISIs with a duration of <3 ms. The waveform peak SNR was determined as: SNR = A_peak_/SD_noise_, where A_peak_ is the absolute amplitude of the peak of the mean waveform, and SD_noise_ is the standard deviation of the raw data trace (filtered between 300 and 3000 Hz).

#### Single-neuron recordings: neuronal activity during hippocampal ripples

We were interested in understanding how the firing rates of neurons in various MTL regions behaved during hippocampal ripples (Fig. 2M; Fig. S5D). Thus, in subjects with single-neuron recordings, we computed instantaneous firing rates across the entire experiment (smoothed with a Gaussian filter with a kernel length of 0.2 s) and *z*-scored the firing rates across time. We then extracted the smoothed and *z*-scored firing rates during ±3 s relative to the hippocampal ripple peak time points and averaged across ripples afterward (separately for each neuron–ripple channel combination).

To test whether neuronal firing rates were significantly elevated during hippocampal ripples, we performed cluster-based permutation tests (1001 surrogates). In these cluster-based permutation tests, we first performed a one-sample *t*-test against 0, separately for each time bin. We then identified contiguous clusters of time bins with uncorrected *P*-values of *P* < 0.05 and calculated the sum *t*-value for each cluster (*t*_cluster-empirical_). To create surrogate cluster statistics, we inverted the sign of a random subset of the neuronal firing rates 1001 times. Using each set of surrogate data, we performed exactly the same steps as described above (time bin-wise one-sample *t*-tests against 0; identification of contiguous clusters of time bins with uncorrected *P*-values of *P* < 0.05; calculation of the sum *t*-value for each cluster). In each surrogate round, we kept the maximum sum *t*-value (*t*_max-cluster-surrogate_). We then considered *t*_cluster-empirical_ significant if it exceeded the 95^th^ percentile of all *t*_max-cluster-surrogate_ values. We separately tested for significantly elevated neuronal firing rates during hippocampal ripples depending on the regions in which the neurons were located (Fig. S5D).

We also examined the ripple-associated firing rates in different trial periods, separately for trials with good or bad memory performance (Fig. S4E). We tested whether the ripple-locked firing rates were significantly different between trials with good versus bad memory performance using cluster-based permutation tests (1001 surrogates) in Fieldtrip (*68*), by calculating surrogate cluster statistics based on surrogate data that were created by randomly re-assigning ripple-locked firing rates to trials with good or bad memory performance.

#### Single-neuron recordings: object cells

We designed the analysis of object cells to identify neurons that exhibited significant firing-rate increases in response to one particular object during the cue period. Hence, each object cell had to fulfill two criteria: (1) The absolute average firing rates during cue periods with the cell’s “preferred” object had to be significantly higher than the absolute average firing rates during cue periods with the other, “unpreferred” objects; (2) the time-resolved firing rates during cue periods with the preferred object had to exhibit a significant cluster of time points in which the relative firing rates (relative to a 1-s baseline period immediately before the onset of the cue period) were significantly higher than during trials with the unpreferred objects. We used criterion 2 in addition to criterion 1 to ensure that the cell’s preferred object was associated with a circumscribed firing-rate increase during the cue period.

To evaluate criterion 1, we performed the following steps: We computed the average firing rate of the cell during each cue period; we designated the object with the highest associated grand-average firing rate as the “preferred object”; we performed a two-sample *t*-test between the average firing rates from cue periods with the preferred object versus the average firing rates from cue periods with the unpreferred objects (*t*_empirical_); we created 1001 surrogate statistics (*t*_surrogate_) by performing the previous steps on randomly shuffled average firing rates (breaking up the assignment between average firing rates and object identity); and we then considered a cell as fulfilling criterion 1, if *t*_empirical_ exceeded the 95^th^ percentile of *t*_surrogate_.

To evaluate criterion 2, we performed the following steps: We computed the time-resolved firing rates of the cell during each cue period (temporal resolution, 0.01 s; smoothing with a Gaussian filter with a kernel length of 0.5 s); we baseline-corrected the time-resolved firing rates relative to a 1-s baseline period (immediately preceding the onset of the cue period); and we then used a cluster-based permutation test in Fieldtrip (*68*) to examine whether there was a time window during the cue period in which the baseline-corrected, time-resolved firing rates were significantly higher during cue periods with the preferred object as compared to cue periods with the unpreferred objects (1001 surrogates; one-sided α = 0.05). We then labeled a neuron as an object cell if both criteria were fulfilled. The cells’ preferred objects are indicated by orange color in all object cell-related figures.

To characterize object cells in greater detail (Fig 3, B–E), we calculated the percentage of object cells in the different MTL regions. To further understand their tuning, we calculated the sum of all significant time windows across object cells and their average time-resolved firing rates in response to the preferred and unpreferred objects. To examine the temporal stability of their tuning, we estimated each object cell’s time-resolved firing rate in response to the cell’s preferred object in the first and second half of the data and then compared the two tuning curves using a Pearson correlation across time. We present the results regarding the tuning strength and temporal stability of object cells mainly for illustration purposes because these analyses are not fully independent from our procedure of identifying the object cells.

#### Single-neuron recordings: place cells

We designed the analysis of place cells to identify cells that exhibited significant firing-rate increases when the subject was at a particular location of the virtual environment. Similar to previous single-neuron studies in humans (*9, 12, 57, 72–75*), the firing-rate profiles of our human place cells were less specific than those of spatially tuned cells in rodent studies, which is why these cells are sometimes referred to as “place-like cells” (*9, 74*). Our analysis nevertheless ensured that our human place cells exhibited distinct place fields in the virtual environment in which the cells’ firing rates were significantly higher than in the other parts of the environment. The weaker spatial tuning of human place cells is presumably due to several different reasons including the fact that the epilepsy patients did not physically navigate but rather performed virtual navigation.

To identify place cells in our dataset, we first resampled the behavioral information about the subject’s (x/y)-position in the environment to a time resolution of 10 Hz (*9, 72*) and calculated the neuronal firing rate (Hz) in each 0.1-s time bin. We then estimated the average firing rate within each bin of a 25 x 25 grid overlaid onto the environment (edge length of 400 vu) and excluded areas of the environment that the subject traversed less than two times. Time periods in which the subject did not move or did not turn around for more than 2 s were excluded from this firing-rate map (in order to exclude periods when the subject was idle). The firing-rate map was then smoothed with a Gaussian kernel (kernel size, 5; standard deviation, 1.5; using MATLAB’s *fspecial* and *conv2* functions). Next, we thresholded the firing-rate map at the 75^th^ percentile of the firing-rate values and considered contiguous bins with firing rates above this threshold as candidate place fields. We kept the candidate place field with the highest sum firing rate as the potential place field of this cell. We quantified the strength of this potential place field as the *t*-statistic of a two-sample *t*-test comparing the firing rates when the subject was inside the place field with the firing rates when the subject was outside the place field. We considered the empirical *t*-statistic (*t*_empirical_) significant if it exceeded the 95^th^ percentile of 1001 surrogate *t*-statistics (*t*_surrogate_), which we obtained by performing exactly the same procedure as described above with the only difference that we circularly shifted the firing rates relative to the behavioral data by a random lag (with the end of the session wrapped to the beginning), following previous studies [e.g., (*9, 75*)]. If *t*_empirical_ was above the 95^th^ percentile of all *t*_surrogate_ values, we considered the place field significant and designated the cell as a place cell.

To characterize place cells (Fig. 3, H–L), we estimated the percentage of place cells in the different MTL regions. We also calculated the cumulative distribution of place fields across the virtual environment (relative to the cumulative distribution of the firing-rate maps across the environment) and the size of the place fields across all place cells (expressed as percentages relative to the spatial extent of the firing-rate maps). We furthermore estimated the average firing rate of place cells inside versus outside the place fields and quantified the temporal stability of the place cells by performing a Pearson correlation between the firing-rate map estimated using the first half of the data and the firing-rate map estimated using the second half of the data. We assessed the statistical significance of the temporal stability values by performing a one-sample *t*-test of the correlation values against 0. We present the results regarding the tuning strength and temporal stability of place cells mainly for illustration because these analyses are not fully independent from our procedure of identifying place cells.

#### Single-neuron recordings: conjunctive object–place cells

We defined conjunctive object–place cells as cells that exhibited significant place tuning only during trials with one particular object (but not during trials with any other object). To identify conjunctive object–place cells, we performed the place-cell analysis (as described above) eight times for each cell, each time only considering the trials with a particular object (for an example, see Fig. S7A). If a cell exhibited significant place tuning at a Bonferroni-corrected alpha-level of α = 0.05/8 for exactly one object, we considered the cell as a conjunctive object–place cell.

To characterize conjunctive cells (Fig. S7, B–D), we estimated the percentage of conjunctive cells in different MTL regions and estimated their overlap with object cells and place cells. To provide supplemental evidence for the fact that conjunctive cells exhibited spatial tuning that was specific to trials with one particular object—instead of demonstrating overall spatial tuning as in place cells— we estimated the pairwise similarity of the spatial firing-rate maps between trials with different objects using Pearson correlations (and averaged the similarity values across all pairwise comparisons afterward). We then performed one-sample *t*-tests against 0 to show that these similarity values were not significantly above 0 for conjunctive cells (in line with spatial tuning being specific to trials with a particular object) but significantly above 0 for place cells (in line with spatial tuning being stable across time and thus independent of particular objects).

#### Single-neuron recordings and intracranial EEG: coactivity of object cells and place cells during hippocampal ripples

Our main hypothesis pertained to the question whether object and place cells activated together during the same hippocampal ripples. To answer this question, we performed the following procedure.

For each neuron–ripple channel combination (*n* = 1,716), we first estimated whether the neuron was active in various time bins relative to the ripple peaks (301 time bins at −0.75 to 0.75 s relative to the ripple peaks with a bin width of 0.1 s and a step size of 0.005 s; 95% overlap between neighboring time bins). In total, there were 192 neuron–ripple channel combinations in which the neuron was an object cell, and there were 182 neuron–ripple channel combinations in which the neuron was a place cell.

Next, for each simultaneously recorded pair of an object and a place cell, we quantified the coactivity of both cells using a previously developed coactivity *z*-score (*76*). For a given combination of time bins used to estimate the activations of both cells, we calculated the coactivity *z*-score as:

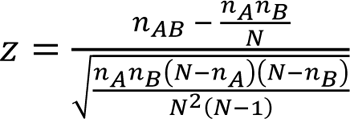

where *N* is the total number of ripples, *n_A_* is the number of ripples in which cell A spiked, *n_B_* is the number of ripples in which cell B spiked, and *n_AB_* is the number of ripples in which both cells spiked. An illustration of the coactivity score is shown in Fig. S8. Because we estimated the activity of each cell at various time points relative to the ripple peaks (see above), we were able to compute the coactivity *z*-score for various combinations of time bins (i.e., for all possible time bins *i*_A_ and *j*_B_ of cell A and cell B, respectively). By considering all possible time-bin combinations, this procedure resulted in a two-dimensional time-by-time coactivity map for each cell pair that showed the cells’ coactivity at various time points relative to the hippocampal ripple peaks. Example coactivity maps of individual cell pairs are shown in Fig. S12.

To statistically evaluate the coactivity maps across all associative object cell–place cell pairs, we used a series of cluster-based permutation tests. During retrieval, associative cell pairs were selected as cell pairs in which the subject’s response location in response to the object cell’s preferred object was inside the place field of the place cell. During re-encoding, associative cell pairs were selected as the pairs in which the correct location of the preferred object of the object cell was inside the place field of the place cell. We used the cluster-based permutation tests to compare the coactivity maps (±0.25 s around the ripple peaks) against chance (denoted as “> 0” or as “> surrogates” in the figure legends); against coactivity maps from a baseline period (−0.75 to −0.25 s before the ripple peaks and 0.25 to 0.75 s after the ripple peaks, averaged across these two time windows; denoted as “> baseline” in the figure legends); and against the coactivity maps of object cell–place cell pairs encoding non-associative information (denoted as “> pref. object & response location outside place field” or “> pref. object and object location outside place field” in the figure legends). We reasoned that together these three tests would provide robust information about the significance of ripple-locked coactivity of object and place cells.

The different cluster-based permutation tests are illustrated in Fig. S9. When using cluster-based permutation tests to compare the coactivity maps against chance, we proceeded as follows. We performed a one-sample *t*-test of the coactivity *z*-values against zero across all object cell–place cell– ripple channel combinations, separately for each bin of the two-dimensional time-by-time coactivity maps. We next identified clusters of contiguous bins, for which the uncorrected *P*-value was significant (α = 0.05) and the *t*-statistic was above zero (given our *a priori* hypothesis of increased object cell–place cell coactivity during hippocampal ripples). For each cluster, we then computed the sum of all *t*-values (*t*_cluster-empirical_) and compared these empirical cluster statistics against 1001 surrogate statistics. To obtain each of the surrogate statistics, we inverted the sign of the coactivity *z*-values of a random subset of the empirical coactivity maps. We then estimated the surrogate cluster statistics exactly as for the empirical data and kept the maximum surrogate cluster statistic (*t*_max-cluster-surrogate_; *n* = 1001). For each empirical cluster statistic (*t*_cluster-empirical_), we finally tested whether it exceeded the 95^th^ percentile of all maximum surrogate cluster statistics (*t*_max-cluster-surrogate_). If so, the empirical cluster was considered significant. We also performed a variant of this cluster-based permutation test against chance (Fig. S10), where we subtracted a surrogate coactivity map from each corresponding empirical coactivity map before calculating the statistics across cell pairs. The surrogate coactivity map of a given cell pair was estimated by circularly shifting the ripple-locked activity levels of the object cell relative to the ripple-locked activity levels of the place cell by a random latency before calculating the two-dimensional coactivity map. The statistics across associative cell pairs were then performed as described above for the comparison against zero.

When using cluster-based permutation tests to compare the coactivity maps of associative object cell– place cell–ripple channel combinations (set A) against baseline data or against the coactivity maps of non-associative object cell–place cell–ripple channel combinations (set B), we proceeded as follows. We performed a two-sample *t*-test between the coactivity *z*-values of the two sets of coactivity maps, separately for each bin of the two-dimensional time-by-time coactivity maps. We next identified clusters of contiguous bins, for which the uncorrected *P*-value was significant (α = 0.05) and the *t*-statistic was positive. For each cluster, we then computed the sum of all *t*-values (*t*_cluster-empirical_) and compared these empirical cluster statistics against 1001 surrogate statistics. To obtain each of the surrogate statistics, we swapped a random subset of the set A and set B coactivity maps (when comparing the coactivity maps against baseline coactivity maps) or randomly reassigned the coactivity maps to the two sets A and B (when comparing the coactivity maps against coactivity maps of non-associative cell pairs). We then estimated cluster statistics exactly as for the empirical data and kept the maximum cluster statistic (*t*_max-cluster-surrogate_). For each empirical cluster statistic (*t*_cluster-empirical_), we finally tested whether it exceeded the 95^th^ percentile of all maximum surrogate cluster statistics (*t*_max-cluster-surrogate_). If so, the empirical cluster was considered significant.

To further describe the data underlying the coactivity maps in Fig. 4, Fig. S11 shows the number of object cell–place cell–ripple channel combinations contributing to the coactivity maps (which can vary between the bins in the coactivity maps); the total number of ripples underlying the coactivity maps (which can also vary between the different bins in the coactivity maps); the individual coactivity *z*-scores underlying the maxima in the coactivity maps; and the brain regions of the object cells and place cells contributing to the maxima in the coactivity maps. To validate using *z*-scores to compute coactivations (Fig. 4), we also computed the coactivations using Pearson correlations (between the activity vectors of object cells and place cells) and obtained qualitatively identical results (Fig. S13).

In our main results (Fig. 4, C–H), we separately considered early and late ripples (i.e., the first and the second half of all ripples, respectively). To understand whether this distinction mainly followed a distinction between ripples occurring before the initial formation and ripples occurring after the initial formation of associative memories, we performed a separate analysis in which we estimated the trial in which the subject exhibited the strongest improvement in memory performance, separately for each object. We identified the object-specific trial of strongest memory improvement by (1) estimating the memory performance on each trial; (2) smoothing these performance values with a running average of three trials (to attenuate the effect of potential outliers); (3) iteratively computing a two-sample *t*-test between the memory-performance values from trials (*i*+1):*n* and those from trials 1:*i*, where *i* is the current trial index and *n* is the total number of trials; (4) smoothing the resulting *t*-statistics with a running average of three trials (again to attenuate the effect of potential outliers); and by (5) identifying the trial with the largest *t*-statistic (where the *t*-statistics of the first and last trial were not considered in order to exclude the possibility that they were selected as the trial with the largest *t*-statistic). We considered all ripples occurring before or in the trial with the largest *t*-statistic as ripples occurring “before initial memory formation.” All other ripples (occurring after the trial with the largest *t*-statistic) were considered as ripples occurring “after initial memory formation.” The results for ripples occurring before initial memory formation and those occurring after initial memory formation were very similar to the results for early versus late ripples (Fig. S15), suggesting that the significant object cell–place cell–coactivations during late ripples were at least partly dependent on an initial formation of the associative memories.

To understand whether the key coactivity results (Fig. 4, E and H) were driven by ripples occurring during movement or during non-movement, we categorized each time point of the task according to whether or not the subject was moving, and analyzed the data separately for both conditions. We observed that the cellular coactivations were mainly driven by ripples occurring during non-movement periods (Fig. S16).

### Supplementary Text

#### Supplementary Text S1: Epileptic activity, ripples, and behavior

Between epileptic seizures, epilepsy patients exhibit interictal epileptiform discharges (IEDs), which are pathological bursts of neuronal activity. IEDs are readily visible in intracranial EEG recordings based on their high amplitudes, sharp amplitude changes, and power increases across a broad frequency range (Fig. S2A) (*34, 77*).

IEDs complicate the detection of ripples in human epilepsy patients. Following previously established procedures for the automated detection of IEDs (*34, 39, 45*), we therefore identified IEDs in our dataset and conservatively excluded 16.591 ± 2.692% (mean ± SEM; *n* = 62 channels) of the data because of IEDs. We additionally excluded a small amount of the data due to ripple-like phenomena in the grand-average signal, which are presumably muscle or other artifacts, leading to 17.380 ± 2.668% (mean ± SEM; *n* = 62 channels) of the data being excluded before ripple detection. As expected, we observed that a higher prevalence of IEDs was correlated with lower ripple rates across channels, both when computing ripple rates relative to the entire data and relative to data periods without artifacts (Spearman’s *rho* = −0.740, *P* < 0.001 and Spearman’s *rho* = −0.451, *P* < 0.001, respectively; *n* = 62 channels), potentially indicating that the detected ripples were indeed physiologic and that IEDs lead to a reduction of such physiological ripples (*41*).

Previous studies showed that IEDs lead to transitory cognitive impairments and that they impede the encoding and retrieval of associative memories (*41, 78*). We therefore analyzed the relationship between IEDs and behavior in our associative object–location memory task and found that the prevalence of IEDs in a given trial was modulated by trial phase [repeated measures ANOVA: *F*(4, 244) = 3.705, *P* = 0.006; Fig. S2C]. Specifically, IEDs were less prevalent during cue periods as compared to ITI and retrieval periods (post-hoc comparisons: *P*_Tukey-Kramer_ = 0.012 and *P*_Tukey-Kramer_ < 0.001, respectively). We did not observe significant relationships between IED prevalence and memory performance (Fig. S2D), which replicates a previous report with a similar paradigm and which may be due to the self-paced nature of this task (*79*). Instead, we found that IED prevalence slightly increased with time, which was most clearly visible for the ITI period [one-sample *t*-test against 0: *t*(60) = 2.985, *P* = 0.021, Bonferroni corrected for five tests] and the feedback period [one-sample *t*-test against 0: *t*(57) = 3.339, *P* = 0.007, Bonferroni corrected for five tests; Fig. S2E], but also irrespective of trial phase [one-sample *t*-test of channel-wise correlation coefficients between trial index and IED prevalence versus 0: *t*(61) = 2.207, *P* = 0.031; Fig. S2F].

We furthermore used a linear mixed model to investigate the relationship between IEDs and behavior. This linear mixed model used IED prevalence in a given trial phase as dependent variable and trial-wise memory performance, trial index, and trial phase as fixed effects (Table S3). Channel index was included as a random effect. In line with the above-mentioned results, the linear mixed model showed that memory performance was not related to IED prevalence and that trial index was positively correlated with IED prevalence. As compared to the ITI period, the cue period and the feedback period showed a lower prevalence of IEDs. Together, these results indicate that IEDs are modulated by our associative object–location memory task, but that they are not directly related to memory performance in this task.

#### Supplementary Text S2: Hippocampal ripples and behavior

Previous studies demonstrated that characteristics of hippocampal ripples can vary between different cognitive tasks, between different components of the same task, and as a function of the subjects’ behavioral performance (*33, 36–41, 50*). Hence, we aimed at understanding whether hippocampal ripples were related to the subjects’ behavioral state and memory performance in our object–location memory task.

We first tested whether ripple characteristics (rate, duration, and frequency) varied between the different phases of each trial (ITI, cue, retrieval, feedback, and re-encoding; Fig. 1B). In line with previous results (*36, 39*), ripple rates were increased during the ITI and cue periods, when subjects rested and viewed pictures of the objects that cued them to remember particular locations in the environment, respectively [repeated measures ANOVA: *F*(4, 244) = 19.942, *P* < 0.001; post-hoc comparisons between ITI and retrieval, feedback, or re-encoding: all *P*_Tukey-Kramer_ < 0.024; post-hoc comparisons between cue and retrieval, feedback, or re-encoding: all *P*_Tukey-Kramer_ < 0.001; Fig. S4A]. Ripple durations showed a similar modulation pattern [repeated measures ANOVA: *F*(4, 236) = 2.463, *P* = 0.046; Fig. S4A], but post-hoc comparisons were unsignificant (all *P*_Tukey-Kramer_ > 0.138). Ripple frequency was not modulated by trial phase [repeated measures ANOVA: *F*(4, 236) = 0.562, *P* = 0.690; Fig. S4A]. These results demonstrate that the rate of human hippocampal ripples changes with the subjects’ behavioral state in our associative object–location memory task.

We next examined the memory relevance and temporal stability of hippocampal ripple rates. To this end, we estimated across-trial correlations between ripple rates and memory performance or trial index, separately for each channel, and tested for a consistent relationship across channels. During the cue period, ripple rates correlated positively with memory performance, i.e., with more frequent ripples predicting better memory performance in the subsequent retrieval period [one-sample *t*-test against 0: *t*(60) = 2.763, *P* = 0.038, Bonferroni corrected for five tests; Fig. S4B]. During re-encoding, ripple rates correlated negatively with memory performance meaning that higher ripple rates followed memory responses with lower memory performance [one-sample *t*-test against 0: *t*(60) = −4.181, *P* < 0.001, Bonferroni corrected for five tests; Fig. S4B]. This result implicates hippocampal ripples in the formation, or updating, of associative memories as subjects corrected their memories by viewing the objects in their true locations during the re-encoding periods.

Exploratory time-resolved analyses showed that during the cue period the difference in ripple rates between good and bad trials (defined by a median split of each subject’s memory performance values) was strongest at 1.162–1.470 s after cue onset (cluster-based permutation test: *t*_cluster_ = 1023.281, *P* = 0.011), whereas the difference during re-encoding was broadly distributed over time (Fig. S4C). Ripple rates did not correlate with memory performance during the retrieval period and ripple rates did not increase at a fixed interval prior to successful retrieval (Fig. S4, B–C), which is presumably a result of the self-paced nature of the task. When we investigated the relationship between ripple rates and trial index, we found that ripple rates were largely stable across the course of the experiment, with a decrease of ripples over time during the feedback period [one-sample *t*-test against 0: *t*(59) = −3.178, *P* = 0.012, Bonferroni corrected for five tests; Fig. S4D].

To corroborate these results (Fig. S4), we performed follow-up analyses using linear mixed models where we also controlled for the trial-wise prevalence of artifacts and obtained qualitatively identical results (Tables S4 and S5). In these linear mixed models, ripple rates were entered as the dependent variable and memory performance, trial index, and trial phase were modeled as fixed effects. We included channel index as a random effect. In the second linear mixed model (Table S5), the trial- and phase-wise prevalence of artifacts (i.e., IEDs and ripple-like events in the grand-average signal) was included as a covariate. Both linear mixed models showed that ripple rates were higher during the cue period and lower during the retrieval, feedback, and re-encoding periods as compared to the ITI period. As compared to the ITI period, ripple rates and memory performances were more positively correlated with each other during the cue period and more negatively correlated with each other during the re-encoding period. Furthermore, ripple rates during the cue period and the feedback period decreased with increasing trial index.

These findings demonstrate that the rate of human hippocampal ripples is associated with the subjects’ behavioral state and memory performance in our associative object–location memory task. Increased ripple rates during the cue period preceded the successful retrieval of associative memories, which implicates hippocampal ripples in retrieval processes. Increased ripple rates during the re-encoding period followed the unsuccessful retrieval of associative memories, suggesting an additional role for hippocampal ripples in establishing or updating associative memories. These observations thus extend the previously established links between hippocampal ripples and memory processes in awake humans (*33, 36–41, 50*).

#### Supplementary Text S3: Ripple-locked neuronal firing and behavior

Our analyses in the main text showed that hippocampal ripples were associated with a state of increased neural activations across the human MTL (Fig. 2, J–M), including a general increase of single-neuron firing during hippocampal ripples (Fig. 2M). To further understand the functional relevance of this recruitment of neuronal activity during hippocampal ripples, we asked whether ripple-related single-neuron firing would differ as a function of trial phase and the subjects’ memory performance (using a median split of each subject’s memory performance values across trials).

We found that ripple-locked firing rates during re-encoding periods following inferior memory responses were significantly increased as compared to re-encoding periods after better memory responses (cluster-based permutation test: *t*_cluster1_ = −357.511, *P* = 0.002, 0.113–0.385 s relative to the ripple peaks; *t*_cluster2_ = −206.887, *P* = 0.010, −0.115–0.033 s; *n* = 1716 neuron–ripple-channel combinations; Fig. S4E). In the other trial phases (ITI, cue, retrieval, and feedback), ripple-locked firing rates exhibited similar increases during trials with better as compared to inferior memory performance (Fig. S4E). Together with the increased ripple rates during re-encoding periods following inferior memory retrieval (Fig. S4B), this elevated single-neuron firing supports the idea that neuronal resources are broadly recruited in situations following inferior memory performance— potentially to newly establish or update existing memory representations.

#### Supplementary Text S4: Supplementary discussion

##### Associative memory and the lateral entorhinal cortex

In the main text we discussed two broader hypotheses on the possible neural implementation of associative memories (the “conjunctive hypothesis” and the “coactivity hypothesis”). Both hypotheses align with the common view that the hippocampus holds a crucial function in binding the separate elements of associative memories together (*23, 80, 81*).

In addition to these two hypotheses, associative memory presumably rests upon multiple other neural mechanisms as well. In particular, several recent studies indicate that the lateral entorhinal cortex holds a crucial function in associative memory, as rats with lesions to the lateral entorhinal cortex are unable to recognize object–context associations (*82*). Furthermore, over the course of associative learning, oscillatory coupling at frequencies between 20–40 Hz evolves between the lateral entorhinal cortex and the dorsal hippocampus (*83*), suggesting that interregional (entorhinal–hippocampal) neural communication contributes to associative memory (*84, 85*). Specifically, fan cells of the lateral entorhinal cortex may be important for associative memory as inhibiting them optogenetically impairs the learning of new associative memories (*86, 87*). These studies provide strong evidence that cells and oscillations in the lateral entorhinal cortex are key components of the neural substrate of associative memories. Some of our object and place cells may have indeed been located in the lateral entorhinal cortex and this brain region may thus have significantly contributed to our coactivity results.

##### Neural changes during hippocampal ripples

We found that hippocampal ripples did not only coincide with highly specific coactivations of object cells and place cells, but that they were also associated with broad increases in neural activity across the human MTL (Fig. 2; Fig. S4E; Fig. S5). Specifically, we observed that hippocampal ripples occurred simultaneously with ripples in extrahippocampal MTL regions; that they were associated with elevated high-frequency power and decreased low-frequency power in these regions; and that they were coupled to increased single-neuron spiking across the human MTL. Such brain states of increased excitation may provide a basis for establishing and activating connections between previously unconnected neurons. Hippocampal ripples may thus support various cognitive functions that rely on interactions between separate and widespread neural representations.

Our finding of wide-ranging neural changes during hippocampal ripples aligns with previous reports in both animals and humans. In rodents, hippocampal sharp-wave ripples are accompanied by widespread cortical and subcortical activations (*22, 88, 89*) and coincide with transient brain-wide increases in functional connectivity (*24*). In humans, hippocampal ripples are coupled with ripples in many cortical areas (*23*, *33*, *37*) and are accompanied by increased high-frequency broadband activity in widely distributed neocortical sites (*40*). They coincide with memory-specific high-frequency broadband activity in high-order visual areas during memory retrieval (*36*) and support the reinstatement of memory-specific single-neuron sequences in the temporal cortex (*38*). These studies support our conclusion that ripples in the hippocampus provide a time window for increased activation and excitation across the brain, which may support associative memory by establishing and reactivating connections between separate neural elements.

##### Ripples in the anterior hippocampus versus ripples in the posterior hippocampus

We recorded ripples solely from the anterior human hippocampus because this was the part of the hippocampus with the most frequent electrode coverage, given that the patients’ implantation schemes were determined by clinical needs. The human anterior hippocampus corresponds to the rodent ventral hippocampus, which poses a challenge when trying to directly compare our results to studies on hippocampal sharp-wave ripples in rodents, which typically record sharp-wave ripples from the dorsal hippocampus.

Prior rodent studies showed that ripples in the ventral segment of the hippocampus are often isolated from ripples in the intermediate and dorsal segments of the hippocampus (*76, 90*). Accordingly, ripples in different parts of the hippocampus may play different functional roles and may process different types of information. Indeed, it has been shown that sharp-wave ripples in the dorsal hippocampus of the rat are strongly enhanced by novel and rewarding experiences, whereas ventral hippocampal sharp-wave ripples are not modulated by novelty and reward (*76*). Concordantly, ripples in the dorsal and ventral hippocampus activate distinct and opposing patterns of spiking activity in the nucleus accumbens, whereby only those neurons of the nucleus accumbens are tuned to task- and reward-related information whose activity is coupled to ripples in the dorsal hippocampus (*76*). These results point at differences between sharp-wave ripples in the dorsal versus ventral rodent hippocampus. Building on these findings in rodents, future studies in humans should thus investigate whether ripples in the anterior versus posterior human hippocampus have similar or different physiological and functional properties.

##### Ripples versus gamma and epsilon oscillations

It is of ongoing debate whether the periods of elevated power around 80–140 Hz in this and other human studies [e.g., (*33, 36–41, 50*)] are the human homolog of ripples in rodents (*15*). Furthermore, whereas the frequency band of ripples in rodents (about 150–250 Hz) is different from the frequency bands of gamma (about 30–90 Hz) and epsilon (about 90–150 Hz) oscillations (*46*), the ripple band in human studies (about 80–140 Hz) overlaps with the frequency ranges of gamma and epsilon oscillations. It could thus be the case that the transient high-frequency events that we referred to as ripples are actually pronounced gamma or epsilon oscillations.

Furthermore, if human ripples indeed occur at frequencies of about 80–140 Hz, currently employed ripple-detection algorithms in human studies may detect a mixture of human ripples, gamma oscillations, and epsilon oscillations. For example, the ripples that we detected during the cue period and for which we found a positive relationship to memory performance in the subsequent retrieval period (Fig. S4B) might also constitute increased power of gamma or epsilon oscillations (elicited by the visual presentation of the objects), for which a positive relationship with successful recall has been demonstrated before (*91–95*).

In addition, it is still unclear how physiological ripples can exactly be distinguished from pathological ripples in iEEG recordings of human epilepsy patients (*42*). In this study, we used a conservative rejection of IEDs and required ripple candidates to pass several criteria in order to increase the likelihood of only including physiological ripples in our analyses. Nevertheless, despite our best efforts these putatively physiological ripples may still have contained some pathological ripples. Future studies may identify more precise markers to differentiate between physiological and pathological ripples and thus enable the investigation of strictly physiological ripples in human epilepsy patients.

## Supplementary Tables

**Table S1.**
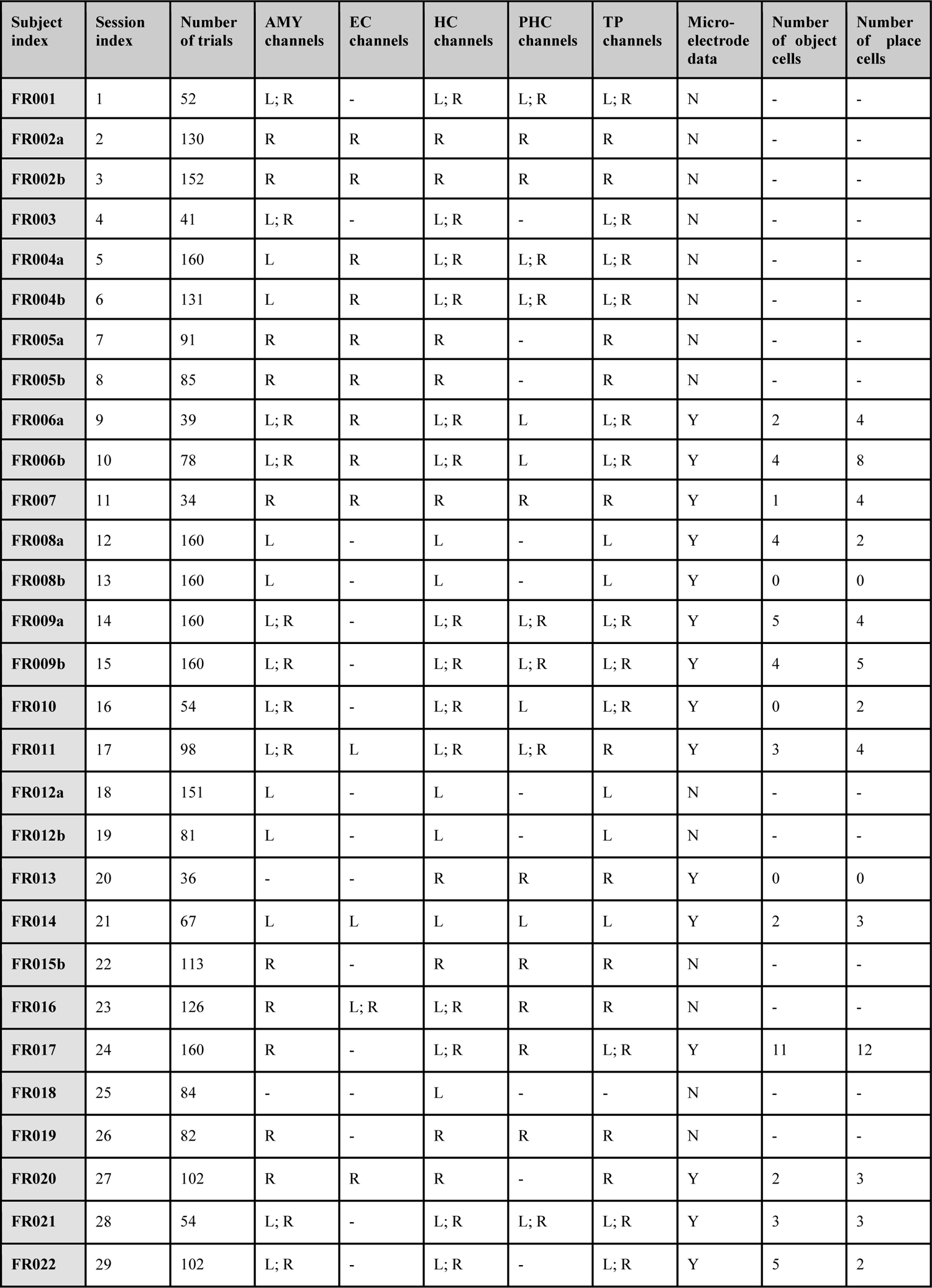
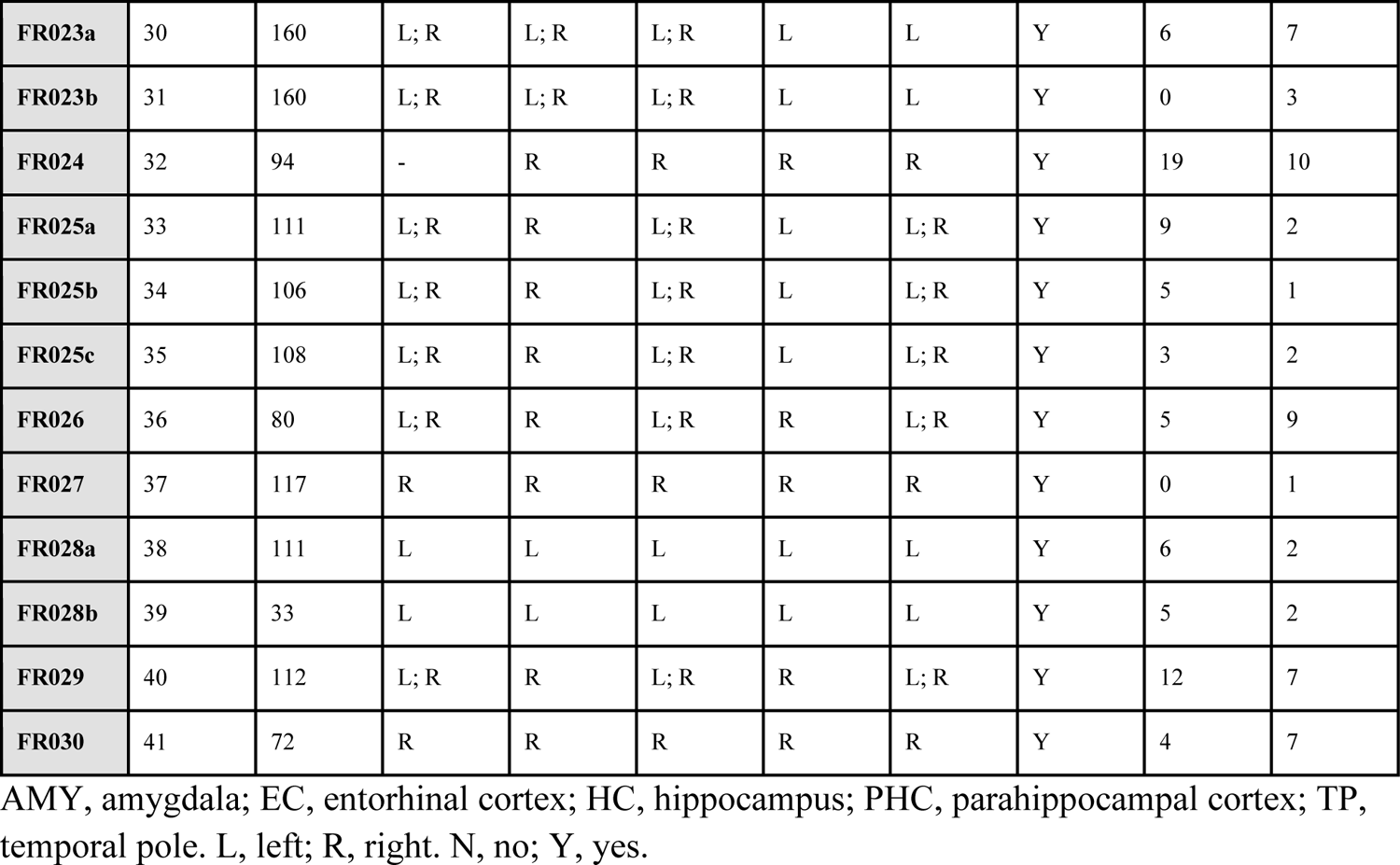
Subject information.

| Subject index | Session index | Number of trials | AMY channels | EC channels | HC channels | PHC channels | TP channels | Micro-electrode data | Number of object cells | Number of place cells |
| --- | --- | --- | --- | --- | --- | --- | --- | --- | --- | --- |
| FR001 | 1 | 52 | L; R | - | L; R | L; R | L; R | N | - | - |
| FR002a | 2 | 130 | R | R | R | R | R | N | - | - |
| FR002b | 3 | 152 | R | R | R | R | R | N | - | - |
| FR003 | 4 | 41 | L; R | - | L; R | - | L; R | N | - | - |
| FR004a | 5 | 160 | L | R | L; R | L; R | L; R | N | - | - |
| FR004b | 6 | 131 | L | R | L; R | L; R | L; R | N | - | - |
| FR005a | 7 | 91 | R | R | R | - | R | N | - | - |
| FR005b | 8 | 85 | R | R | R | - | R | N | - | - |
| FR006a | 9 | 39 | L; R | R | L; R | L | L; R | Y | 2 | 4 |
| FR006b | 10 | 78 | L; R | R | L; R | L | L; R | Y | 4 | 8 |
| FR007 | 11 | 34 | R | R | R | R | R | Y | 1 | 4 |
| FR008a | 12 | 160 | L | - | L | - | L | Y | 4 | 2 |
| FR008b | 13 | 160 | L | - | L | - | L | Y | 0 | 0 |
| FR009a | 14 | 160 | L; R | - | L; R | L; R | L; R | Y | 5 | 4 |
| FR009b | 15 | 160 | L; R | - | L; R | L; R | L; R | Y | 4 | 5 |
| FR010 | 16 | 54 | L; R | - | L; R | L | L; R | Y | 0 | 2 |
| FR011 | 17 | 98 | L; R | L | L; R | L; R | R | Y | 3 | 4 |
| FR012a | 18 | 151 | L | - | L | - | L | N | - | - |
| FR012b | 19 | 81 | L | - | L | - | L | N | - | - |
| FR013 | 20 | 36 | - | - | R | R | R | Y | 0 | 0 |
| FR014 | 21 | 67 | L | L | L | L | L | Y | 2 | 3 |
| FR015b | 22 | 113 | R | - | R | R | R | N | - | - |
| FR016 | 23 | 126 | R | L; R | L; R | R | R | N | - | - |
| FR017 | 24 | 160 | R | - | L; R | R | L; R | Y | 11 | 12 |
| FR018 | 25 | 84 | - | - | L | - | - | N | - | - |
| FR019 | 26 | 82 | R | - | R | R | R | N | - | - |
| FR020 | 27 | 102 | R | R | R | - | R | Y | 2 | 3 |
| FR021 | 28 | 54 | L; R | - | L; R | L; R | L; R | Y | 3 | 3 |
| FR022 | 29 | 102 | L; R | - | L; R | - | L; R | Y | 5 | 2 |
| **FR023a** | 30 | 160 | L; R | L; R | L; R | L | L | Y | 6 | 7 |
| **FR023b** | 31 | 160 | L; R | L; R | L; R | L | L | Y | 0 | 3 |
| **FR024** | 32 | 94 | - | R | R | R | R | Y | 19 | 10 |
| **FR025a** | 33 | 111 | L; R | R | L; R | L | L; R | Y | 9 | 2 |
| **FR025b** | 34 | 106 | L; R | R | L; R | L | L; R | Y | 5 | 1 |
| **FR025c** | 35 | 108 | L; R | R | L; R | L | L; R | Y | 3 | 2 |
| **FR026** | 36 | 80 | L; R | R | L; R | R | L; R | Y | 5 | 9 |
| **FR027** | 37 | 117 | R | R | R | R | R | Y | 0 | 1 |
| **FR028a** | 38 | 111 | L | L | L | L | L | Y | 6 | 2 |
| **FR028b** | 39 | 33 | L | L | L | L | L | Y | 5 | 2 |
| **FR029** | 40 | 112 | L; R | R | L; R | R | L; R | Y | 12 | 7 |
| **FR030** | 41 | 72 | R | R | R | R | R | Y | 4 | 7 |

AMY, amygdala; EC, entorhinal cortex; HC, hippocampus; PHC, parahippocampal cortex; TP, temporal pole. L, left; R, right. N, no; Y, yes.

**Table S2.**
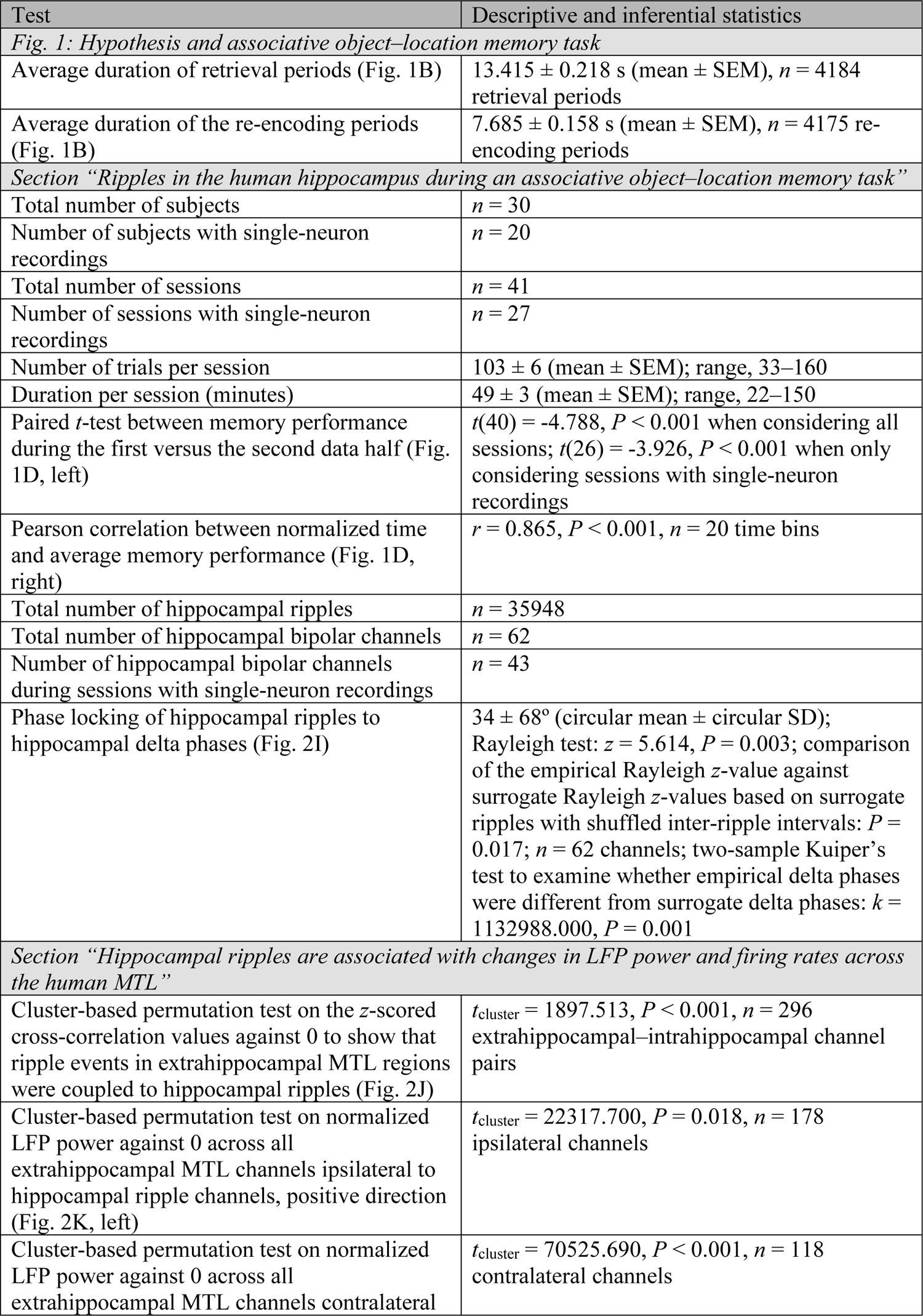
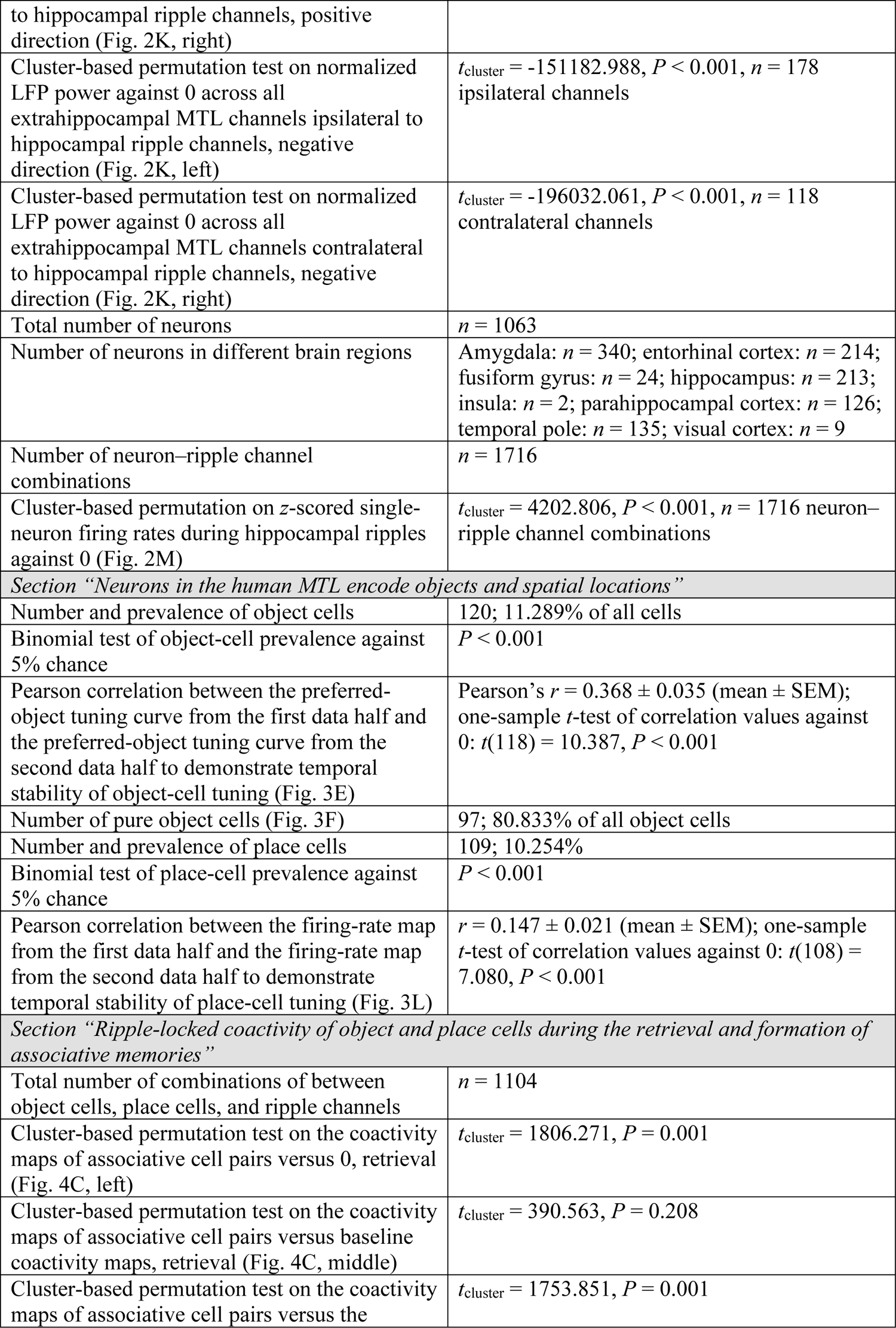
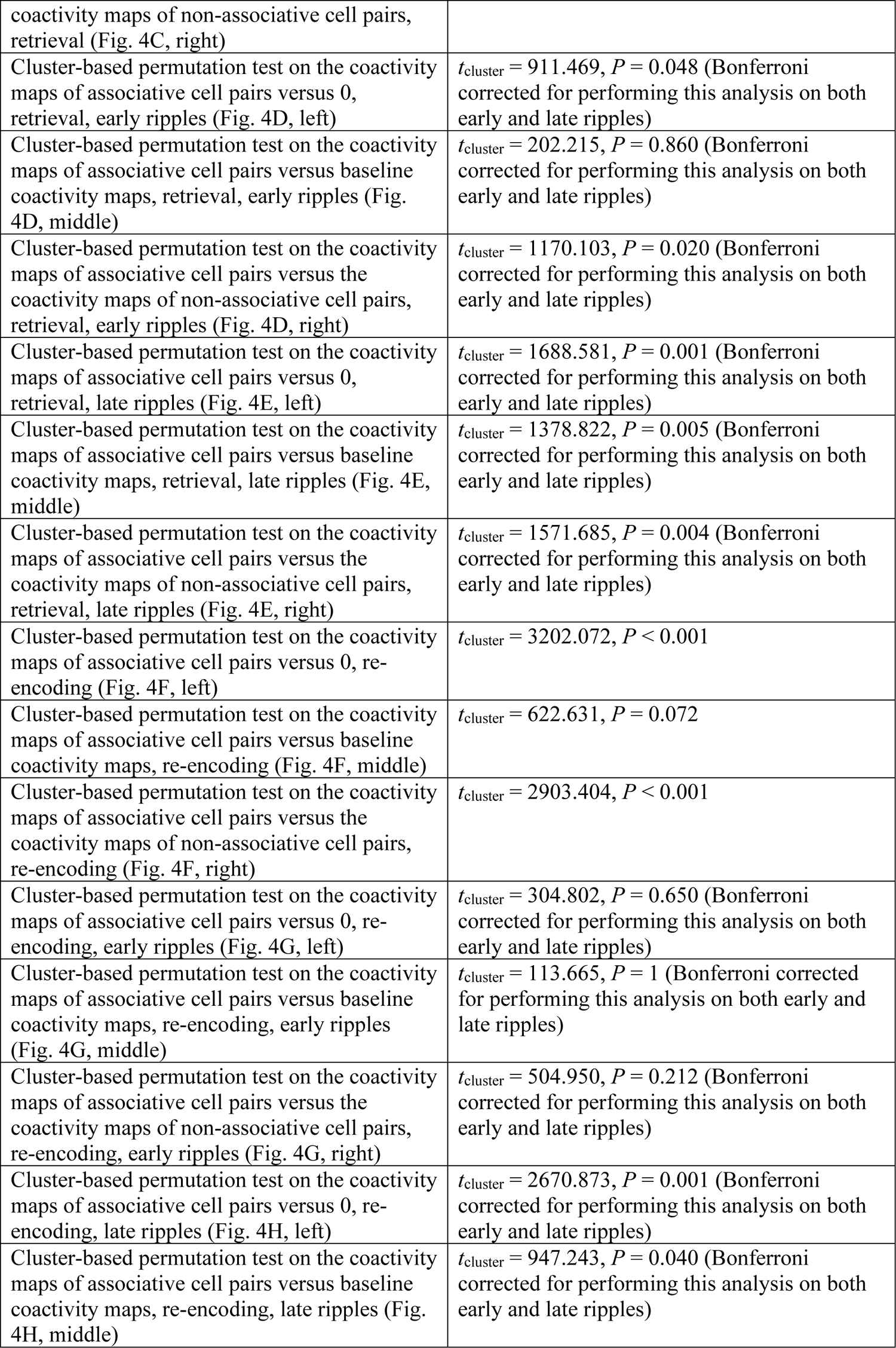
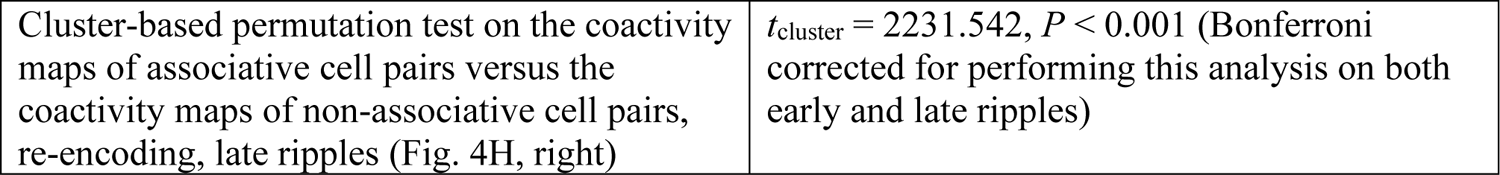
Detailed statistical information for analyses in the main text.

**Table S3.** Linear mixed model to analyze the prevalence of interictal epileptic discharges (IEDs; dependent variable) as a function of behavior.

| Predictor | *t* | DF | *P* |
| --- | --- | --- | --- |
| Memory performance | -0.031 | 32133 | 0.975 |
| Trial index | 2.398 | 32133 | 0.017* |
| Cue | -5.247 | 32133 | <0.001*** |
| Retrieval | 1.781 | 32133 | 0.075 |
| Feedback | -2.134 | 32133 | 0.033* |
| Re-encoding | -1.787 | 32133 | 0.074 |
| Memory performance : trial index | 1.458 | 32133 | 0.145 |
| Memory performance : cue | -0.004 | 32133 | 0.996 |
| Memory performance : retrieval | 0.407 | 32133 | 0.684 |
| Memory performance : feedback | 1.329 | 32133 | 0.184 |
| Memory performance : re-encoding | 1.628 | 32133 | 0.103 |
| Trial index : cue | 0.093 | 32133 | 0.926 |
| Trial index : retrieval | -1.802 | 32133 | 0.071 |
| Trial index : feedback | 0.826 | 32133 | 0.409 |
| Trial index : re-encoding | -0.837 | 32133 | 0.403 |
| Memory performance : trial index : cue | -0.711 | 32133 | 0.477 |
| Memory performance : trial index : retrieval | -0.285 | 32133 | 0.776 |
| Memory performance : trial index : feedback | -1.029 | 32133 | 0.304 |
| Memory performance : trial index : re-encoding | -0.690 | 32133 | 0.490 |

\* $P < 0.05$ ; \*\*\* $P < 0.001$ . DF, degrees of freedom; :, interaction.
Model formula: IED prevalence $\sim 1 + \text{memory performance} * \text{trial index} + \text{memory performance} * \text{trial phase} + \text{trial index} * \text{trial phase} + \text{memory performance} : \text{trial index} : \text{trial phase} + (1 | \text{channel index})$ .

**Table S4.** Linear mixed model to analyze ripple rates (dependent variable) as a function of behavior.

| Predictor | *t* | DF | *P* |
| --- | --- | --- | --- |
| Memory performance | -0.106 | 32133 | 0.915 |
| Trial index | 0.343 | 32133 | 0.732 |
| Cue | 3.351 | 32133 | <0.001*** |
| Retrieval | -4.815 | 32133 | <0.001*** |
| Feedback | -9.416 | 32133 | <0.001*** |
| Re-encoding | -5.808 | 32133 | <0.001*** |
| Memory performance : trial index | 1.351 | 32133 | 0.177 |
| Memory performance : cue | 2.599 | 32133 | 0.009** |
| Memory performance : retrieval | 0.250 | 32133 | 0.803 |
| Memory performance : feedback | 0.583 | 32133 | 0.560 |
| Memory performance : re-encoding | -2.754 | 32133 | 0.006** |
| Trial index : cue | -3.483 | 32133 | <0.001*** |
| Trial index : retrieval | -0.801 | 32133 | 0.423 |
| Trial index : feedback | -2.824 | 32133 | 0.005** |
| Trial index : re-encoding | 1.259 | 32133 | 0.208 |
| Memory performance : trial index : cue | -1.327 | 32133 | 0.185 |
| Memory performance : trial index : retrieval | -1.672 | 32133 | 0.094 |
| Memory performance : trial index : feedback | -1.333 | 32133 | 0.183 |
| Memory performance : trial index : re-encoding | -0.173 | 32133 | 0.862 |

\*\**P* < 0.01; \*\*\**P* < 0.001. DF, degrees of freedom; :, interaction.
Model formula: ripple rate ~ 1 + memory performance \* trial index + memory performance \* trial phase + trial index \* trial phase + memory performance : trial index : trial phase + (1 | channel index).

**Table S5.** Linear mixed model to analyze ripple rates (dependent variable) as a function of behavior and artifacts.

| **Predictor** | ***t*** | **DF** | ***P*** |
| --- | --- | --- | --- |
| Artifact prevalence | -22.317 | 32132 | <0.001*** |
| Memory performance | -0.119 | 32132 | 0.905 |
| Trial index | 0.628 | 32132 | 0.530 |
| Cue | 2.685 | 32132 | 0.007** |
| Retrieval | -4.696 | 32132 | <0.001*** |
| Feedback | -9.772 | 32132 | <0.001*** |
| Re-encoding | -6.166 | 32132 | <0.001*** |
| Memory performance : trial index | 1.564 | 32132 | 0.118 |
| Memory performance : cue | 2.613 | 32132 | 0.009** |
| Memory performance : retrieval | 0.308 | 32132 | 0.758 |
| Memory performance : feedback | 0.775 | 32132 | 0.438 |
| Memory performance : re-encoding | -2.568 | 32132 | 0.010* |
| Trial index : cue | -3.495 | 32132 | <0.001*** |
| Trial index : retrieval | -1.021 | 32132 | 0.307 |
| Trial index : feedback | -2.747 | 32132 | 0.006** |
| Trial index : re-encoding | 1.149 | 32132 | 0.251 |
| Memory performance : trial index : cue | -1.438 | 32132 | 0.150 |
| Memory performance : trial index : retrieval | -1.722 | 32132 | 0.085 |
| Memory performance : trial index : feedback | -1.464 | 32132 | 0.143 |
| Memory performance : trial index : re-encoding | -0.268 | 32132 | 0.788 |

\* $P < 0.05$ ; \*\* $P < 0.01$ ; \*\*\* $P < 0.001$ . DF, degrees of freedom; :, interaction.
Model formula: ripple rate $\sim 1 + \text{artifact prevalence} + \text{memory performance} * \text{trial index} + \text{memory performance} * \text{trial phase} + \text{trial index} * \text{trial phase} + \text{memory performance} : \text{trial index} : \text{trial phase} + (1 | \text{channel index})$ .

## Supplementary Figures

**Fig. S1.**
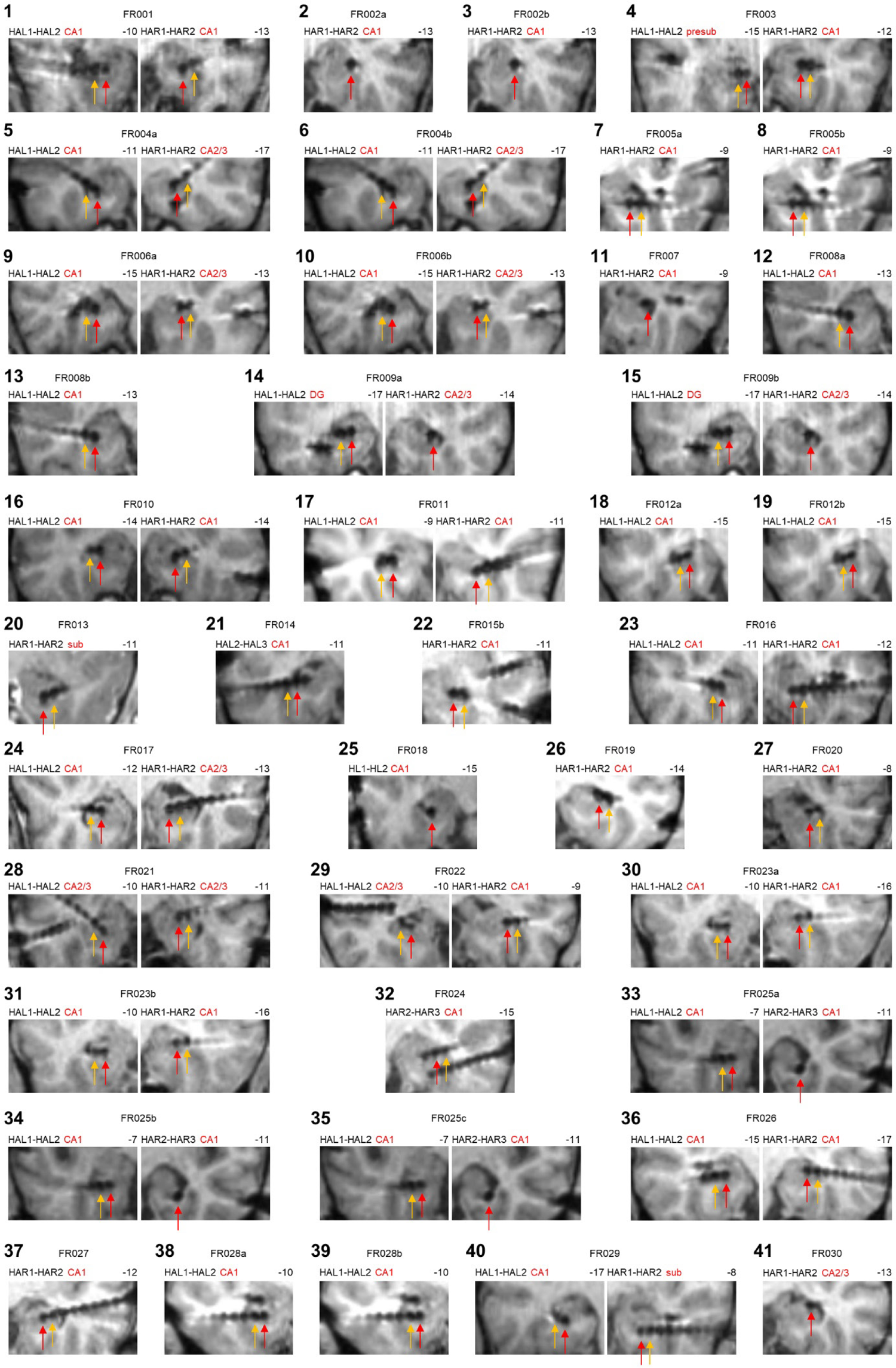
Depiction of the hippocampal macroelectrode channels from all 41 sessions. The locations of the hippocampal electrodes are presented on coronal slices of the post-operative MRI scans. Red arrow, first electrode channel contributing to the bipolar channel; yellow arrow, second electrode channel contributing to the bipolar channel. Black bold large numbers indicate the session numbers. Numbers right and above the images indicate the y-coordinate of the coronal slice in MNI space. Red labels, hippocampal subregions based on visual inspection following (*36*). CA1, cornu ammonis region 1; CA2/3, cornu ammonis region 2 or 3; DG, dentate gyrus; presub, presubiculum; sub, subiculum. HAL, hippocampus anterior left; HAR, hippocampus anterior right; HL, hippocampus left.

**Fig. S2.**
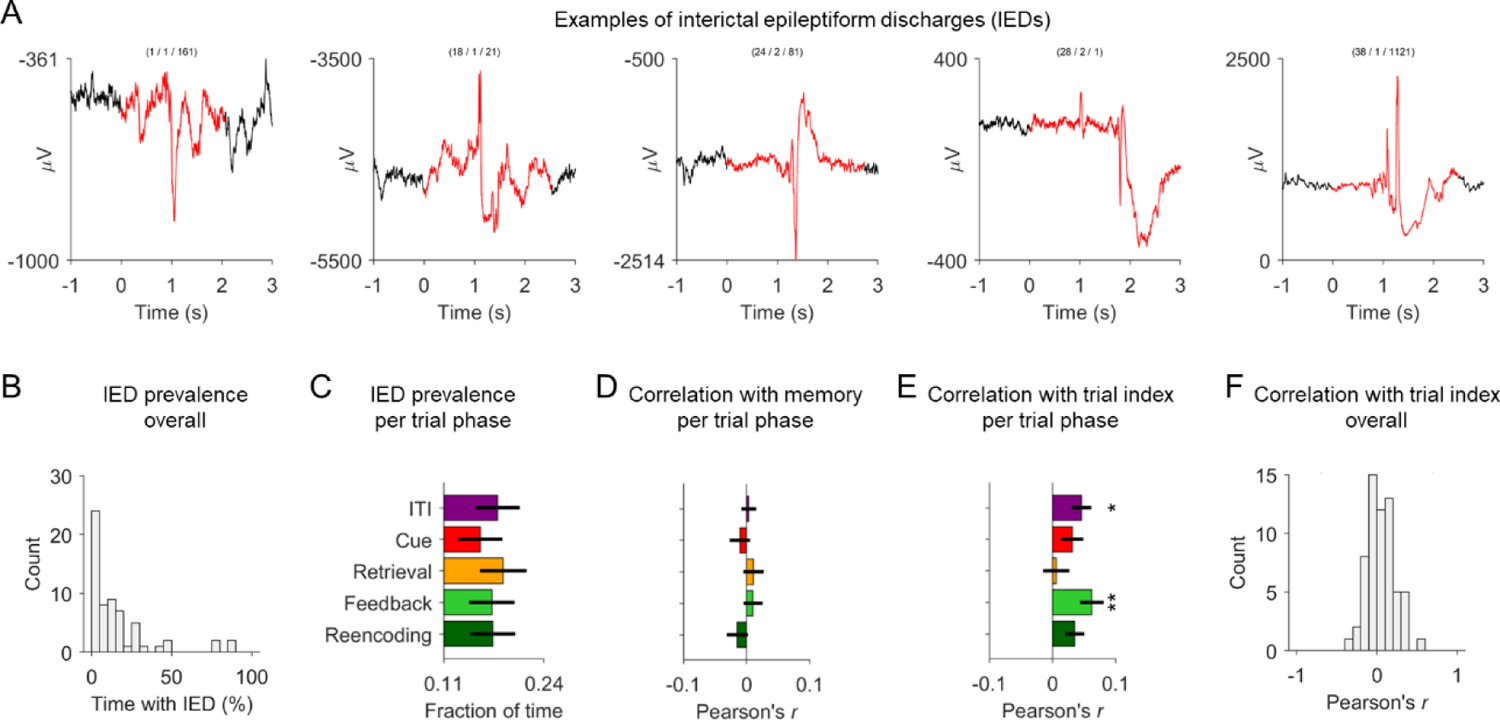
Interictal epileptiform discharges (IEDs). (**A**) Examples of IEDs from five different sessions, which were identified using an automated procedure. Around each IED, an additional time period of ±1 s was excluded (*34*). Black, local field potential (not filtered); red, local field potential with an IED and thus excluded from subsequent ripple detection. (**B**) Histogram showing the prevalence of IEDs in each session. On average, 16.591 ± 2.692% (mean ± SEM; *n* = 62 channels) of the data was designated as belonging to an IED. (**C**) Prevalence of IEDs in different trial phases. A repeated measures ANOVA identified a significant modulation of IED prevalence by trial phase [*F*(4, 244) = 3.705, *P* = 0.006]. Post-hoc comparisons showed that the prevalence of IEDs was significantly reduced during the cue period as compared to the ITI period (*P*Tukey-Kramer = 0.012) and the retrieval period (*P*Tukey-Kramer < 0.001). (**D**) Correlation between the prevalence of IEDs and memory performance, separately for each trial phase. Correlations were computed separately for each channel and then averaged across channels. Correlation coefficients were not significantly different from 0 (one-sample *t*-tests across channel-wise correlation coefficients: all *t* < 0.899, all *P* = 1, Bonferroni corrected for five tests). (**E**) Correlation between the prevalence of IEDs and trial index, separately for each trial phase. Correlations were computed separately for each channel and then averaged across channels. Correlation coefficients were significantly above 0 for ITI [one-sample *t*-test across channel-wise correlation coefficients: *t*(60) = 2.985, *P* = 0.021, Bonferroni corrected for five tests] and feedback [*t*(57) = 3.339, *P* = 0.007, Bonferroni corrected for five tests]. (**F**) Correlation between the prevalence of IEDs and trial index, irrespective of trial phase. Correlation coefficients were significantly above 0, indicating that the prevalence of IEDs increased over the course of a session [one-sample *t*-test: *t*(61) = 2.207, *P* = 0.031]. \**P* < 0.05; \*\**P* < 0.01.

**Fig. S3.**
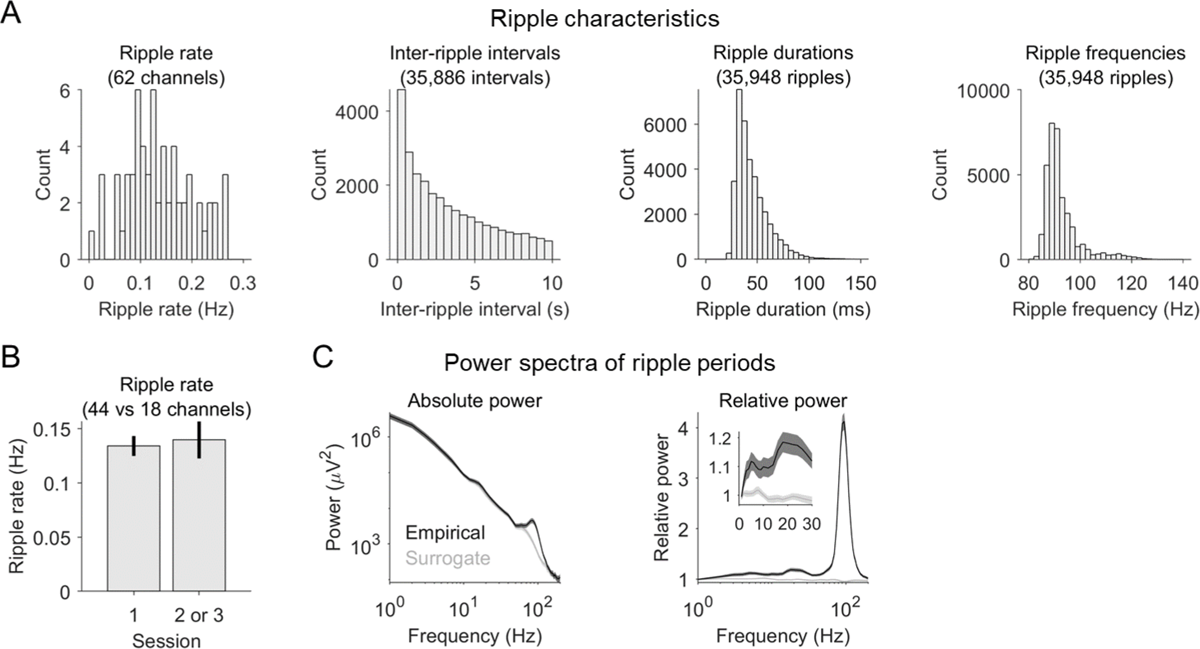
Characteristics of hippocampal ripples. (**A**) Different ripple characteristics including ripple rates, inter-ripple intervals, ripple durations, and ripple frequencies. The identified ripples occurred at a rate of 0.136 ± 0.008 ripples/s (mean ± SEM; *n* = 62 channels; 0.159 ± 0.007 ripples/s when only considering artifact-free time periods) and exhibited an average inter-ripple interval of 7.382 ± 0.084 s (mean ± SEM; *n* = 35886 inter-ripple intervals). The ripples had an average duration of 45.210 ± 0.086 ms (mean ± SEM; *n* = 35948 ripples) and an average frequency of 92.603 ± 0.036 Hz (mean ± SEM; *n* = 35948 ripples). These ripple characteristics are comparable to previous studies using similar ripple-detection algorithms [e.g., (*34*, *39*, *45*)]. (**B**) Ripple rates during first sessions were not different from ripple rates during second or third sessions [two-sample *t*-test: *t*(60) = −0.307, *P* = 0.760, *n* = 44 channels in first sessions, *n* = 18 channels in second or third sessions]. (**C**) Raw and relative power spectrum of hippocampal ripples and hippocampal surrogate ripples. Relative power was estimated by dividing the power values during the ripples by the mean power values averaged across the entire experiment (separately for each frequency). As expected, raw and relative power spectra of ripple periods showed marked peaks at about 93 Hz.

**Fig. S4.**
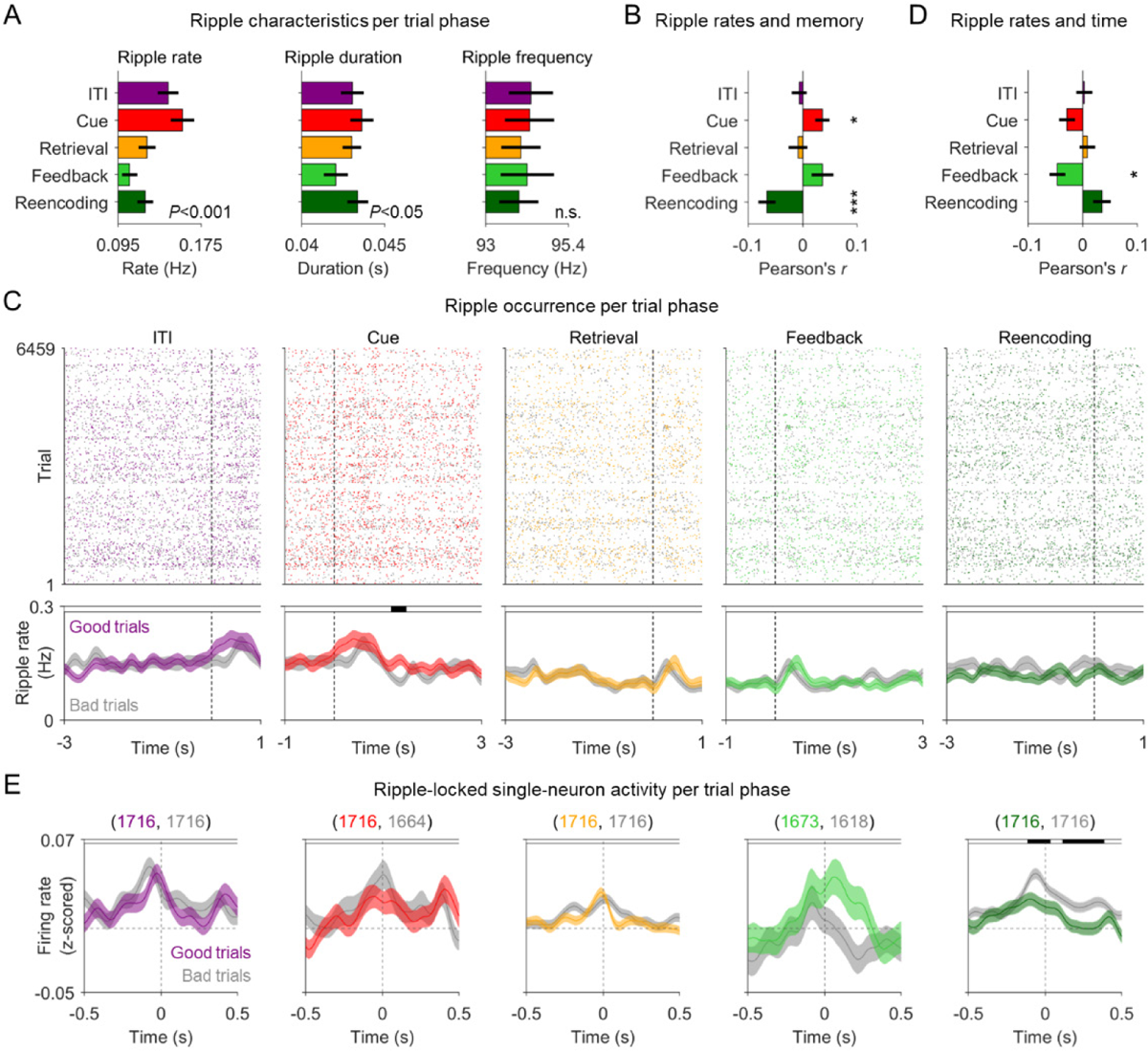
Hippocampal ripple rates are linked to behavioral state and memory performance in the associative memory task. (**A**) Ripple characteristics (rate, duration, and frequency) during the different trial phases. As compared to inter-trial interval (ITI) periods, ripple rates were significantly increased during cue periods and significantly reduced during retrieval, feedback, and re-encoding periods. Ripple durations showed a similar pattern as ripple rates (but no significant post-hoc comparisons). Ripple frequency was not modulated by trial phase. (**B**) Correlation between ripple rates and memory performance, separately for the different trial phases. (**C**) Correlation between ripple rates and trial index, separately for the different trial phases. (**D**) Time-resolved ripple occurrence across all trials as a function of absolute time relative to the onset (cue and feedback) or offset (ITI, retrieval, and re-encoding) of a given trial phase (dashed vertical lines), separately for the different trial phases. Each dot is a ripple (colored dots, ripples during good-performance trials; gray dots, ripples during bad-performance trials). Colored lines and shadings, ripple rates during good-performance trials (“good trials”; mean ± SEM across channels); gray lines and shadings, ripple rates during bad-performance trials (“bad trials”; mean ± SEM across channels). Black shadings at top indicate time periods with significant differences between good and bad trials (two-sided cluster-based permutation tests across the entire depicted time window: *P* < 0.025). For ITI, retrieval, and re-encoding, the plots focus on the last three seconds before the end of these periods (at time 0), although their durations differed between trials. (**E**) Neuronal firing rates during hippocampal ripples occurring in specific trial phases. Firing rates are smoothed with a Gaussian kernel of 0.2-s duration and *z*-scored relative to the entire experiment. Colored lines and shadings (mean ± SEM) indicate firing rates during ripples from trials with good memory performance; gray lines and shadings (mean ± SEM) indicate firing rates during ripples from trials with bad memory performance. Time 0, hippocampal ripple peak. Black shadings at the top of the subpanels indicate significantly different firing rates during ripples from good-performance trials versus firing rates during ripples from bad-performance trials (two-sided cluster-based permutation tests across the entire depicted time window: *P* < 0.025). Colored numbers indicate the number of neuron–ripple channel combinations contributing to the data. ITI, inter-trial interval. \**P* < 0.05; \*\*\**P* < 0.001.

**Fig. S5.**
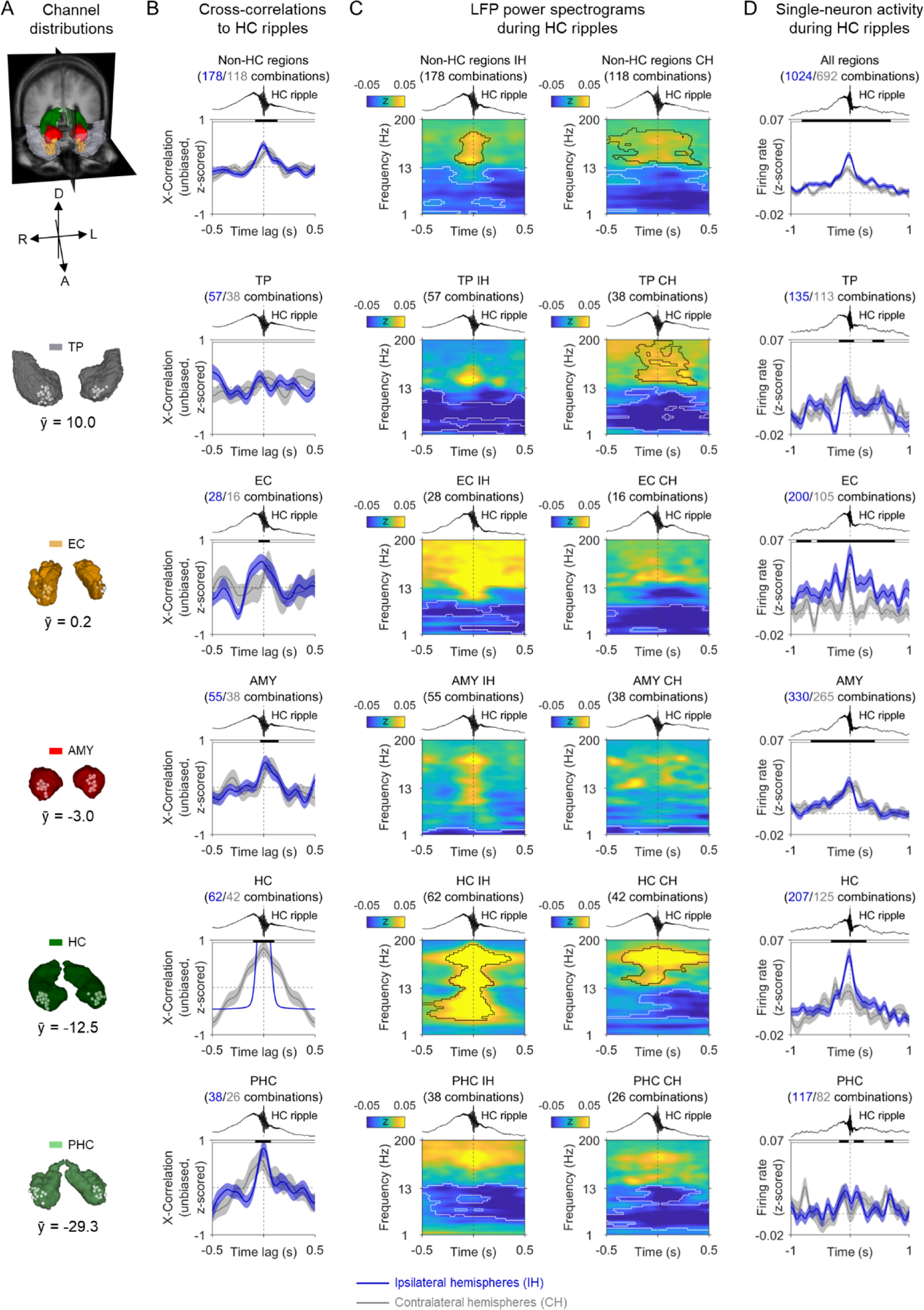
Hippocampal ripples are associated with changes in LFP power and firing rates across the human MTL. (**A**) Probabilistic visualizations of regions of interest including temporal pole (TP), entorhinal cortex (EC), amygdala (AMY), hippocampus (HC), and parahippocampal cortex (PHC), listed according to their anterior–to–posterior position in the human brain. White dots, locations of bipolar electrode contacts pooled across subjects; ȳ, average MNI y-value of the contacts in a given region of interest. (**B**) Cross-correlations between hippocampal ripples and ripples in the different MTL regions. Blue and gray numbers indicate the number of channel pairs from ipsilateral and contralateral hemispheres, respectively. Time 0 indicates the peak of the hippocampal ripples; cross-correlation maxima at positive time lags indicate that the ripples from a particular region of interest numerically precede hippocampal ripples. Cross-correlations are smoothed with a Gaussian kernel of 0.2-s duration and are normalized by *z*-scoring cross-correlation values over the displayed time lags of ±0.5 s. Shaded region, SEM across channels pairs. Black shadings at top indicate *z*-scored cross-correlations from both ipsilateral and contralateral channel pairs being significantly above 0 (one-sided cluster-based permutation tests across the entire depicted time window: *P* < 0.05). The blue line in the subpanel for hippocampal channels is the smoothed temporal autocorrelation and is only shown for the sake of completeness. (**C**) Time-frequency resolved LFP power (*z*-scored relative to the entire experiment) in the different MTL regions during hippocampal ripples, both for ipsilateral channel pairs (left column) and contralateral channel pairs (right column). Power values are smoothed over time with a Gaussian kernel of 0.2-s duration. Time 0 indicates the peak of the hippocampal ripples. Black contours, significantly increased power; white contours, significantly decreased power (two-sided cluster-based permutation tests across the entire depicted time window: *P* < 0.025). (**D**) Neuronal firing rates (*z*-scored relative to the entire experiment) in hippocampal and extrahippocampal regions (recorded using microelectrodes) during hippocampal ripples (recorded using macroelectrodes). Firing rates are smoothed over time with a Gaussian kernel of 0.2-s duration. Blue and gray numbers indicate the counts of ipsilateral and contralateral neuron–ripple channel pairs, respectively. Black shadings at top indicate firing rates from both ipsilateral and contralateral neuron–ripple channel combinations significantly above 0 (one-sided cluster-based permutation tests across the entire depicted time window: *P* < 0.05). IH, ipsilateral hemispheres; CH, contralateral hemispheres. AMY, amygdala; EC, entorhinal cortex; HC, hippocampus; PHC, parahippocampal cortex; TP, temporal pole. A, anterior; D, dorsal; L, left; R, right. X-Correlation, cross-correlation.

**Fig. S6.**
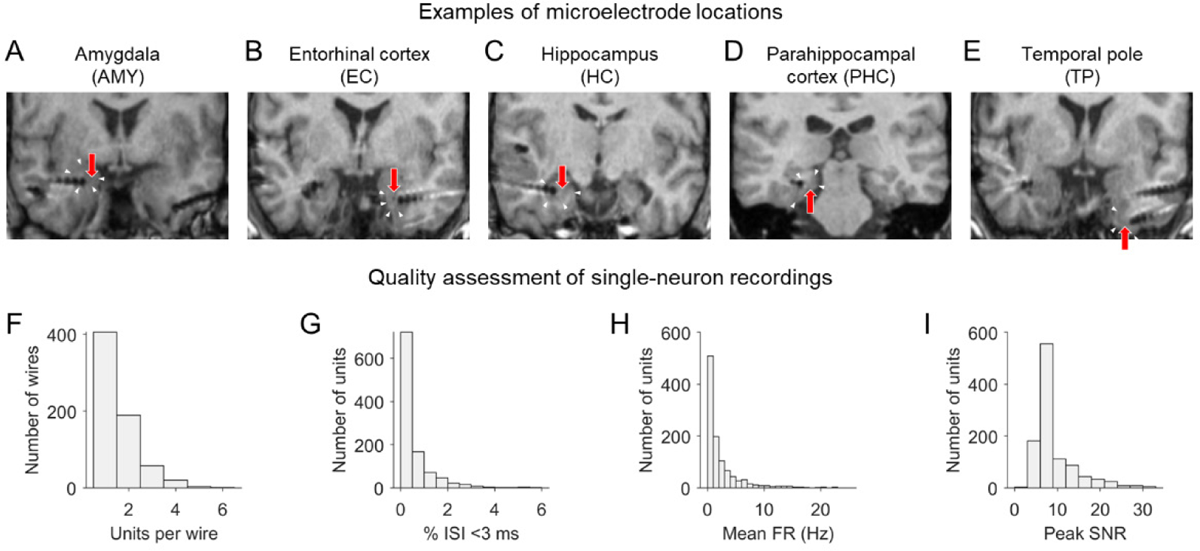
Single-neuron recordings: anatomical locations and quality assessment. (**A–E**) Example microelectrodes in amygdala (AMY), entorhinal cortex (EC), hippocampus (HC), parahippocampal cortex (PHC), and temporal pole (TP). Electrode contacts of macroelectrodes appear as dark circles on the MRI scans. Red arrows point at putative microelectrode locations, which protrude 3–5 mm from the tip of the depth electrode (often not visible on MRI scans). White triangles indicate the borders of the different brain regions. (**F**) Histogram of units per wire. On average, 1.570 ± 0.032 (mean ± SEM) units per wire were recorded (only considering wires with at least one unit). (**G**) Histogram of the percentages of inter-spike intervals (ISIs) that were shorter than 3 ms. On average, units exhibited 0.507 ± 0.024% (mean ± SEM) ISIs that were shorter than 3 ms. (**H**) Histogram of mean firing rates (FRs). On average, units exhibited mean FRs of 2.402 ± 0.108 (mean ± SEM) Hz, which is comparable to previous human single-neuron studies [e.g., (*38*)]. (**I**) Histogram of the mean waveform peak signal-to-noise ratio (SNR) of each unit. On average, the SNR of the mean waveform peak was 9.181 ± 0.160 (mean ± SEM).

**Fig. S7.**
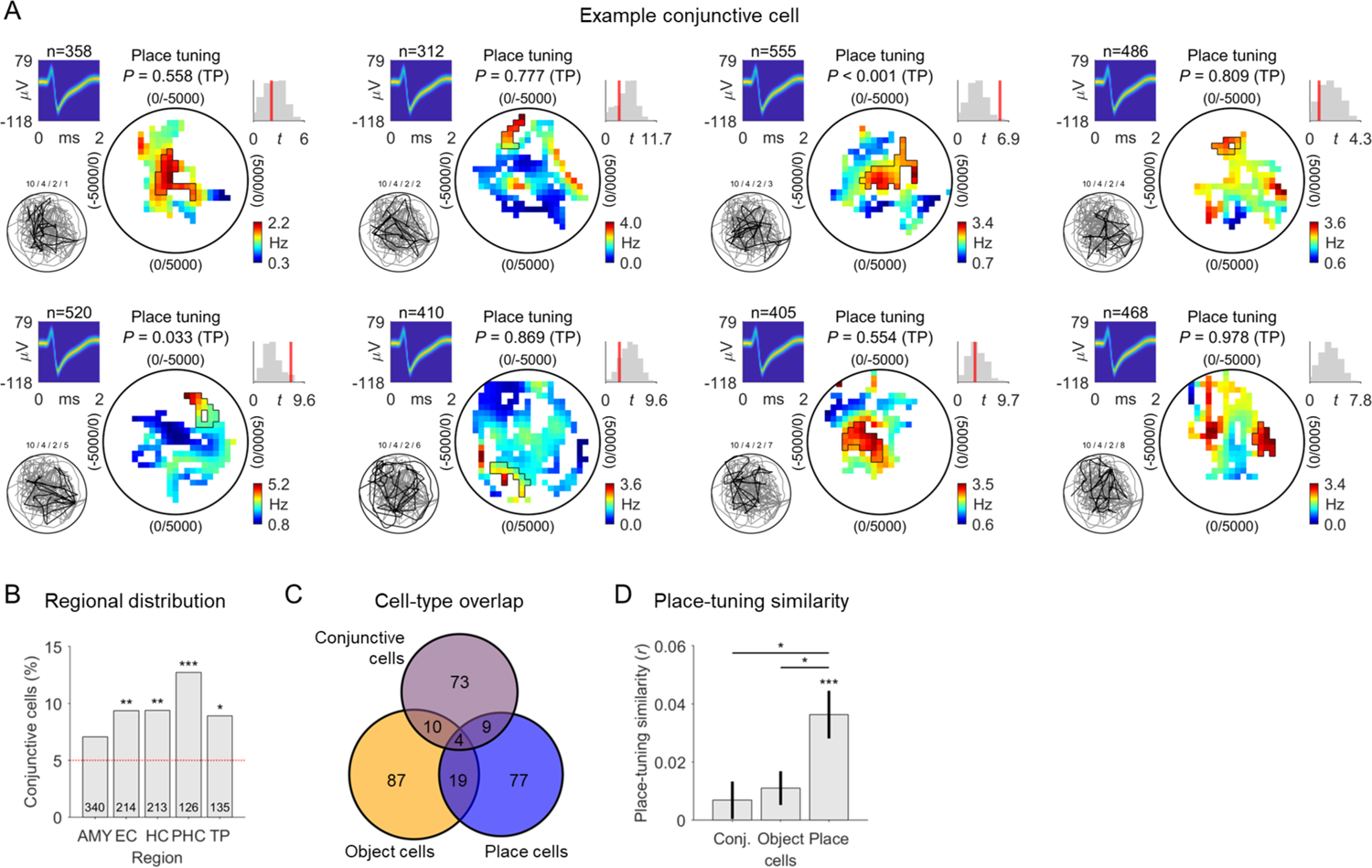
Conjunctive cells. (**A**) Example conjunctive cell. The eight different panels show the cell’s place tuning when only considering data from trials with a particular object. This example conjunctive cell exhibited significant place tuning at a Bonferroni-corrected alpha-level of α = 0.05/8 only during trials with object #3 (first row, third column). For each panel, from left to right: action potentials as density plot; entire (gray line) and object-specific (black line) navigation path of the subject through the environment; smoothed firing-rate map (unvisited areas are shown in white); empirical *t*-statistic (red line) and surrogate *t*-statistics (gray histogram); color bar, firing rate. Candidate place fields are outlined in black. (**B**) Distribution of conjunctive cells across brain regions; red line, 5% chance level. (**C**) Overlap between object cells, place cells, and conjunctive cells. The large majority of object and place cells were not also conjunctive cells. (**D**) Similarity of place tuning across trials with the eight different objects, which we estimated using pairwise Pearson correlations between the firing-rate maps from trials with different objects. Place tuning was significantly similar (and, thus, stable) across trials with different objects for place cells [one-sample *t*-test, *t*(108) = 4.417, *P* < 0.001], but not for conjunctive cells [*t*(95) = 1.056, *P* = 0.294] and also not for object cells [*t*(119) = 1.888, *P* = 0.061]. Place-tuning similarity was significantly higher in place cells than in conjunctive cells [two-sample *t*-test: *t*(203) = 2.772, *P* = 0.006] and also significantly higher in place cells than in object cells [*t*(227) = 2.550, *P* = 0.011]. These results underline the general spatial coding of place cells (being stable across time irrespective of the current object) and confirms that conjunctive cells (as well as object cells) do not exhibit such general spatial coding, as expected. Conj., conjunctive. AMY, amygdala; EC, entorhinal cortex; HC, hippocampus; PHC, parahippocampal cortex; TP, temporal pole. \**P* < 0.05; \*\**P* < 0.01; \*\*\**P* < 0.001.

**Fig. S8.**
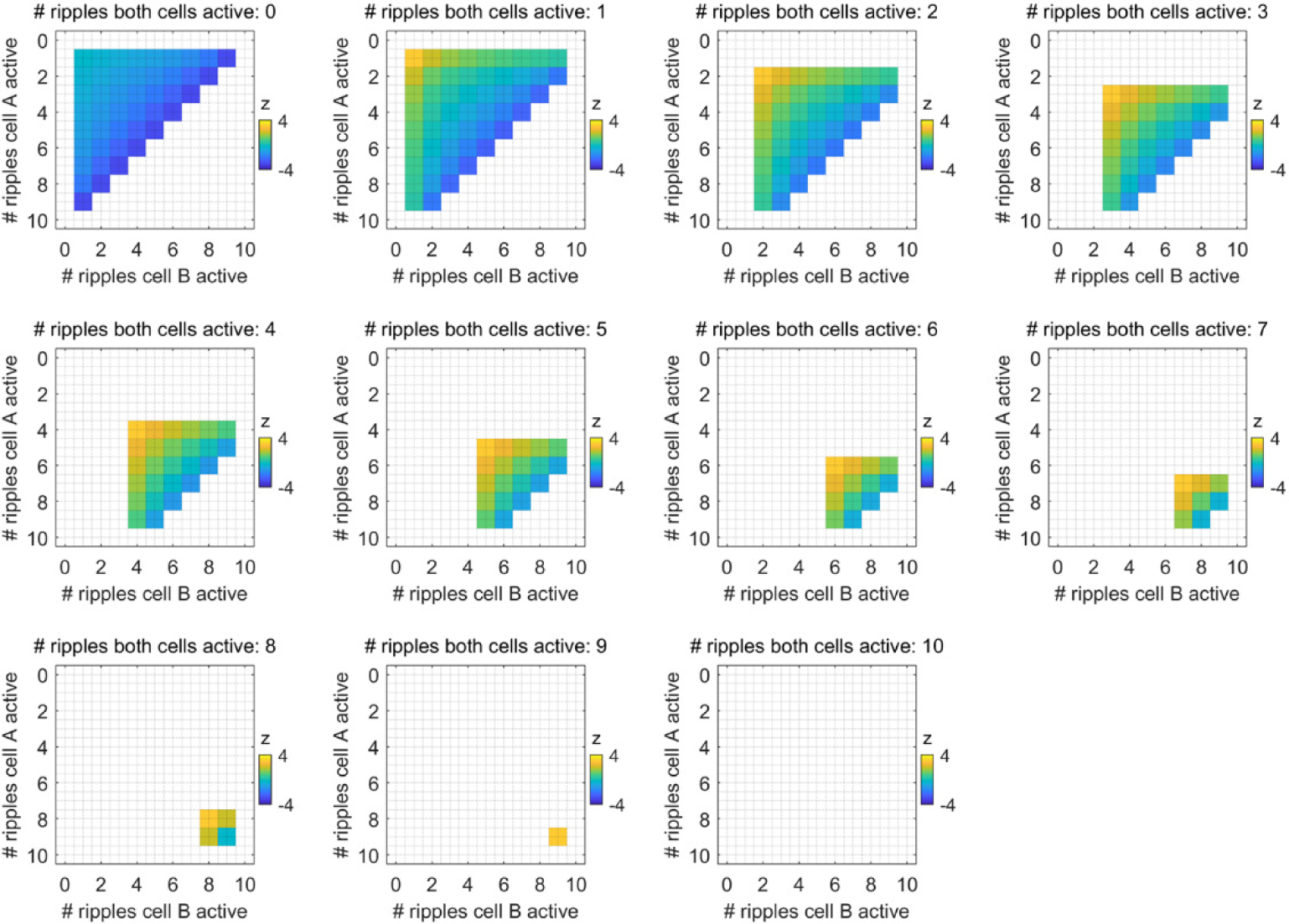
Illustration of the coactivity *z*-score. Coactivity *z*-scores between two cells (cell A and cell B) in a hypothetical situation with ten ripples. The different subplots display the coactivity *z*-scores as a function of the number of ripples in which cell A is active (y-axis), the number of ripples in which cell B is active (x-axis), and the number of ripples in which both cells are active (title). High coactivity *z*-scores are achieved when the number of ripples in which both cells are active matches the number of ripples in which cell A (irrespective of cell B) or cell B (irrespective of cell A) is active. The coactivity *z*-value is not defined (white areas) for situations (i) in which cell A and/or cell B are not active during any ripple; (ii) in which cell A and/or cell B are active during all ripples; and (iii) in which the activity combination is logically impossible. #, number.

**Fig. S9.**
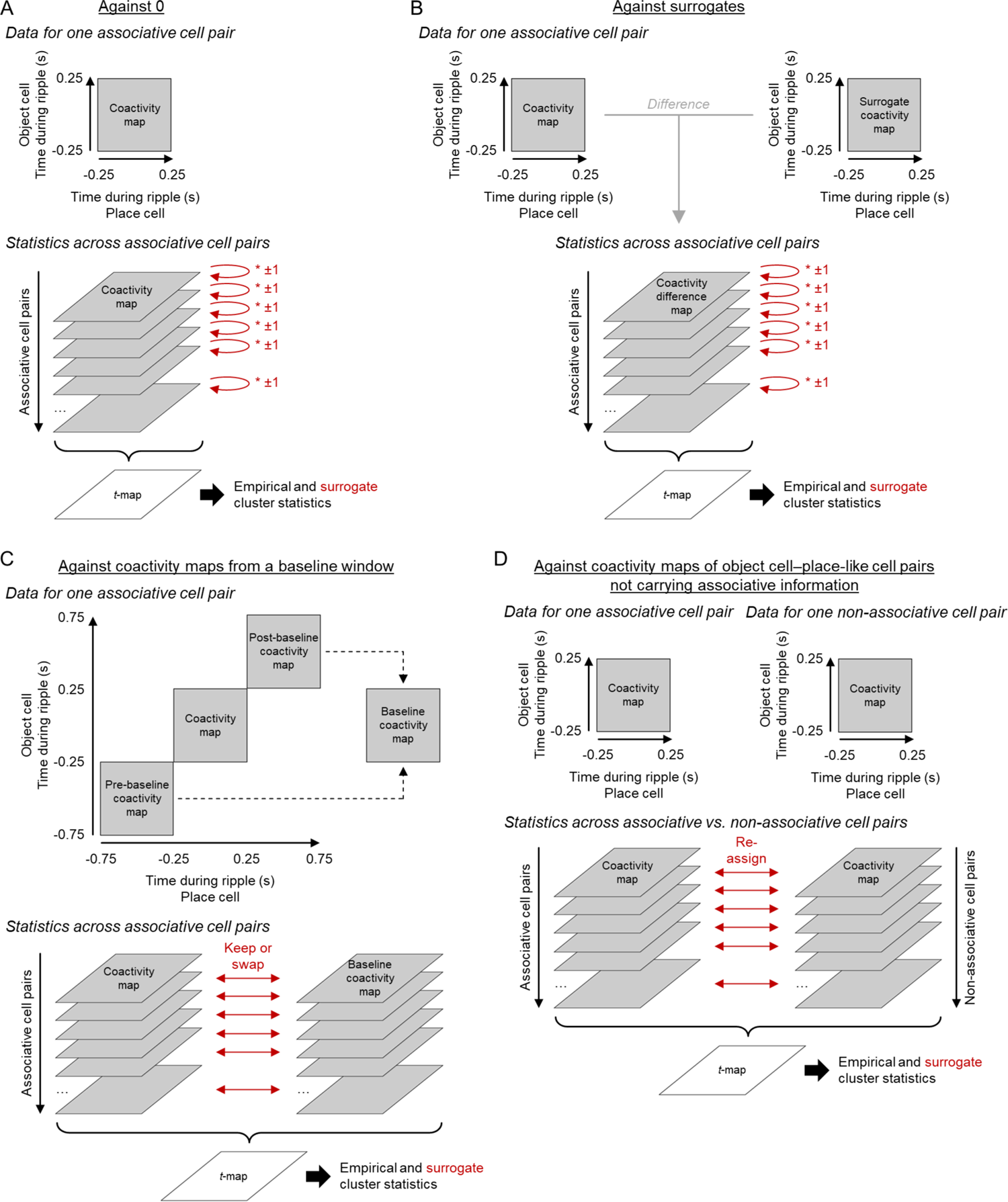
Illustration of the cluster-based permutation tests. (**A**) Cluster-based permutation test of the coactivity maps against chance (i.e., 0). To identify empirical clusters, a one-sample *t*-test against 0 was performed for each time–by–time bin. Contiguous clusters of significant bins were identified, and the corresponding *t*-values were summed, resulting in empirical cluster statistics. To create surrogate data, a randomly selected subset of the coactivity maps was multiplied by −1. Performing the same steps as for the empirical data, surrogate cluster statistics were then created based on the surrogate data. (**B**) Variant of the cluster-based permutation test against chance, where a surrogate coactivity map was subtracted from each corresponding empirical coactivity map. The surrogate coactivity of a given cell pair was estimated by circularly shifting the ripple-locked activity levels of the object cell relative to the ripple-locked activity levels of the place cell by a random latency before calculating the two-dimensional coactivity map. The statistics across associative cell pairs were performed as in A. (**C**) Cluster-based permutation test of the coactivity maps against coactivity maps from a baseline window. The pre-baseline and post-baseline windows spanned −0.75 to −0.25 s before and 0.25 to 0.75 s after the ripple peaks, respectively. Baseline coactivity maps were then estimated by averaging the pre-baseline and the post-baseline coactivity maps. To identify empirical cluster statistics, a two-sample *t*-test between the coactivity maps and the baseline coactivity maps was performed for each time-by-time bin. Contiguous clusters of significant bins were identified, and the corresponding *t*-values were summed up, resulting in empirical cluster statistics. To create surrogate data, a random subset of corresponding coactivity maps and baseline coactivity maps was swapped. Performing the same steps as for the empirical data, surrogate cluster statistics were then created based on the surrogate data. (**D**) Cluster-based permutation test of the coactivity maps of associative cell pairs versus non-associative cell pairs (for which the location of the preferred object of the object cell was not located inside the place field of the place cell). To identify empirical clusters, a two-sample *t*-test between the coactivity maps of associative cell pairs and those of non-associative cell pairs was performed for each time-by-time bin. Contiguous clusters of significant time bins were identified, and the corresponding *t*-values were summed up, resulting in empirical cluster statistics. To create surrogate data, the coactivity maps of associative and non-associative cell pairs were randomly reassigned to the two cell groups. Performing the same steps as for the empirical data, surrogate cluster statistics were then created based on the surrogate data. In all different tests, empirical clusters were considered significant if their cluster statistics exceeded the 95^th^ percentile of all surrogate cluster statistics. Red arrows indicate how surrogate data were created.

**Fig. S10.**
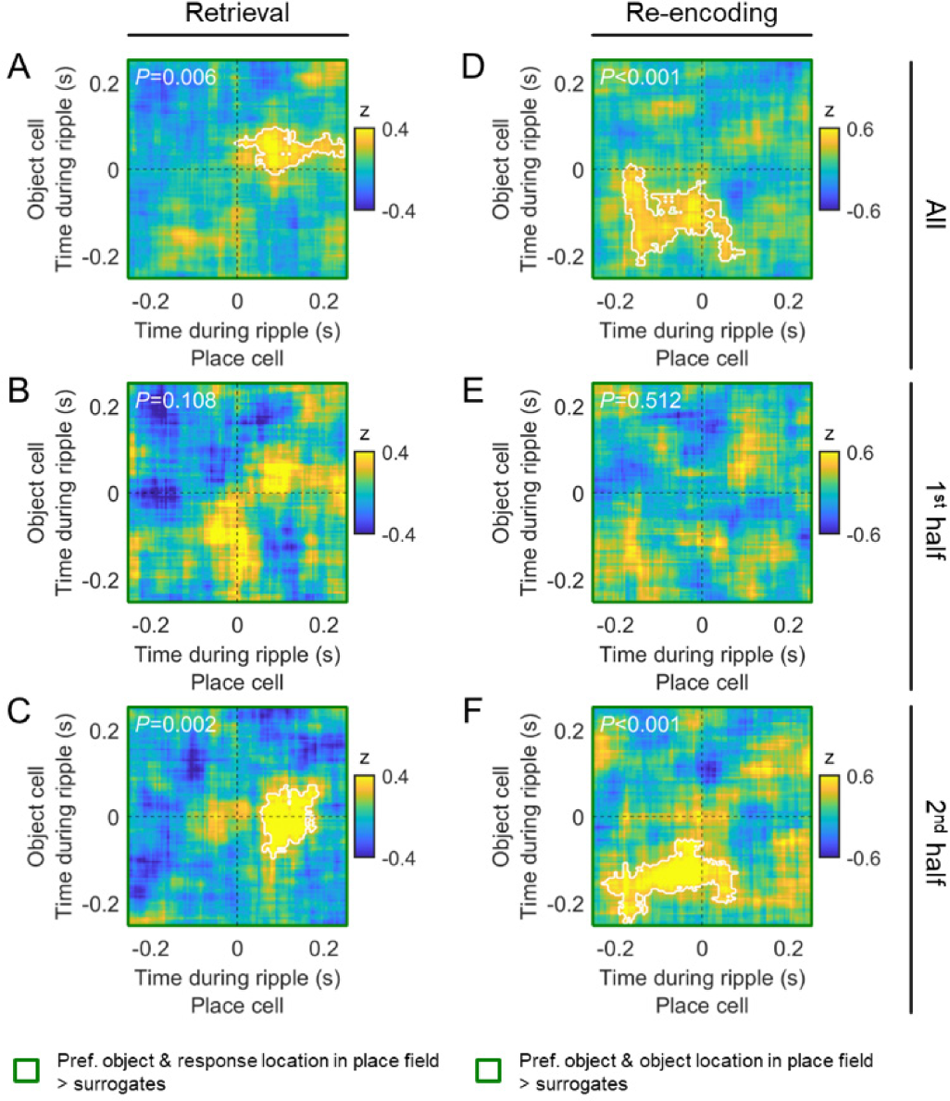
Coactivity of object cells and place cells during hippocampal ripples: comparison against surrogate coactivity maps. (**A–F**) Same as the left panels in Fig. 4, C–H, with the difference that the coactivity maps are contrasted against surrogate coactivity maps instead of against 0. For each cell pair, one surrogate coactivity map was estimated by circularly shifting the spiking activity of the object cell relative to the spiking activity of the place cell by a random latency across ripples. White lines delineate significant clusters based on cluster-based permutation tests, whose *P*-values are stated in the upper left corners of the coactivity maps. >, larger than; pref., preferred.

**Fig. S11.**
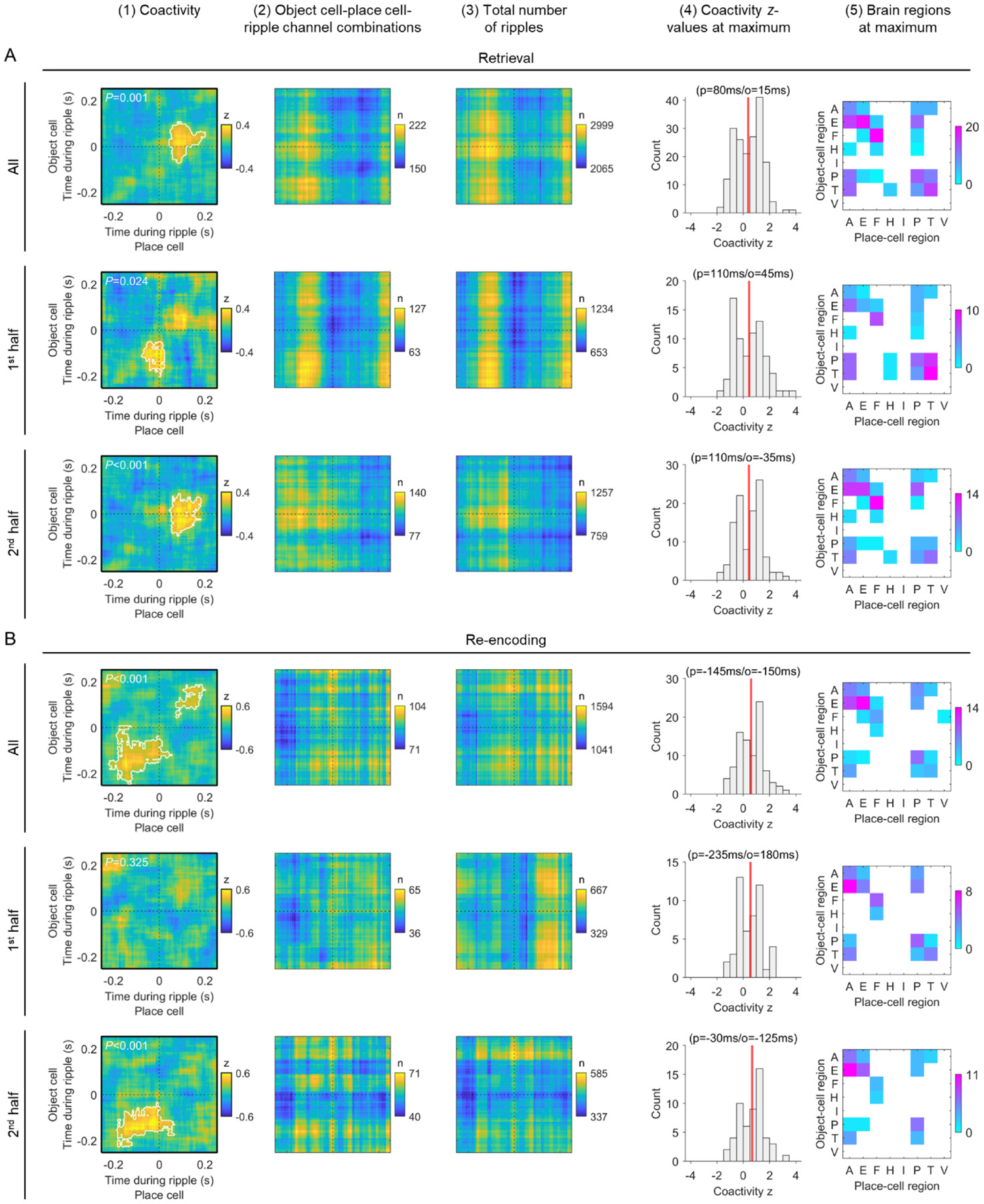
Coactivity of object cells and place cells during hippocampal ripples: additional information. Columns from left to right: (1) coactivity maps reproduced from Fig. 4, C–H, left column. (2) Number of object cell–place cell–ripple channel combinations underlying the coactivity maps in the first column. (3) Number of ripples underlying the coactivity maps in the first column, pooled across object cell–place cell–ripple channel combinations. (4) Histograms of individual coactivity *z*-values underlying the global maxima in the coactivity maps in the first column. Red line, mean; numbers above the histograms indicate the temporal coordinates of the global maxima in the coactivity maps (p, temporal coordinate for the place-cell time axis; o, temporal coordinate for the object-cell time axis). (5) Brain regions of the object cells and place cells contributing the coactivity *z*-values of the global peaks of the coactivity maps in the first column; color bars, number of object cell–place cell combinations. (**A**) Data for retrieval. (**B**) Data for re-encoding. A, amygdala; E, entorhinal cortex; F, fusiform gyrus; H, hippocampus; I, insula; P, parahippocampal cortex; T, temporal pole; V, visual cortex.

**Fig. S12.**
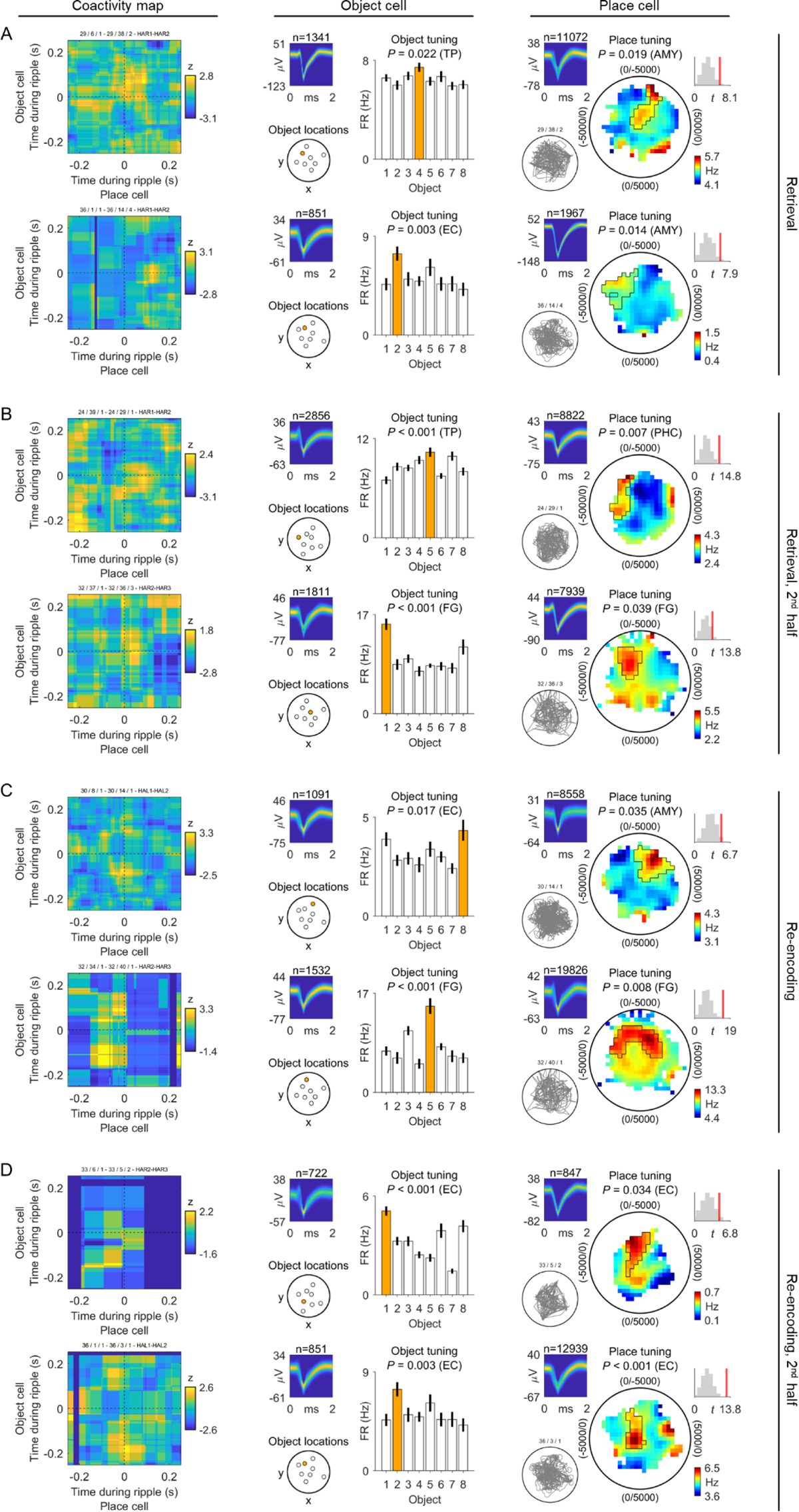
Coactivity of object cells and place cells during hippocampal ripples: individual examples. (**A**) Both rows show an individual example of the coactivity for an object cell–place cell–ripple channel combination, from ripples during the retrieval periods of the associative object–location memory task. Left column, coactivity map between a given object cell and a given place cell during the ripples of a given ripple channel. Middle column, information about the object cell including a density plot of the spike waveforms (upper left subpanel; number above subpanel indicates spike count), locations of the objects including the location of the preferred object (orange dot in the lower left subpanel), and average firing rates in response to the different objects during the cue periods (right subpanel). Right column, information about the place cell including a density plot of the spike waveforms (upper left subpanel), the subject’s navigation path through the virtual environment (lower left subpanel), the cell’s firing-rate map (middle subpanel), and the comparison between the cell’s empirical test statistic and its surrogate test statistics (upper right subpanel). Color bar, firing rate. (**B**) Same as in A, but for ripples occurring during the second half of all retrieval-related ripples. (**C**) Same as in A, but for ripples occurring during the re-encoding periods. (**D**) Same as in A, but for ripples occurring during the second half of all re-encoding-related ripples. AMY, amygdala; EC, entorhinal cortex; FG, fusiform gyrus; PHC, parahippocampal cortex; TP, temporal pole.

**Fig. S13.**
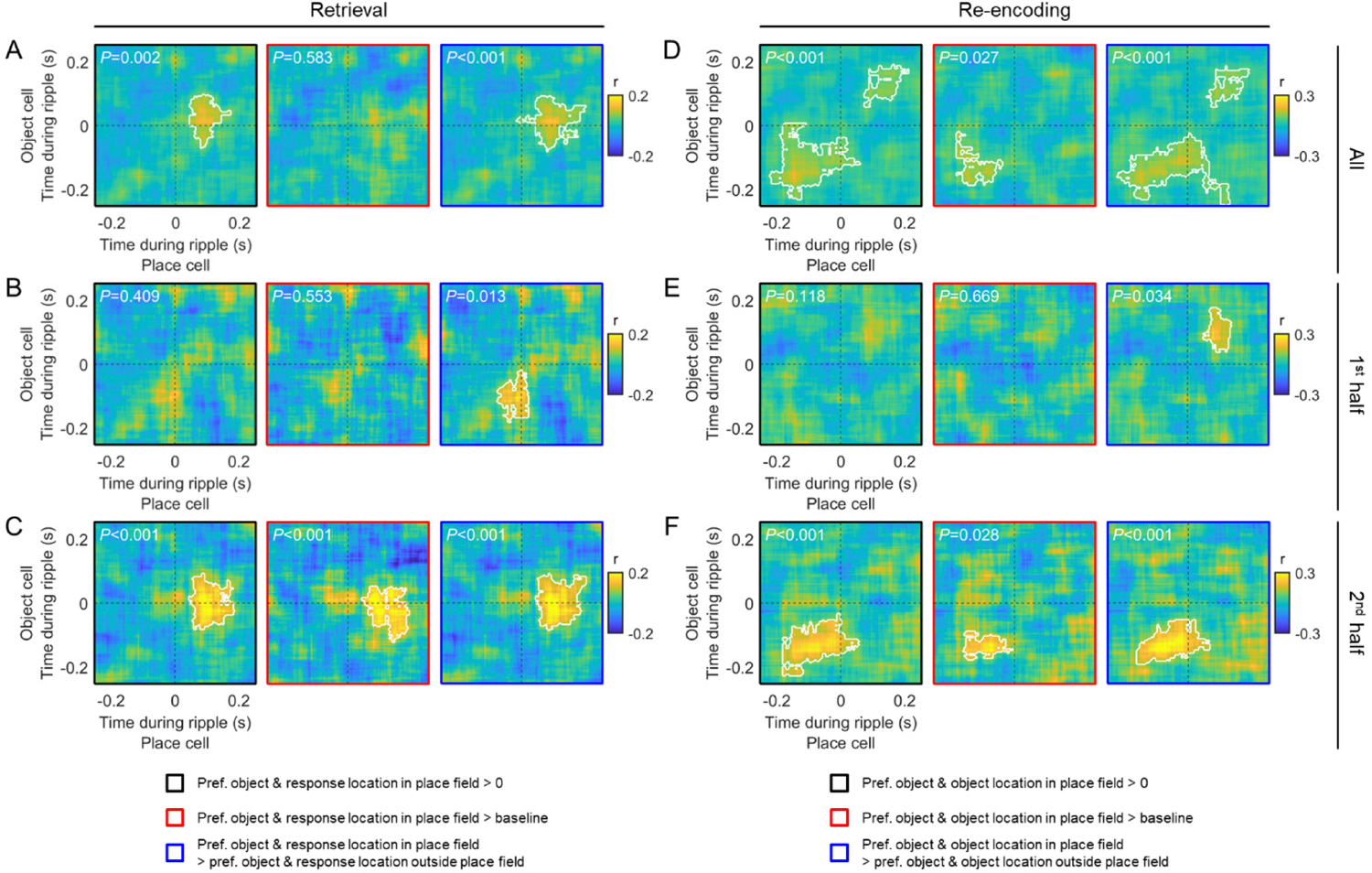
Coactivity of object cells and place cells during hippocampal ripples: Pearson correlations. (**A–F**) Same as Fig. 4, C–H, with the difference that coactivity was estimated with Pearson correlations. White lines delineate significant clusters based on cluster-based permutation tests, with *P*-values stated at the upper left. >, larger than; pref., preferred.

**Fig. S14.**
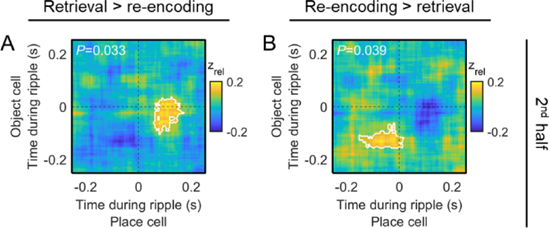
Coactivity of object cells and place cells during hippocampal ripples: temporal shift between retrieval and re-encoding. (**A**) Direct comparison of the coactivity maps from retrieval versus the coactivity maps from re-encoding. This demonstrates that the coactivations are higher for retrieval than for re-encoding at time points during and slightly after the ripple peaks (with regard to object cells and place cells, respectively). The retrieval-related coactivity maps are estimated using the 2^nd^ half of all ripples during retrieval, considering only trials in which the subject is asked to remember the location of the preferred object of the object cell and in which the subject’s response location is inside the place field of the place cell (Fig. 4E, left panel). The re-encoding-related coactivity maps are estimated using the 2^nd^ half of all ripples during re-encoding, considering only trials in which the subject is asked to re-encode the correct location of the preferred object of the object cell and in which the object’s correct location is inside the place field of the place cell (Fig. 4H, left panel). For this analysis, the coactivity *z*-scores of each cell pair were normalized (zrel) so that a particular coactivity *z*-score was: *z*rel-i = (*z*i − min(*z*)) / max(*z*), where *i* is the index of a particular coactivity *z*-score in a given coactivity map. For each cell pair, the normalized coactivity *z*-scores thus ranged between 0 and 1. This was done in order to enable the direct comparisons between retrieval- and re-encoding-related coactivity maps, as re-encoding-related coactivations were generally higher than retrieval-related coactivations (Fig. 4). (**B**) Direct comparison of the coactivity maps from re-encoding versus the coactivity maps from retrieval. This demonstrates that the coactivations were higher for re-encoding than for retrieval at time points slightly earlier than the ripple peaks. Same conditions and statistical procedure as in A. White lines delineate significant clusters based on cluster-based permutation tests, whose *P*-values are stated in the upper left corners of the coactivity maps.

**Fig. S15.**
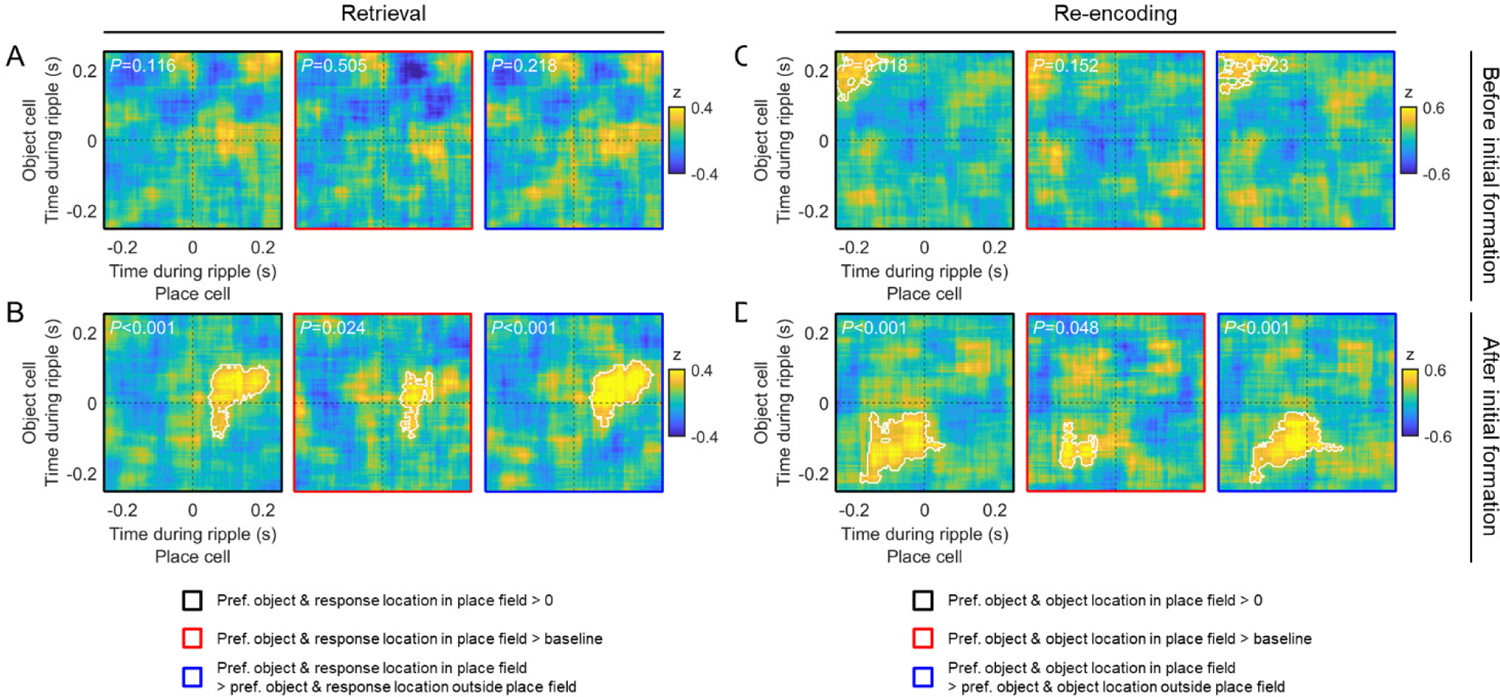
Coactivity of object cells and place cells during hippocampal ripples: exploratory analysis of the effect of the initial formation of the associative memories. (**A–D**) Same as Fig. 4, D, E, G, and H, but differentiating between ripples occurring “before initial formation” and ripples occurring “after initial formation” of the associative memories (instead of differentiating between early and late ripples). “Before initial formation” and “after initial formation” are defined by trial *i* such that the two-sample *t*-test between the memory performance values from trials *i*+1 to *n* and those from trials 1 to *i* leads to the highest *t*-statistic (where *n* is the maximum number of trials). Note though that learning still continued in “after initial formation” trials (Fig. 1D). (**A**) Coactivity maps estimated using ripples from the retrieval periods occurring “before initial formation,” considering only trials in which the subject was asked to remember the location of the preferred object of the object cell and in which the subject’s response location was inside the place field of the place cell. Left panel, comparison of the coactivity *z*-score maps against chance (i.e., 0). Middle panel, comparison of the coactivity *z*-score maps against baseline coactivity *z*-score maps. Right panel, comparison of the coactivity *z*-score maps against coactivity *z*-score maps estimated using ripples from trials in which the subject was asked to remember the location of the preferred object of the object cell and in which the subject’s response location was outside the place field of the place cell. (**B**) Same as in A, but only considering retrieval-related hippocampal ripples occurring “after initial formation.” (**C**) Coactivity maps estimated using ripples from the re-encoding periods occurring “before initial formation,” considering only trials in which the subject was asked to re-encode the correct location of the preferred object of the object cell and in which the object’s correct location was inside the place field of the place cell. Left panel, comparison of the coactivity *z*-score maps against chance (i.e., 0). Middle panel, comparison of the coactivity *z*-score maps against baseline coactivity *z*-score maps. Right panel, comparison of the coactivity *z*-score maps against coactivity *z*-score maps estimated using ripples from trials in which the subject was asked to re-encode the location of the preferred object of the object cell and in which the object’s correct location was outside the place field of the place cell. (**D**) Same as in C, but only considering re-encoding-related hippocampal ripples occurring “after initial formation.” White lines delineate significant clusters based on cluster-based permutation tests, whose *P*-values are stated in the upper left corners of the coactivity maps. >, larger than; pref., preferred.

**Fig. S16.**
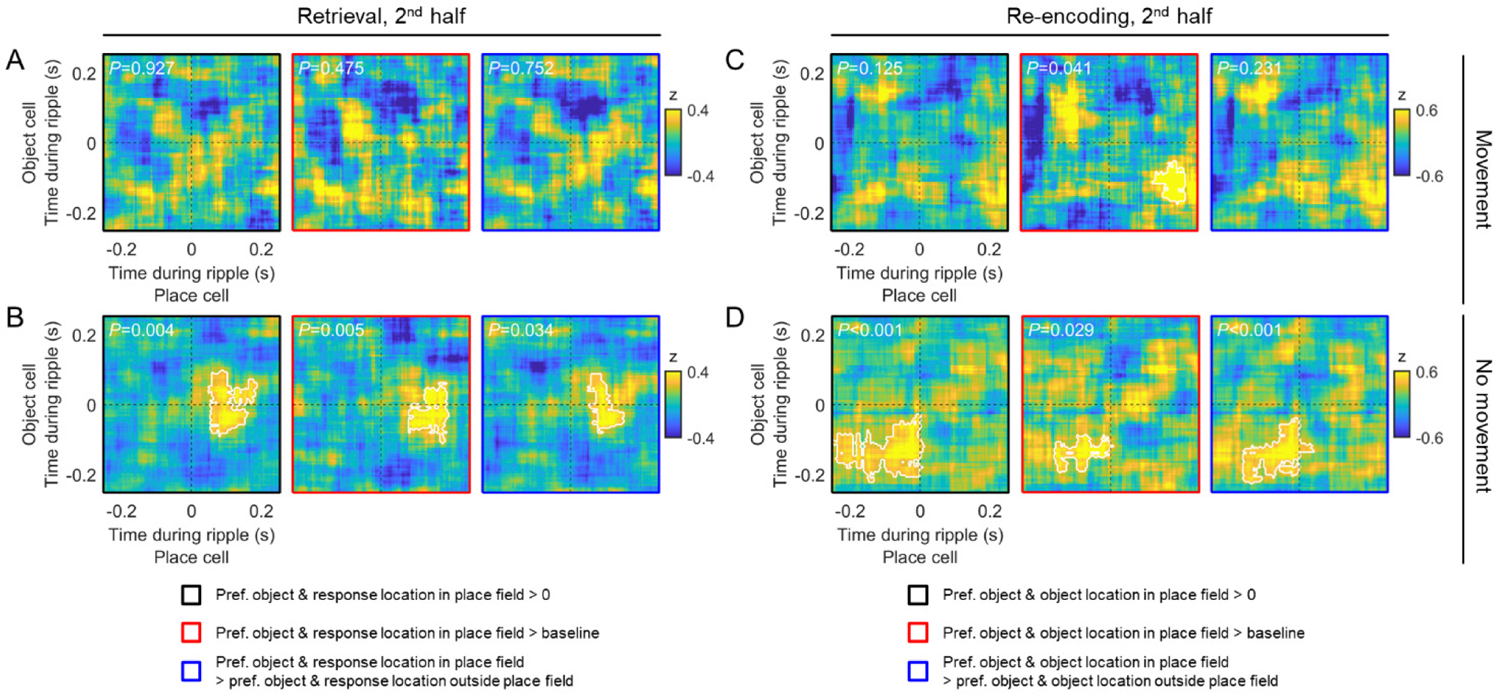
Coactivity of object cells and place cells during hippocampal ripples: exploratory analysis of the effect of (non)movement. (A–D) Same as Fig. 4, E and H, but only considering ripples during movement or non-movement periods. This shows that the coactivity effects were mainly driven by ripples occurring during non-movement periods. (A) Coactivity of object cells and place cells during hippocampal ripples from the second half of the retrieval periods, only considering time periods when the subject was moving. (B) Coactivity of object cells and place cells during hippocampal ripples from the second half of the retrieval periods, only considering time periods when the subject was not moving. (C) Coactivity of object cells and place cells during hippocampal ripples from the second half of the re-encoding periods, only considering time periods when the subject was moving. (D) Coactivity of object cells and place cells during hippocampal ripples from the second half of the re-encoding periods, only considering time periods when the subject was not moving. White lines delineate significant clusters based on cluster-based permutation tests, whose *P*-values are stated in the upper left corners of the coactivity maps. >, larger than; pref., preferred.

